# Regulation of oxidative phosphorylation by Nuclear myosin 1 protects cells from metabolic reprogramming and tumorigenesis in mice

**DOI:** 10.1101/2022.06.30.498328

**Authors:** Tomas Venit, Oscar Sapkota, Wael Said Abdrabou, Palanikumar Loganathan, Renu Pasricha, Syed Raza Mahmood, Nadine Hosny El Said, Sneha Thomas, Youssef Idaghdour, Mazin Magzoub, Piergiorgio Percipalle

## Abstract

Metabolic reprogramming is one of the hallmarks of tumorigenesis. Using a combination of multi-omics, here we show that nuclear myosin 1 (NM1) serves as a key regulator of cellular metabolism. As part of the nutrient-sensing PI3K/Akt/mTOR pathway, NM1 forms a positive feedback loop with mTOR and directly affects mitochondrial oxidative phosphorylation (OXPHOS) via transcriptional regulation of mitochondrial transcription factors TFAM and PGC1α. NM1 depletion leads to suppression of PI3K/Akt/mTOR pathway, underdevelopment of mitochondria inner cristae, and redistribution of mitochondria within the cell, which is associated with reduced expression of OXPHOS genes, decreased mitochondrial DNA copy number and deregulated mitochondrial dynamics. This leads to metabolic reprogramming of NM1 KO cells from OXPHOS to aerobic glycolysis and with a metabolomic profile typical for cancer cells, namely, increased amino acid-, fatty acid-, and sugar metabolism, and increased glucose uptake, lactate production, and intracellular acidity. We show that NM1 KO cells form solid tumors in a nude mouse model even though they have suppressed the PI3K/Akt/mTOR signaling pathway suggesting that the metabolic switch towards aerobic glycolysis provides a sufficient signal for carcinogenesis. We suggest that NM1 plays a key role as a tumor suppressor and that NM1 depletion may contribute to the Warburg effect at the early onset of tumorigenesis.

## Introduction

Functional mitochondria are crucial for a healthy cell as they maintain intracellular calcium levels, communicate with the nucleus via metabolites produced from the Krebs cycle to initiate epigenetic changes, and modulate their dynamics to fit the bio-energetic demands of cells (Salminen et al., 2015; Salminen et al., 2014; Sironi et al., 2020; Vandecasteele et al., 2001). However, their primary role is to produce energy in the form of up to 36 ATP molecules via OXPHOS In hypoxic conditions, when oxygen levels are limited, cells switch to the less efficient glycolysis pathway which converts glucose to lactate producing only 2 molecules of ATP per molecule of glucose. As the majority of cells use OXPHOS as a primary energy source, the expression of both nuclear and mitochondrial genes encoding macromolecular complexes involved in the OXPHOS electron transport chain is tightly regulated (Sunnucks et al., 2017). This is not true for highly proliferating undifferentiated pluripotent and cancer stem cells which use glycolysis as a primary source of energy production even in the presence of oxygen. The so-called aerobic glycolysis or Warburg effect was initially explained as a consequence of dysfunctional mitochondria in cancer cells. Nowadays it is highly accepted that aerobic glycolysis in these cells does not serve as a rescue mechanism for defective mitochondria but it is a rather universal, highly regulated metabolic pathway, which, even though less energetically effective, provides advantages to the cells. Glycolysis produces ATP faster than OXPHOS, as a process, it is less dependent on environmental factors, it regulates the tumor microenvironment by increasing intra- and extracellular acidity, and it allows signal transduction through different second messengers and promotes flux of byproducts into biosynthetic pathways (Liberti and Locasale, 2016). Metabolic switches characterized by increased mitochondrial OXPHOS and decreased glycolysis to produce ATP are key features that mark the differentiation of progenitor cells to committed cell lineages (Folmes et al., 2011). Similarly, metabolic switches are observed in tumorigenesis, where the prevalence of glycolytic metabolism over OXPHOS is connected to poor survival prognosis in many different types of cancers (Yu et al., 2019). Mitochondrial metabolism and more specifically, the relationship between OXPHOS and aerobic glycolysis plays, therefore, an important role in key cellular processes such as stemness and differentiation, but is also a defining feature during carcinogenesis. While several cytoskeletal proteins such β actin, Myosin II, or Myosin XIX have been shown to regulate mitochondrial dynamics directly (Korobova et al., 2014; Majstrowicz et al., 2021; Sato et al., 2022; Yang and Svitkina, 2019) and some of them are even present within the mitochondria (Shi et al., 2022; Xie et al., 2018), changes from OXPHOS to glycolysis are likely to require complex gene expression regulation yet to be fully understood.

Nuclear myosin 1 (NM1), a Myosin 1C isoform, facilitates transcription activation as part of the chromatin remodeling complex B-WICH (Percipalle et al., 2006; Pestic-Dragovich et al., 2000; Philimonenko et al., 2004; Sarshad et al., 2013; Ye et al., 2008). NM1 interacts with the remodelers’ ATPase SNF2h, enabling nucleosome repositioning and NM1-dependent recruitment of histone acetyl-transferase (HAT) PCAF and histone methyl-transferase (HMT) Set1B, which acetylate and methylate histone H3 to craft a chromatin landscape favorable for transcription activation (Almuzzaini et al., 2015; Sarshad et al., 2013; Vintermist et al., 2011). Recent work indicated a global role for NM1 in transcriptional regulation (Venit et al., 2020a; Venit et al., 2020b). Several studies have even proposed that the Myo1C gene could be a possible tumor suppressor as it is often mutated in various types of cancers (Hedberg Oldfors et al., 2015; Visuttijai et al., 2016) and we have recently shown that deletion of Nuclear myosin 1 leads to genome instability, elevated cell proliferation and increased DNA damage (Venit et al., 2020b). As these are all typical hallmarks of cancer cells, we examined whether dysregulated transcription upon NM1 depletion correlates with changes in energy metabolism these cells use to tackle their energetic needs.

Here we show evidence that metabolic reprograming is transcriptionally regulated by NM1 and this has a direct effect on mitochondrial biogenesis and function. Indeed, NM1 loss leads to a metabolic switch from OXPHOS to aerobic glycolysis and therewith associated mitochondrial phenotypes such as reduction of mitochondrial networks, underdeveloped inner membrane cristae, and transcriptomic changes in mitochondrial biogenesis. NM1 knockout (KO) cells exhibited transcriptional and metabolome profiles typical of cancer cells and were found to form solid tumors in immunosuppressed mice. Mechanistically, we found that NM1 activates the expression of specific mitochondrial transcription factors working as effectors of the PI3K/AKT/mTOR axis and NM1 is part of a positive feedback loop with mTOR. We suggest a new pathway in which, upon stimuli, mTORC1 phosphorylates NM1 which then binds to the transcription start site (TSS) of mitochondrial transcription factors PGC1α and TFAM as well as to the mTOR itself. This leads to an accumulation of mitochondrial factors, increased mitochondrial biogenesis, and expression of additional mTOR protein which can further activate NM1 until stimuli persist. We speculate that these novel mechanisms are important during tumorigenesis and that a positive feedback loop between mTOR and NM1 can play a role in cancer cell survival during mTOR-targeted cancer therapies.

## Results

### NM1 deletion suppresses the expression of OXPHOS genes and dysregulates the expression of mitochondrial genes

Seeing the role of NM1 in the transcriptional response to DNA damage (Venit et al., 2020b), we specifically investigated the potential effect of NM1 deletion on the expression of mitochondrial genes. Total protein extracts from stable NM1 WT and KO mouse embryonic fibroblasts were subjected to immunoblotting with the Total OXPHOS Rodent WB Antibody Cocktail kit containing antibodies against subunits of the five complexes (complexes I – V) in the OXPHOS system - Nduf88, Sdhb, Ucqrc2, MtCo1, and ATP5a (Figure 1A). Densitometric quantification of the bands from four independent immunoblots shows a significant decrease in the amount of each protein across all OXPHOS complexes in KO cells compared to WT condition, suggesting a general decrease in OXPHOS protein expression level in the absence of NM1 (Figure 1B). RTqPCR analysis of nuclear-encoded OXPHOS genes, representing all five subunits of the OXPHOS chain (Ndufs1, SDHA, UQCRB, Cox5A, and ATPf51) and mitochondrial-encoded OXPHOS genes (mt-CO1, mt-Cyb, and mt-ND1) confirmed the results from immunoblots (Figure 1C and 1D) (Supplementary table 1). In addition, analysis of RNA sequencing data from primary mouse embryonic fibroblasts (Venit et al., 2020b) derived from NM1 WT and KO embryos (Venit et al., 2013) corroborated the above findings and further demonstrated that the majority of differentially expressed OXPHOS genes are downregulated in NM1 KO cells (Figure 1E) supporting previous results observed in stable NM1 KO MEFs. The comparison of all differentially expressed genes with all genes under the MGI Gene Ontology (GO) term “Mitochondrion” (GO:0005739) revealed a high correlation between the groups with over 40% of all mitochondria-associated genes being differentially expressed in the NM1 KO condition (Figure 1F) affecting processes such as mitochondrial organization, translation, and transport together with previously described OXPHOS (Figure 1G). Overall, these results indicate a wide transcriptional change in mitochondria-related gene expression accompanied by downregulation of OXPHOS genes in NM1 deficient cells.

**Figure 1.**
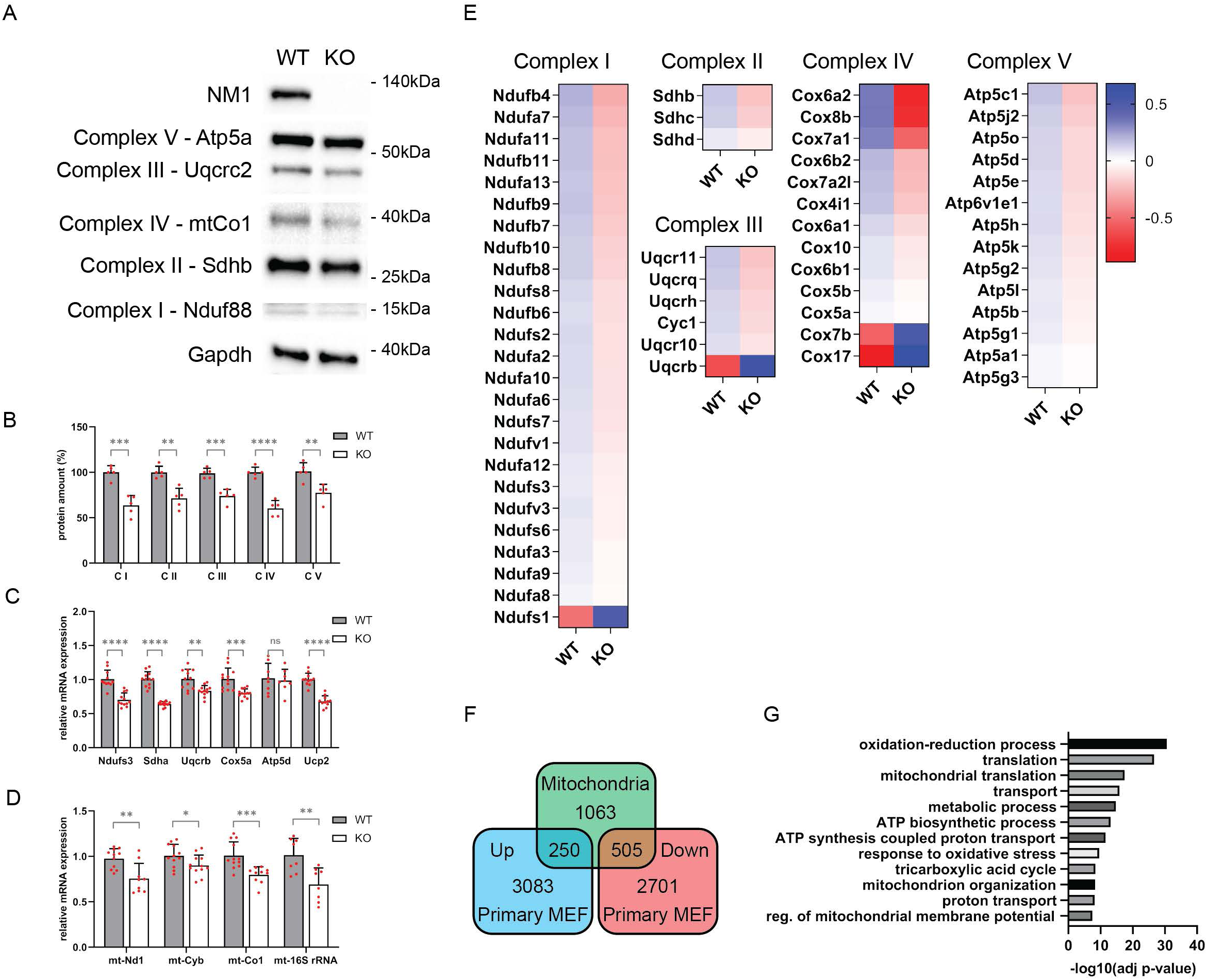
NM1 deletion suppresses the expression of OXPHOS genes and leads to dysregulation of mitochondrial gene expression. (A) Representative western blots from WT and NM1 KO cells stained with antibodies against proteins in the OXPHOS electron transfer chain, NM1, and control GAPDH. (B) Quantification of protein levels in immunoblots from (A) in WT and NM1 KO MEFs, normalized against GAPDH levels, *n*≥4. (C) RT-qPCR analysis of nuclear-encoded mitochondrial OXPHOS genes in WT and NM1 KO MEFs. Gene expression levels are relative to Nono mRNA levels. *n*≥8, \*\**p* < 0.01, \*\*\**p* < 0.001, \*\*\*\**p* < 0.0001. Bars represent mean with error bars representing SD. (D) RT-qPCR analysis of mitochondria-encoded genes in WT and NM1 KO MEFs. Expression levels are relative to Nono mRNA. *n*≥8, \*\**p* < 0.01, \*\*\**p* < 0.001, \*\*\*\**p* < 0.0001, ns (not significant). Bars represent mean SEM. (E) Heatmap of differentially expressed nuclear OXPHOS genes in primary WT or NM1 KO MEFs. Heatmap represents the mean log2-normalized counts for 3 WT and 3 KO samples in descending order. (F) Venn diagram shows the intersection between genes associated with the GO term “Mitochondrion,” and all up- and down- regulated differentially expressed genes in primary WT and NM1 KO MEFs. (G) Differentially expressed genes associated with the GO term “Mitochondrion” were subjected to GO analysis, the top 12 enriched biological processes are shown in descending order.

### NM1 depletion leads to reduced mitochondrial mass, perinuclear localization and altered mitochondrial morphology and structure

To address the effect of NM1 deletion on mitochondrial function and structure, we first analyzed mitochondrial mass and cellular distribution by high-content phenotypic profiling of MitoTracker® DeepRed (MitoTracker DR) stained mitochondria. We used Compartment Analysis BioApplication software to generate masks for measuring quantitative parameters. The mask covering the whole cell except the nuclear region was used for quantification of mitochondrial mass (Figure 2A) and the mask distinguishing between cytoplasmic and perinuclear regions was used to define mitochondrial localization (Figure 2B). Results from this analysis showed that NM1 KO cells display decreased average intensity and area of Mitotracker DR staining (Figure 2A) with increased perinuclear/cytoplasmic ratio of the same parameters in comparison to WT cells (Figure 2B). This suggests that NM1 depletion leads to reduced mitochondrial mass and predominant perinuclear localization of mitochondria. To support these findings and bypass possible microscopy artifacts in high-content screening, we also quantified the mitochondrial mass by spectrophotometric analysis of MitoTracker DR staining normalized to nuclear Hoechst staining. Similar to previous results, NM1 KO cells showed decreased mitochondrial staining in comparison to WT cells (Figure 2C). Importantly, the staining of mitochondria with mitochondrial membrane potential-dependent MitoTracker® Orange dye showed a proportional decrease in NM1 KO cells comparable to the decrease of total mitochondrial mass (Figure 2D).

**Figure 2.**
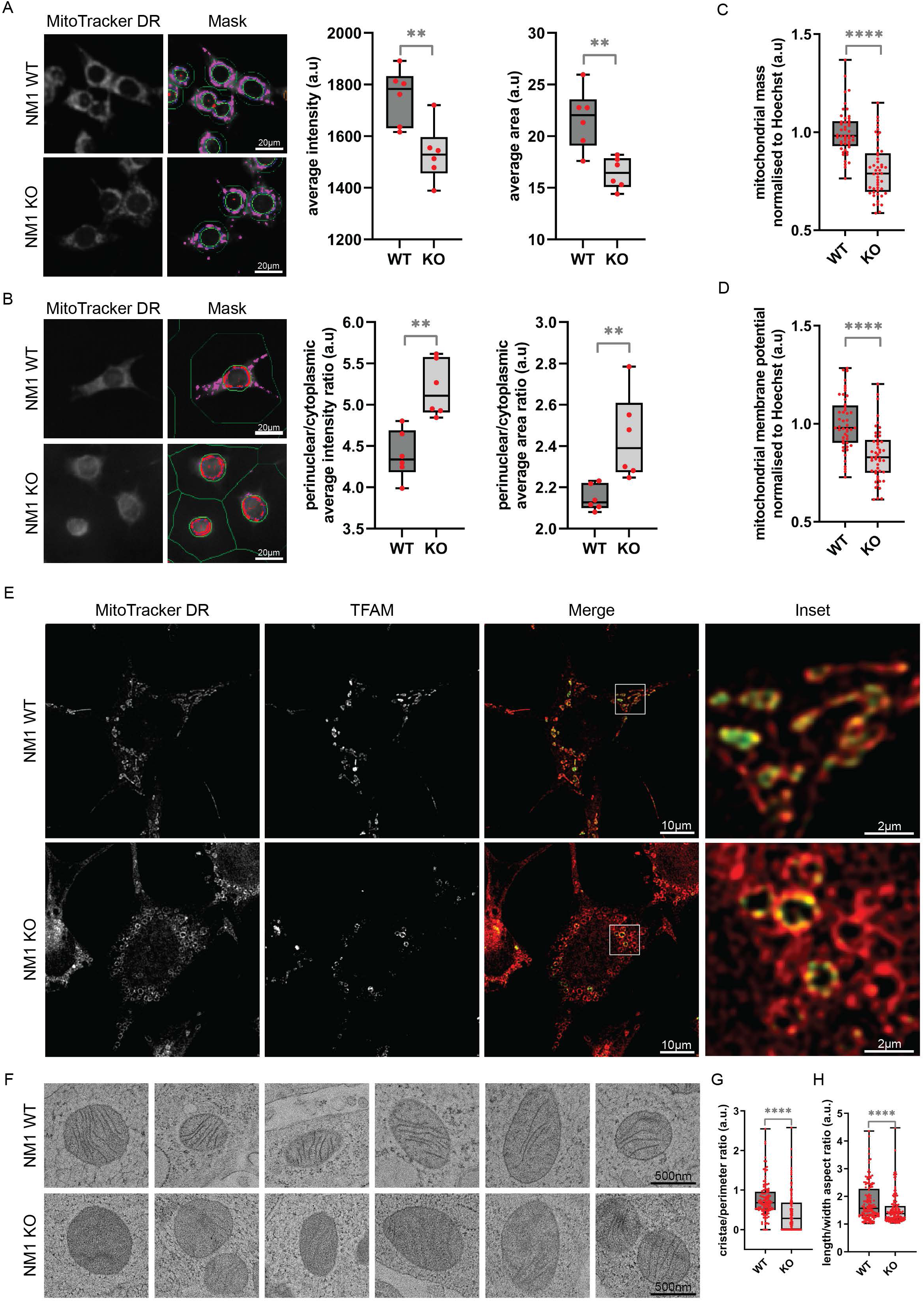
NM1 depletion leads to reduced mitochondrial mass, perinuclear localization, and altered mitochondrial morphology and structure. (A) High content phenotypic profiling of WT and NM1 KO MEFs. Representative pictures show MitoTracker Deep Red (DR) staining and respective masks used for quantification. The average intensity and average area of MitoTracker DR-stained mitochondria are plotted. Each box plot represents the mean value and first and third quartile values. Error bars represent minimum and maximum values. Each dot represents a mean value for one measurement. For each measurement, at least 1000 cells have been quantified. *n* = 6. \*\**p* < 0.01. (B) High content phenotypic profiling of WT and NM1 KO MEFs. Representative pictures show MitoTracker DR staining and respective masks used for differentiating and quantification of perinuclear and cytoplasmic mitochondrial staining. The ratio between perinuclear and cytoplasmic mitochondrial intensity and area of MitoTracker DR stained mitochondria are plotted. Each box plot represents the mean value and first and third quartile values. Error bars represent minimum and maximum values. Each dot represents a mean value for one measurement. For each measurement, at least 1000 cells have been quantified. *n* = 6. \*\**p* < 0.01. (C) Spectrophotometric analysis of mitochondrial mass in WT and NM1 KO MEFs. The graph represents the MitoTracker DR fluorescence signal normalized to Hoechst staining in each condition. Each box plot represents the mean value and first and third quartile values. Error bars represent minimum and maximum values. The visualized data represent the compilation of 3 independent experiments with 16 wells separately measured for each condition in each experiment. *n* = 48. \*\*\*\**p* < 0.0001. (D) Spectrophotometric analysis of mitochondrial membrane potential in NM1 WT and KO cells. The graph represents the MitoTracker Orange fluorescence signal normalized to Hoechst staining in both conditions. Each box plot represents the mean value and first and third quartile values. Error bars represent minimum and maximum values. The visualized data represent the compilation of 3 independent experiments with 16 wells separately measured for each condition in each experiment. *n* = 48. \*\*\*\**p* < 0.0001. (E) Representative confocal microscopy images of MitoTracker DR and TFAM stained mitochondria in WT and NM1 KO MEFs. (F) Representative electron microscopy images of mitochondria in WT and NM1 KO MEFs. (G) Quantification of mitochondrial cristae length to mitochondrial perimeter ratio in both conditions. Each box plot represents the mean value and first and third quartile values. Error bars represent minimum and maximum values. Each dot represents a single mitochondria measurement. *n* = 150. \*\*\*\**p* < 0.0001. (H) Quantification of mitochondrial circularity is defined as length to width ratio. Each box plot represents the mean value and first and third quartile values. Error bars represent minimum and maximum values. Each dot represents a single mitochondria measurement. *n* = 150. \*\*\*\**p* < 0.0001

To see if decreased mitochondrial mass and membrane potential in NM1 KO are accompanied by changes in mitochondrial morphology, we first visualized WT and NM1 KO MEFs by confocal microscopy (Figure 2E). Cells were stained with MitoTracker DR and antibody against mitochondrial transcription factor A (TFAM), which is the major regulator of mitochondrial transcription and mitochondrial genome replication (Picca and Lezza, 2015; Pohjoismaki et al., 2006). We found that WT cells exhibited a network of elongated and circular mitochondria decorated with relatively constant levels of TFAM staining, typical for differentiated cells (Noguchi and Kasahara, 2018). In contrast, NM1 KO cells displayed predominantly circular mitochondria with limited mitochondrial network formation and heavily reduced TFAM staining, typical for undifferentiated stem cells and cancer cells (Lonergan et al., 2006; Vander Heiden et al., 2009). To study the mitochondrial structure in more detail, we applied transmission electron microscopy (TEM) on both WT and KO NM1 cells stained with osmium as a contrasting agent to label cellular lipids and mitochondria with clearly visible outer membrane (Figure 2F). Quantification of mitochondrial inner membrane cristae and mitochondrial shape in both cell types showed that NM1 KO mitochondria have underdeveloped or missing cristae and more dispersed osmium staining in comparison to the WT condition (Figure 2F) as revealed by measuring the ratio between cristae length and mitochondrial perimeter (Figure 2G). Similarly, measurement of mitochondrial length to width ratio supports previous findings from confocal microscopy showing higher circularity of mitochondria in NM1 KO cells (Figure 2H). Taken together these results suggest that NM1 depletion leads to altered mitochondrial morphology and distribution within cells.

### NM1 depletion results in reduced mitochondrial copy number and dysregulated mitochondrial dynamics

In the context of mitochondrial biogenesis, the mitochondrial transcription regulator TFAM is also involved in mitochondrial genome replication and mtDNA copy number (Ekstrand et al., 2004). Seeing the reduction of TFAM staining in KO cells, we investigated differences in mitochondrial DNA copy number by quantitative-PCR (qPCR) of genomic DNA (nuclear and mitochondrial) isolated from NM1 WT and KO MEFs. For this, we compared the relative abundances of three mitochondrial genes mtCyb, mtNd1, and mtATP6 normalized to the nuclear-encoded Terf gene (Xie et al., 2018) (Supplementary Table 1). In agreement with previous studies and our results, the lack of NM1 and subsequent decrease in TFAM level were found to correlate with a reduction of mitochondrial copy number (Figure 3A). To further explore aberrant mitochondrial distribution, structure, and function, we studied gene expression levels of key proteins involved in mitochondrial dynamics. For this purpose, we performed Rt-qPCR analysis on gene markers for mitochondrial fission (DNM1L and Fis1), mitochondrial fusion (Mfn1 and Opa1), and quality control (Pink1 and Snca) and mitophagy (Becn and Sqstm) (Supplementary Table 1). While mitochondrial fission was not affected by NM1 deletion as revealed by the unaltered levels of DNM1L and Fis1 gene expression, all other steps in mitochondrial turnover were found to be dysregulated, including mitochondrial fusion (increased Mfn1 levels), quality control (decreased expression of both Pink1 and alpha-Synuclein) and mitophagy (decreased Becn levels) (Figure 3B). These data amend previous findings and suggest that reduced mitochondrial mass and membrane potential could be caused by decreased levels of mitochondrial biogenesis, while alterations in mitochondrial structure and localization could be due to aberrant mitochondrial dynamics and quality control signaling.

**Figure 3.**
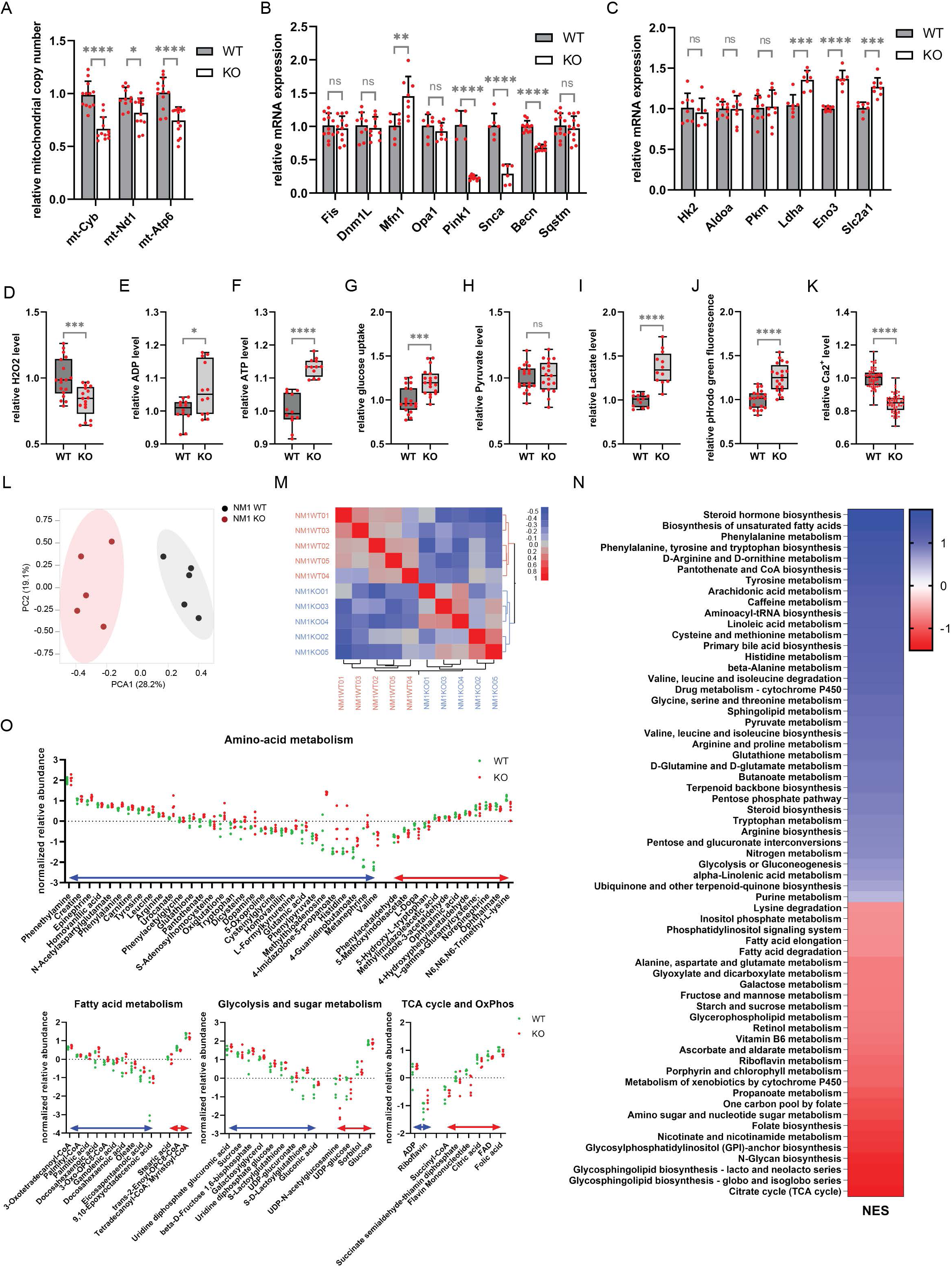
NM1 depletion results in reduced mitochondrial copy number and dysregulated mitochondrial dynamics - NM1 KO cells switch their metabolism from OXPHOS to Glycolysis. (A) qPCR analysis of mitochondrial DNA copy number as determined by the expression of mitochondrial genes mtCyb, mtNd1, and mtATP6 normalized to reads from the nuclear-encoded Terf gene in WT and NM1 KO MEFs. Each dot represents a single measurement. Bars represent mean with SD. *n*≥12, \**p* < 0.05, \*\*\*\**p* < 0.0001. (B) RT-qPCR analysis of key genes regulating mitochondrial dynamics in WT and NM1 KO cells. The expression of each gene is measured relative to the expression of the Nono gene. Each dot represents a single measurement. Bars represent mean with SD. *n*≥5, \*\**p* < 0.01, \*\*\*\**p* < 0.0001, ns (not significant). (C) RT-qPCR analysis of glycolytic genes in WT and NM1 KO cells. The expression of each gene is relative to the expression of the Nono gene. Each dot represents a single measurement. Bars represent mean with SD. *n*≥7, \*\*\**p* < 0.001, \*\*\*\**p* < 0.0001. ns (not significant). (D - K) Relative fluorometric quantification of different metabolites in WT and NM1 KO cells. Each box plot represents the mean value and first and third quartile values. Error bars represent minimum and maximum values. Each dot represents a single measurement. *n*≥12. \**p* < 0.05, \*\*\**p* < 0.001, \*\*\*\**p* < 0.0001, ns (not significant). (L) PCA was performed on the normalized peak dataset from WT and NM1 KO cells. (M) Global metabolomic correlation matrix clustering based on similarity from WT and NM1 KO cells. (N) GSEA analysis of compounds identified using positive ionization organized descending based on their normalized enrichment scores (NES). Blue color represents pathways that are overrepresented and red color pathways that are underrepresented in NM1 KO cells. (O) Graphs represent the normalized relative abundance of each detected compound in the given metabolic pathway. The blue double arrow represents metabolites with relative abundance higher in KO, while the red double arrow represents metabolites with higher abundance in WT. Each dot represents one measurement of a given compound.

### NM1 KO cells switch their metabolism from OXPHOS to Glycolysis

To test whether NM1 KO cells switch from OXPHOS to glycolysis, we first measured the expression of several glycolytic proteins. While some targets did not show altered expression (Hk2, Aldoa, Pkm), glucose transporter Glut1 (Slc2a1), enolase (Eno3), and lactate dehydrogenase A (Ldha), a critical enzyme in pyruvate to lactate conversion, were found to be upregulated in NM1 KO cells (Figure 3C). We next performed fluorometric-based assays to reveal potential differences in mitochondria-related metabolites. Mitochondrial reactive species such as hydrogen peroxide or superoxide (H2O2) are the main by-products during a series of electron flow processes through an electron transport chain (Murphy, 2009). When measuring the level of H2O2 we discovered a significant decrease in NM1 KO cells (Figure 3D) which correlates with a reduction of OXPHOS seen in the transcriptomic analysis. In contrast, ADP and ATP levels were significantly upregulated (Figure 3E and 3F), which seems contradictory to less energetically profitable glycolysis (Mookerjee et al., 2017; Zheng, 2012) but can be explained by the concomitant increased glucose consumption in the NM1 KO cells measured by fluorescently-labeled deoxyglucose analog 2-NDBG uptake (Figure 3G). In the final step of glycolysis, pyruvate can be either metabolized in the TCA cycle in mitochondria or can be fermented to lactate and/or lactic acid in glycolysis. While cellular pyruvate level was found to be constant between WT and KO condition (Figure 3H), lactate levels were significantly increased in NM1 KO cells (Figure 3I). Consistently, measurement of intracellular pH by pHrodo green dye showed higher intracellular acidity in the absence of NM1 (Figure 3J). Apart from the production of ATP and other metabolites, mitochondria serve as the main regulator of calcium homeostasis in cells through their interactions with other organelles and by their calcium buffering capacity (Deak et al., 2014). Therefore, we measured the total amount of intracellular Ca2+ in both cell types and found calcium levels to be significantly decreased in NM1 KO cells (Figure 3K).

Taken together, quantification of mitochondria-related metabolites as well as RTqPCR of glycolytic genes suggest that mitochondrial ATP production and mitochondrial cell signaling are impaired in the NM1 KO condition and upon NM1 deletion cells seem to switch to a glycolytic metabolism even in the presence of oxygen.

### NM1 KO cells exhibit a metabolome profile typical for cancer cells

As the NM1 KO cells seem to move from OXPHOS to aerobic glycolysis (Liberti and Locasale, 2016), we next explored if NM1 depletion leads to metabolic reprogramming. We performed Liquid Chromatography followed by High-Resolution Mass Spectrometry of cellular extracts from NM1 WT and KO cells, and quality control of the peak data resulted in the retention of 7,212 from a total of 15,423 compounds for downstream analysis; 4,378 compounds were identified using positive ionization and 2,839 compounds identified using negative ionization. Principal-component analysis (PCA) of the normalized peak dataset revealed a strong correlation structure in the metabolomic data across both conditions with the first principal component (PC1) capturing the effect of loss of NM1 and showing clear segregation between replicates of the WT and KO. The first two PCs explain 47.3% of the variation in the dataset (Figure 3L). Clustering of the global metabolomic correlation matrix based on similarity also clearly shows that replicates of each condition cluster together and NM1 deletion leads to distinct metabolomic changes in these cells (Figure 3M). To identify perturbations in mouse-specific metabolic pathways or metabolite sets in association with the loss of NM1, functional analysis using Gene Set Enrichment Analysis (GSEA) approach was performed for compounds detected using positive (Figure 3N) and negative (Supplementary figure 1) ionization separately and mapped onto Mus musculus (mouse) [KEGG]. Among others, the analysis revealed glycolysis, sugar metabolism, amino acid, and fatty acid metabolism pathways to be enriched in the NM1 KO Cells with increased levels of the majority of identified compounds while the TCA cycle and OXPHOS pathways were suppressed in these cells (Figure 3O). This correlates well with our previous findings and suggests that NM1 KO cells not only switch to aerobic glycolysis but fully reprogram to cancerous metabolism as several recent studies show similar metabolomic profiles in different cancer types (Han et al., 2021). For example, deregulated fatty acid and glucose metabolism were found in lung cancer patients (Callejon-Leblic et al., 2019); nucleotide, histidine, and tryptophan metabolism in ovarian cancer (Zhang et al., 2013); and most prominently, purine metabolism, glycine, serine, arginine and proline metabolism, steroid biosynthesis, sphingolipid metabolism and bile metabolism deregulated in pancreatic cancer (Luo et al., 2020). All aforementioned metabolic pathways were found to be altered in NM1 KO cells as well. A similar analysis was performed where annotated compounds were mapped onto a curated 912 metabolic data sets predicted to change due to dysfunctional enzymes based on human metabolism. The analysis revealed the enrichment of similar pathways as found for the mouse KEGG database (Supplementary table 2)

In conclusion, global changes in the metabolome of NM1 KO cells associated with increased amino-acid and fatty acid metabolism support the idea of using glycolysis byproducts for rapid biosynthesis of biologically relevant molecules needed for increased proliferation. Herewith associated phenotypes, namely decreased production of ROS, imbalance in calcium homeostasis, increased glucose uptake, increased lactate level coupled with increased intracellular acidity, and upregulated expression of glycolytic enzymes, do not only describe characteristics of the glycolytic metabolism but are also the defining hallmarks of cancerous cells and tumors.

### NM1 regulates mitochondria by regulating specific mitochondrial transcription factors

To address if NM1 binds to mitochondrial gene promoters, we performed chromatin immunoprecipitation followed by deep sequencing (ChIP-Seq) with antibodies to SNF2H and active histone marks H3K9Ac and H3K4me and examined the chromatin accessibility by assay for transposase-accessible chromatin with sequencing (ATAC-Seq) in NM1 WT and KO cells. Remarkably, NM1 deletion did not show significant changes in SNF2H binding or distribution of active histone marks and did not lead to a change in chromatin accessibility around the transcription start sites of mitochondrial genes (Supplementary figure 2A and 2B). We next tested whether changes in gene expression might be a consequence of changes in genome organization. The nuclear genome is hierarchically organized into A and B compartments which respectively correlate with open and closed chromatin or euchromatin and heterochromatin (Fortin and Hansen, 2015). We recently reported that loss of nuclear actin leads to compartment switching, in turn leading to differential expression of genes present within these compartments (Mahmood et al., 2021). Mitochondrial genes tend to cluster together and are often found in the pericentromeric and subtelomeric regions, which are characterized by lesser chromatin accessibility and heterochromatinization (Moon et al., 2008). As around 80 percent of NM1 DNA binding occurs outside of transcription start sites and gene promoters (Almuzzaini et al., 2015), we next tested whether NM1 could regulate mitochondrial gene expression by mediating the 3D spatial organization of the genome. We, therefore, performed Hi-C chromosome conformation capture analysis combined with deep sequencing (Hi-C-seq) of NM1 WT and KO cells to study compartment changes and their potential effect on mitochondrial gene expression. However, we observed a low level of compartment switching in the NM1 KO condition (Supplementary figure 2C), and only a very small portion of mitochondrial genes is associated with these regions (Supplementary figure 2D).

As we were unable to associate NM1 with the expression of mitochondrial genes directly or via reorganization of 3D genome structure, we examined whether NM1 could regulate the expression of specific mitochondrial transcription factors, namely Nrf1, Nrf2, PGC1α, TFAM, Tfb2m, Yy1, and Essra (Leigh-Brown et al., 2010), as upstream regulators of a broader range of mitochondrial genes. Quantification by RTqPCR revealed several of them to be differentially expressed upon NM1 deletion (Figure 4A). We further focused on two major factors, PGC1α and TFAM. PGC1α is a major mitochondrial co-activator that interacts with the majority of other mitochondrial factors and, as mentioned earlier, TFAM is responsible for transcription of mitochondria-encoded genes and mitochondrial DNA replication (Kang et al., 2018; Puigserver and Spiegelman, 2003). We hypothesized that if NM1 is responsible for the expression of OXPHOS genes via regulation of mitochondrial transcription factors, NM1 binding to the promoters of these genes should be reduced once cells switch to glycolysis. We, therefore, reduced the oxygen level in cells to induce hypoxia-driven glycolysis over OXPHOS (Figure 4B) and applied chromatin immunoprecipitation (ChIP) to measure NM1 occupancy at the transcription start sites of PGC1α and TFAM genes in cells under normal or hypoxic conditions. Results from ChIPqPCR revealed that NM1 binds to both TSSs under normoxic conditions, but its binding is heavily decreased upon hypoxia (Figure 4C). Similarly, deletion of NM1 leads to a significant decrease of active histone marks H3K9Ac and H3K4me over PGC1α and TFAM TSSs under normal conditions and is even more prominent upon hypoxia (Figure 4D). Interestingly, there is a positive correlation between NM1 binding and the association of active histone marks over the transcription start sites. NM1 binding is higher at the TFAM gene TSS in comparison to binding to PGC1α and it is reflected in the much stronger association of H3K9Ac and H3K9me3 with the TFAM gene.

**Figure 4.**
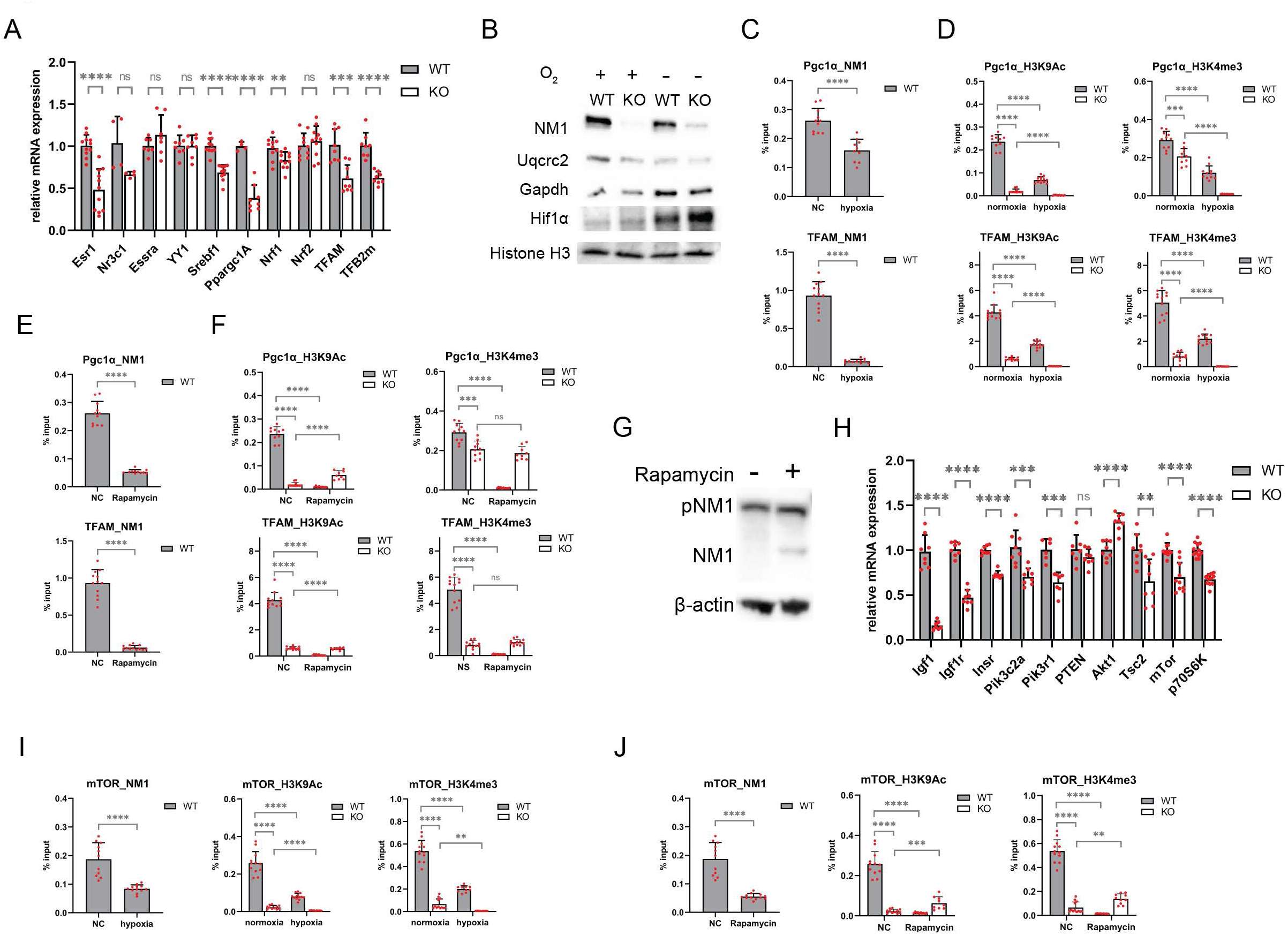
NM1 regulates mitochondrial function through the PI3K/AKT/mTOR pathway and forms a positive feedback loop with mTOR. (A) RT-qPCR analysis of mitochondrial transcription factors in WT and NM1 KO cells. The expression of each gene is measured relatively to Nono gene expression levels. Each dot represents a single measurement. Bars represent mean with SD. *n*≥7, \*\**p* < 0.01, \*\*\**p* < 0.001, \*\*\*\**p* < 0.0001, ns (not significant). (B) Western blotting analysis of cell lysates from WT and NM1 KO cells grown under normoxic conditions (O_2_ +) or hypoxic conditions (O_2_ -). Uqcrc2 is part of the OXPHOS pathway, Hif1α and GAPDH are glycolytic genes, and H3 Histone serves as a loading control. (C) ChIP-qPCR analysis of NM1 binding to transcription start sites of Pgc1α and TFAM normalized to input in normoxia and hypoxia. Each dot represents a single measurement. Bars represent mean with SD. *n*≥9, \*\*\*\**p* < 0.0001. (D) ChIP-qPCR analysis of H3K9Ac and H3K4me3 binding to transcription start sites of Pgc1α and TFAM normalized to input in normoxia and hypoxia. Each dot represents a single measurement. Bars represent mean with SD. *n*≥9, \*\*\**p* < 0.001, \*\*\*\**p* < 0.0001. (E) ChIP-qPCR analysis of NM1 binding to transcription start sites of Pgc1α and TFAM normalized to input upon Rapamycin treatment. Each dot represents a single measurement. Bars represent mean with SD. *n*≥9, \*\*\*\**p* < 0.0001. (F) ChIP-qPCR analysis of H3K9Ac and H3K4me3 binding to transcription start sites of Pgc1α and TFAM normalized to input upon Rapamycin treatment. Each dot represents a single measurement. Bars represent mean with SD. *n*≥9, \*\*\**p* < 0.001, \*\*\*\**p* < 0.0001, ns (not significant). (G) Phos-Tag western blot analysis of NM1 phosphorylation upon Rapamycin treatment. β-actin serves as a loading control. (H) RT-qPCR analysis of PI3K/Akt/mTOR signaling pathway genes in NM1 WT and KO cells. The expression of each gene is relative to Nono gene expression. Each dot represents a single measurement. Bars represent mean with SD. *n*≥8, \*\**p* < 0.01, \*\*\**p* < 0.001, \*\*\*\**p* < 0.0001. ns (not significant). (I) ChIP-qPCR analysis of NM1, H3K9Ac, and H3K4me3 binding to TSS of mTOR gene normalized to input in normoxia and hypoxia. Each dot represents a single measurement. Bars represent mean with SD. *n*≥7, \*\**p* < 0.01, \*\*\**p* < 0.001, \*\*\*\**p* < 0.0001, ns (not significant). (J) ChIP-qPCR analysis of NM1, H3K9Ac, and H3K4me3 binding to TSS of mTOR gene normalized to input upon Rapamycin treatment. Each dot represents a single measurement. Bars represent mean with SD. *n*≥7, \*\**p* < 0.01, \*\*\**p* < 0.001, \*\*\*\**p* < 0.0001, ns (not significant)

We conclude that NM1 deletion does not affect the expression of mitochondrial genes directly by changing the chromatin landscape or global 3D genome architecture, but rather through direct regulation of specific mitochondrial transcription factors.

### NM1 regulates mitochondrial function via the PI3K/AKT/mTOR pathway and forms a positive feedback loop with mTOR

Extracellular and intracellular nutrient sensing and the subsequent metabolic changes that occur are tightly regulated by several mechanisms and pathways. The mTOR kinase as a part of the mTOR complex 1 (mTORC1) controls cellular energetics by regulating transcription and translation of metabolic genes, inducing protein and lipid synthesis upon activation by growth factors through PI3K-AKT signaling (Efeyan et al., 2015). mTORC1 phosphorylates several downstream targets, many of which are mitochondrial transcription factors, which then regulate the expression of mitochondrial genes (de la Cruz López et al., 2019). Treating cells with specific mTORC1 inhibitor rapamycin leads to reduced expression of PGC1α and other mitochondrial transcription factors and decreased OXPHOS in these cells (Cunningham et al., 2007). As we have seen similar effects on protein expression and OXPHOS in the NM1 KO condition, we next tested whether NM1 could be regulated by the mTORC1 complex. We treated NM1 WT and KO cells with rapamycin and examined NM1 binding to TSSs of mitochondrial transcription factors PGC1α and TFAM by ChIPqPCR. In both cases, rapamycin treatment led to a significant reduction of NM1 occupancy over TSS of both genes followed by reduced distribution of active histone marks (Figure 4E and 4F). Interestingly, while in WT cells, rapamycin treatment led to an almost complete drop of active histone marks from TSS, NM1 KO cells seemed to be insensitive to rapamycin treatment, further suggesting the potential importance of an mTOR-NM1 regulatory cascade for cellular metabolism. Since NM1 is phosphorylated by GSK3β to stabilize chromatin association and protect it from proteasome degradation (Sarshad et al., 2014), we reasoned that mTOR may also be involved in regulating NM1 phosphorylation. Cell lysates prepared from DMSO and rapamycin-treated cells were run on Phos-Tag gels and stained with anti-NM1 antibody. Blocking mTORC1 specific phosphorylation by rapamycin led to a shift of the unphosphorylated NM1 band towards lower molecular weight, suggesting that NM1 is indeed a potential phosphorylation target for mTORC1 (Figure 4G) and might be involved in PI3K/AK/ mTOR signaling. We, therefore, measured the expression of the main proteins from growth factors to mTOR effector p70S6K and found the whole signaling cascade to be suppressed in the absence of NM1, including mTOR expression, with exception of Akt1 (Figure 4H). Based on these observations, NM1 could be part of a positive feedback loop with mTOR. To test this possibility, we next examined NM1 and active histone marks occupancies at TSS of the mTOR gene by ChIPqPCR under normoxic and hypoxic conditions as well as upon rapamycin treatment. Results from these experiments show that both hypoxia and rapamycin treatment lead to decreased NM1, H3K9Ac, and H3K4me enrichment around the mTOR TSS (Figure 4I and 4J).

Taken together, we suggest a model where NM1 functions as part of a nutrient-sensing signaling pathway. Extracellular and intracellular signals activate a cascade of phosphorylation events leading to activation of mTOR which subsequently activates NM1 and other factors. NM1 then stimulates the expression of mitochondrial transcription factors as well as mTOR itself which leads to multiplication and strengthening of signaling by newly produced mTOR until the extracellular signal persists. This would allow cells to keep relatively small levels of signaling proteins in a resting state and their robust accumulation upon external stimulation.

### NM1 is a novel tumor suppressor

We next investigated if deletion of NM1 is sufficient to form tumors in mice. We injected WT and NM1 KO cells under mammary pads of the Balb/c nude mouse model and studied tumor formation over a period of 3 weeks (Figure 5A). None of the mice with injected NM1 WT cells formed tumors while injection of NM1 KO cells led to 100% penetration and rapid tumor growth in all tested animals (Figure 5 B-D). Next, we prepared tissue sections from tumors isolated from mice injected with NM1 KO cells and tissue sections from mammary pads tissue from mice injected with WT cells. Immunohistochemistry analysis with antibodies against cancer markers EGFR, Ki-67, Bcl-XL, and Mct1 showed high positivity in tumor tissues while mammary pads of mice injected with NM1 WT cells did not show any staining (Figure 5E). Importantly, Mct1 is a monocarboxylate transporter responsible for the shuttling of lactate and pyruvate and serves as a prognostic marker for glycolytic tumors (Hong et al., 2016). Overexpression of Mct1 in tumors derived from NM1 KO cells therefore further supports our previous findings of the metabolic switch from OXPHOS to glycolysis type of metabolism. Hematoxylin and eosin staining of kidney, liver, lungs, spleen, mammary pads, and tumor tissues was used for histopathology. This revealed a very dense and uniform distribution of NM1 KO cells in tumor tissue which is very different from the mammary pads tissue injected with NM1 WT cells. The morphology of other selected tissues does not show any significant alterations upon injection of both cell types (Figure 5F). We conclude that NM1 deletion in cells is sufficient for carcinogenesis and formation of solid tumors in the place of injection but at least in this model system it does not seem to form metastases and secondary tumors in other organs within the duration of the experiment. To further understand the mechanisms behind the cellular transformation of NM1 KO MEFs to tumor tissue, we performed an RNA-seq analysis of the parental NM1 KO cell line and two tumors isolated from Balb/c nude. For comparative analysis, ee selected the fastest- and the slowest-growing tumors derived from NM1 KO MEFs, as WT MEFs did not lead to any tumor growth in Balb/c nude mice. Correlation matrix clustering between NM1 KO MEFs and both tumors shows high similarity in the transcriptional profiles with the fast-growing tumor being transcriptionally more similar to NM1 KO MEFs than the slow-growing tumor (Figure 6A). Gene expression comparison of the fast- or slow-growing tumor with NM1 KO MEFs, respectively, revealed a high correlation in differential gene expression in both tumor tissues (intersection) with some specific set of genes differentially expressed in each tumor (Figure 6B). We next performed GO analysis of these gene groups and plotted the most abundant biological process GO terms for common genes (Figure 6C), differentially expressed genes in fast-growing tumor (Figure 6D) or slow-growing tumor (Figure 6E). As predicted, tumorigenic transformation in both tumor types is associated with gene programs related to cell adhesion and migration, angiogenesis, inflammatory response, and aberrant cellular signaling (Figure 6C). The spectrum of differentially expressed genes specific for transformation to the fast-growing tumor is very broad and only related GO terms are associated with regulation of transcription and protein stabilization (Figure 6D). On the contrary, gene programs specifically affected in tumorigenic transformation in the slow-growing tumor are related not only to mitochondrial function but also to DNA damage signaling, DNA repair, and cell cycle regulation which are affected by NM1 (Venit et al., 2020b) (Figure 6E). To understand this transcriptomic profile in more detail, we compared gene expression patterns of two tumors and performed GO analysis of genes that are downregulated (Figure 6F) or upregulated (Figure 6G) in slow-growing tumor versus fast-growing tumor. We found that in slow-growing tumors, genes related to DNA damage and cell cycle are less expressed while genes associated with aerobic respiration are more expressed in comparison to fast-growing tumors (Figure 6F and 6G). We next plotted heatmaps with the normalized RNA-seq count values for each sample of all differentially expressed genes related to DNA damage signaling and repair (Figure 6H), cell cycle, cell division (Figure 6I), and OXPHOS (Figure 6J). In all cases, we found that fast-growing tumors are transcriptionally closer to parental NM1 KO MEFs which supports the results from the correlation matrix clustering between samples and suggests NM1 loss in fast-growing tumor has an even more profound effect on the expression of critical genes. We, therefore, conclude that NM1 KO MEFs are capable of forming tumors in mice with different extent of expression of NM1-associated genes and the metabolic status of cells is directly associated with the tumor growth. Finally, NM1 is an alternatively spliced variant of Myo1C but RNA-seq profiling does not distinguish between the isoforms. The Myo1C profile, therefore, represents the overall abundance of all isoforms in a given tissue. Interestingly, the tumorigenic transformation of NM1 KO MEFs led to a further decrease of Myo1C in both tumors suggesting some general mechanism of tumorigenesis progression via reducing NM1/Myo1C levels. We, therefore, looked at the mutagenesis rate (Figure 6L) and deregulation of gene expression (Figure 6M) of Myo1C, p53, and mTOR in human cancer samples collected in the COSMIC Catalogue of Somatic Mutations in Cancer (version COSMIC v96) (Tate et al., 2019). While p53 shows a very high mutagenic rate in the majority of cancers, Myo1C and mTOR show relatively low levels of mutagenesis in different cancer tissues (Figure 6L). However, gene expression analysis shows that Myo1C is predominantly downregulated in several types of cancer, with ovarian cancer being the most prevalent (50% of cases have reduced expression of Myo1C), followed by the large intestine, kidney, lung, urinary tract, and breast cancers. p53 gene as a tumor suppressor, shows a similar pattern to Myo1C with reduced expression in the majority of cancers, while mTOR is predominantly overexpressed in most of the cancer tissues (Figure 6M).

**Figure 5.**
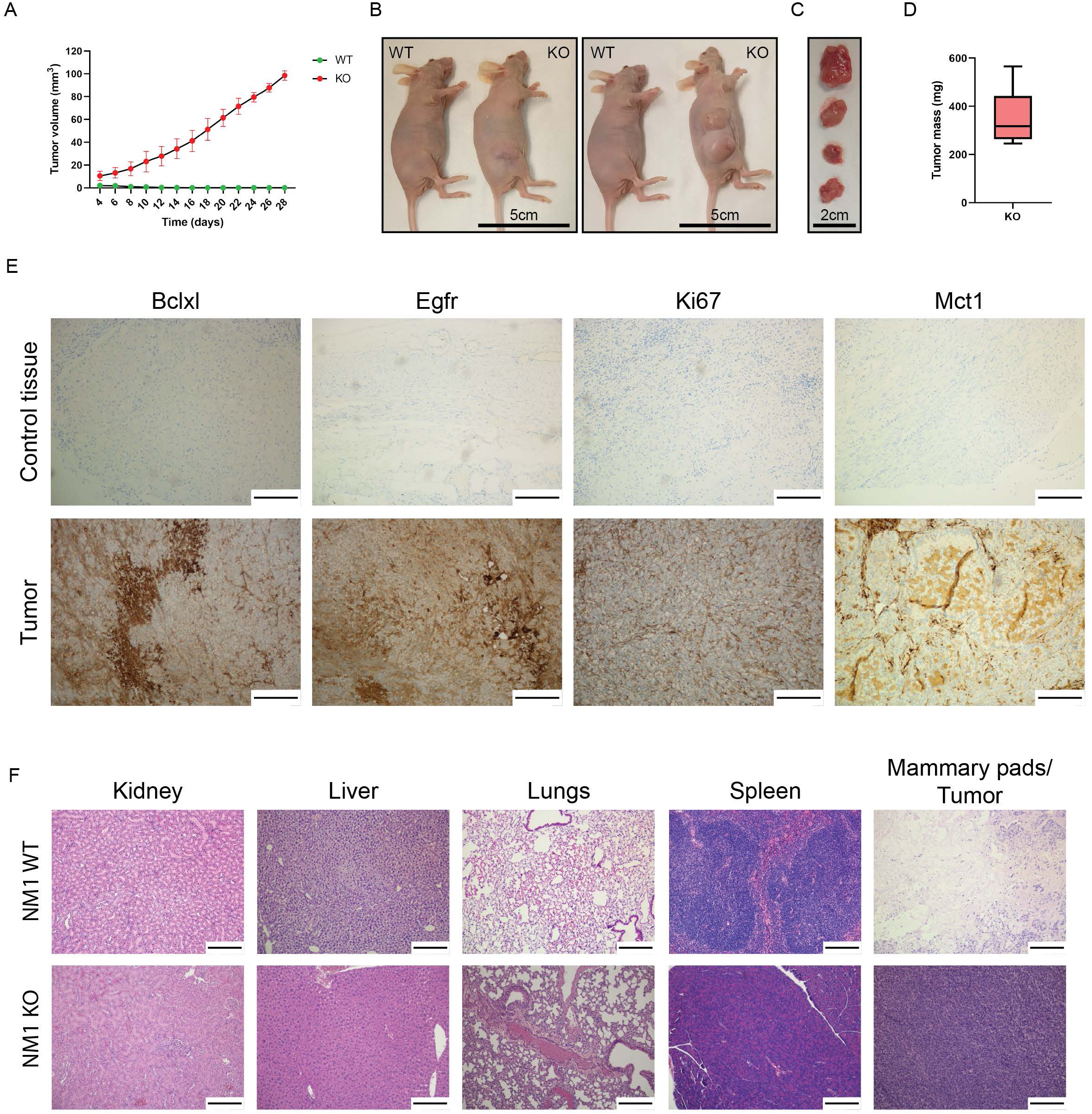
NM1 is a novel tumor suppressor. (A) The tumor growth rate over the period of 28 days in Balb/c nude mice injected with WT and NM1 KO cells. (B) Two representative pictures of Balb/c nude mice 28 days after injection of WT and NM1 KO cells under mammary pads tissue. WT and KO mark NM1 cell type injected into a given mouse. (C) Representative tumor tissues isolated from Balb/c nude mice 28 days after injection with NM1 KO cells. (D) Tumor mass measurement of isolated tumors from NM1 KO cell-injected Balb/c nude mice. (E) Immunohistochemistry staining with antibodies against cancer markers Bclxl, Egfr, Ki-67, and Mct1 of control tissue isolated from the place of injection of NM1 WT cells and tumor tissue isolated from mice injected with NM1 KO cells. The scale bar is 250µm. (F, H, E) Staining of sections of vital organs from mice injected with NM1 WT and KO cells. The scale bar is 250µm. Mammary pads tissue isolated from the area of injection of NM1 WT cells was used as a negative control to tumor tissue isolated from NM1 KO-injected mice. The scale bar is 250µm.

**Figure 6.**
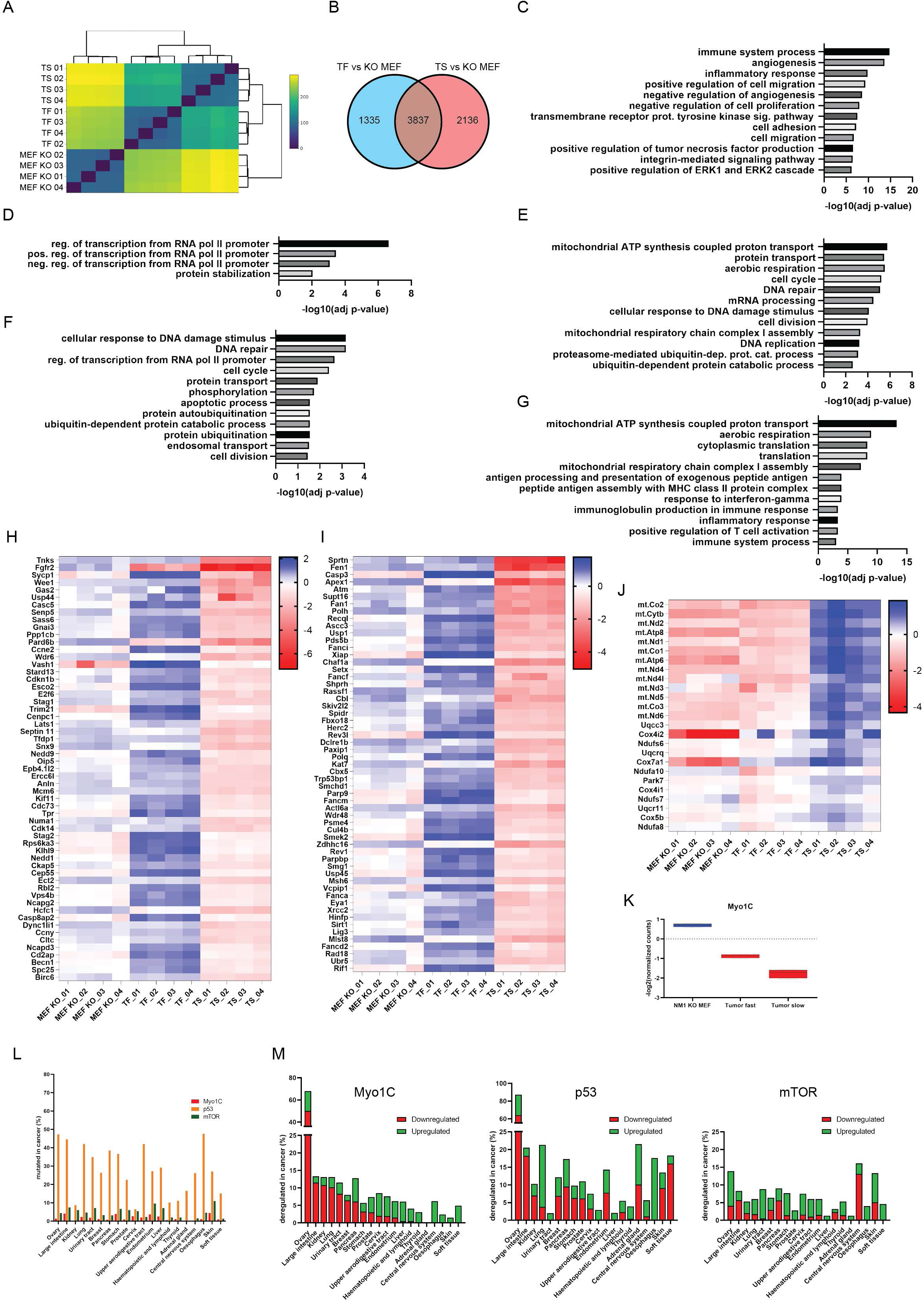
Transcriptomic profiling of tumor tissues derived from NM1 KO MEFs. (A) Correlation matrix clustering based on the similarity between NM1 KO MEFs and fast (TF) and slow (TS) growing tumors isolated from Balb/c nude mice injected with NM1 KO MEFs. (B) Venn diagram showing the number of all differentially expressed genes between NM1 KO MEFs and fast or slow-growing tumors, respectively. The intersection between the two groups represents the differentially expressed genes that are common during the tumorigenic transformation of NM1 KO MEFs to either fast or slow-growing tumors. (C) GO analysis of genes that are found to be differentially expressed during tumorigenic transformation of NM1 KO MEFs to both, fast and slow-growing tumors. The top 12 enriched biological process terms are shown in descending order. (D) GO analysis of genes that are found to be differentially expressed during the tumorigenic transformation of NM1 KO MEFs to fast-growing tumors. The only significantly enriched biological process terms are shown in descending order. (E) GO analysis of genes that are found to be differentially expressed during the tumorigenic transformation of NM1 KO MEFs to slow-growing tumors. The top 12 enriched biological process terms are shown in descending order. (F) GO analysis of differentially expressed genes that are upregulated in fast-growing tumors in comparison to slow-growing tumors. The top 12 enriched biological processes are shown in descending order. (G) GO analysis of differentially expressed genes that are downregulated in fast-growing tumors in comparison to slow-growing tumors. The top 12 enriched biological processes are shown in descending order. (H) Heatmap of differentially expressed genes between fast and slow-growing tumors associated with DNA damage response and repair. Heatmap represents the log2-normalized counts organized in ascending order (in slow-growing tumor samples) for each of 4 replicates of NM1 KO MEFs, fast-growing (TF) and slow-growing (TS) tumors. (I) Heatmap of differentially expressed genes between fast and slow-growing tumors associated with cell cycle and cell division. Heatmap represents the log2-normalized counts organized in ascending order (in slow-growing tumor samples) for each of 4 replicates of NM1 KO MEFs, fast-growing (TF) and slow-growing (TS) tumors. (J) Heatmap of differentially expressed genes between fast and slow-growing tumors associated with OXPHOS. Heatmap represents the log2-normalized counts organized in descending order (in slow-growing tumor samples) for each of 4 replicates of NM1 KO MEFs, fast-growing (TF), and slow-growing (TS) tumors. (K) Expression profile of Myo1C in NM1 KO MEFs, fast- and slow-growing tumors based on RNA-seq analysis. (L) The mutagenesis rate of Myo1C, p53, and mTOR in different human tissue cancer samples based on the COSMIC database. (M) The gene expression rate of Myo1C, p53, and mTOR in different human tissue cancer samples based on the COSMIC database.

We conclude that NM1 serves an essential role as a tumor suppressor and that NM1 deletion leads to changes in cell metabolism that are directly connected to tumorigenesis.

## Discussion

In the present study, we report that NM1 directly regulates the expression of specific mitochondrial transcription factors and forms a regulatory feedback loop with upstream signaling protein mTOR. Cells lacking NM1 show suppressed nutrient-stimulated PI3K/AKT/mTOR signaling pathways and reduced expression of mitochondrial transcription factors. This leads to mitochondrial phenotypic changes associated with a metabolic switch from OXPHOS to aerobic glycolysis which may serve as a potential underlying mechanism for solid tumor formation.

As translation of new proteins is an energetically heavy process, cells tend to keep signaling molecules to a minimum during starvation. In response to external stimuli, phosphorylation cascades are used to amplify the signal leading to a fast and adequate response. As the signal persists over time, cascades get saturated and cells cannot be further stimulated. Therefore, every pathway has several positive and or negative feedback loops which allow for fast adjustment of a given pathway to cellular needs. We show evidence that NM1 is a key element involved in such a positive feedback loop with mTOR and it is, therefore, important for proper nutrient sensing in cells. In response to stimuli, the PI3K/AKT/mTOR pathway is activated and mTOR phosphorylates NM1 which subsequently activates the expression of mitochondrial transcription factors but also mTOR itself. This leads to the accumulation of more mTOR which activates more NM1, potentially until a stimulus persists. After the initial signal is lost, the phosphorylation cascade comes to a halt, NM1 does not activate mTOR expression any further and the mTOR protein levels return to basal levels (Figure 7). Interestingly, except for Akt kinase, the whole PI3K/Akt/mTOR signaling cascade is suppressed in NM1 KO cells, while previous studies suggested that upon inhibition of mTOR, PI3K, and AKT pathways are activated via insulin-like growth factor receptor alternative pathways (O’Reilly et al., 2006; Shi et al., 2005). This partially correlates with the observed increased Akt kinase expression in NM1 depleted cells, however, insulin-like growth factor/receptor, as well as members of the PI3K pathway, are suppressed suggesting some other mechanism is responsible for Akt kinase activation and suppression of PI3K pathway. An explanation could be that NM1, similarly to mTOR, transcriptionally regulates the expression of these genes. However, there is a possibility that NM1 has a direct effect on these proteins as it is directly associated with the plasma membrane-bound phospholipids and regulates plasma membrane dynamics and organization (Dzijak et al., 2012; Venit et al., 2013; Venit et al., 2016). For example, Myo1C has been shown to facilitate exocytosis and delivery of several proteins such as Glut4, Neph1, aquaporin2, or VEGFR2 receptor to the plasma membrane and it is plausible that deletion of NM1 would affect the distribution/expression of plasma membrane proteins such as IGF1R as well (Arif et al., 2011; Barile et al., 2005; Boguslavsky et al., 2012; Tiwari et al., 2013). Another possibility for NM1 regulation of the PI3K pathway could be via competitive binding to phosphatidylinositol (3,4)-bisphosphate (PIP2). PI3K phosphorylates membrane-bound PIP2 to phosphatidylinositol (3,4,5)-trisphosphate (PIP3) and the balance between PIP2 and PIP3 is critical for cellular homeostasis (Tariq and Luikart, 2021). Dysregulation of lipid signaling could therefore affect the PI3K levels as well. Both NM1 and Myo1C were shown to specifically bind to PIP2 at the plasma membrane and we showed that upon deletion of NM1, the amount of myosin molecules bound to PIP2 is reduced by half (Hokanson and Ostap, 2006; Venit et al., 2016). The abundance of free PIP2 molecules and lack of competition between plasma membrane-bound NM1 and PI3K could lead to an adjustment of PI3K expression for proper signaling and cellular homeostasis.

**Figure 7.**
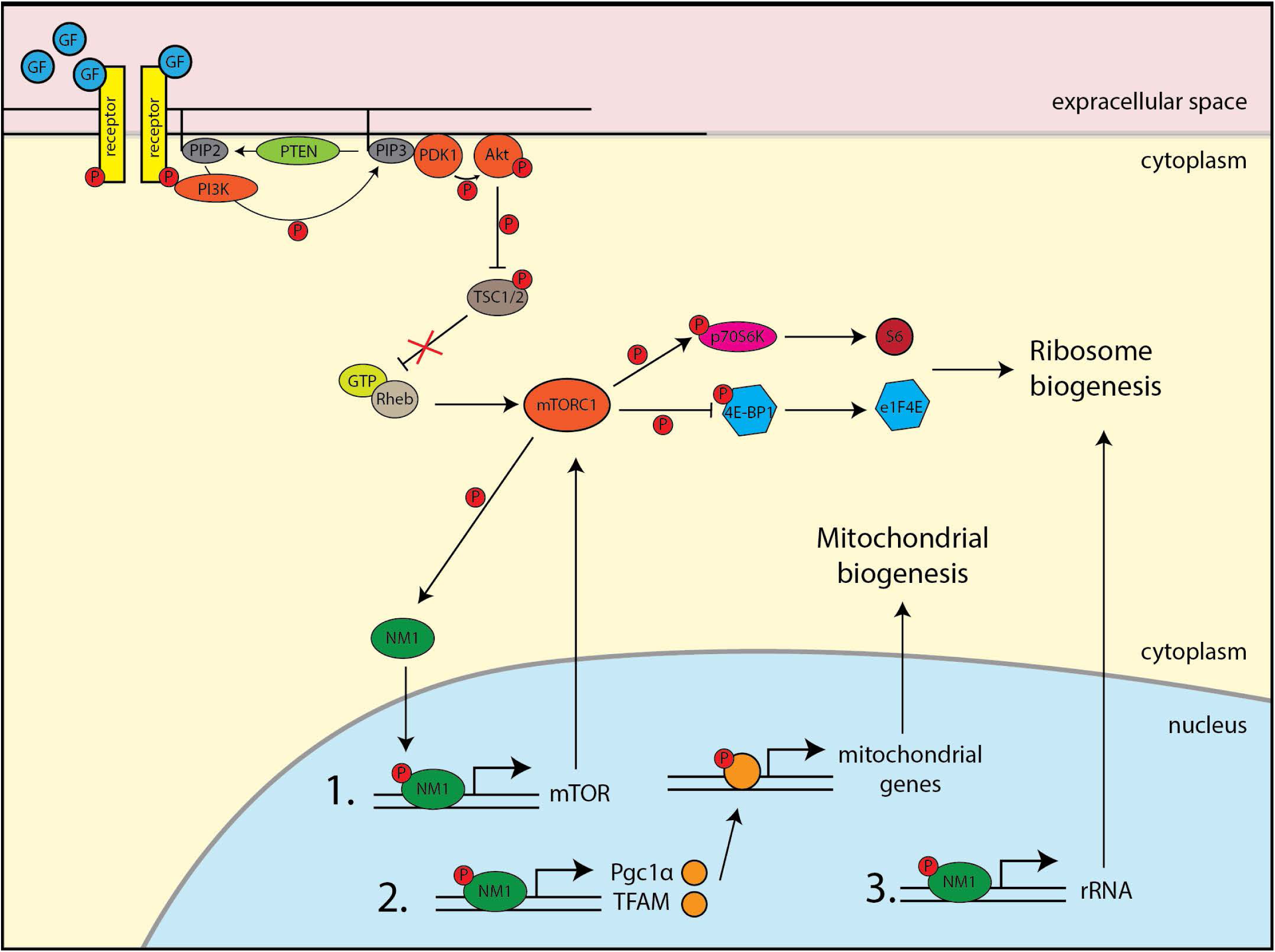
Proposed mechanism of NM1 action in PI3K/Akt/mTOR nutrient signaling. We propose that NM1 has a critical function as a regulator of cellular metabolism and it is part of the nutrient-sensing PI3K/Akt/mTOR pathway. Upon external stimulus, PI3K/Akt/mTOR cascade is activated leading to phosphorylation of NM1 which then transcriptionally regulates mTOR forming a positive feedback loop. This NM1-dependent mechanism is required to regulate the expression of mitochondrial transcription factors TFAM and PGC1α responsible for mitochondrial biogenesis and, as previously shown, rRNA synthesis.

The nutrient-sensing signaling network defined by PI3K, AKT, and mTOR controls several essential biological functions such as cellular growth, cell metabolism, and survival, and as a pro-proliferative pathway is often deregulated in cancers. It is therefore interesting that upon deletion of NM1, cells proliferate faster and can form tumors in a nude mouse model even though the PI3K/AKT/mTOR signaling pathway is suppressed. This could be at least partially explained by the previously described role of NM1 in cell cycle regulation (Sarshad et al., 2013; Venit et al., 2020b) although the possibility that NM1 affects other signaling pathways cannot be excluded. In favor of this, several studies suggested that even though often upregulated in cancers, targeting the mTOR pathway with its inhibitors brings only poor outcomes due to the promiscuity of signaling cascades and the plethora of possible targets which can be activated/deactivated depending on the microenvironment (Sun, 2021). Apart from insulin-like growth factor receptor-driven activation of PI3K and Akt mentioned above (O’Reilly et al., 2006; Shi et al., 2005), activation of other pathways such as DNA-PK or MAPK/ERK or inactivation of GSK3-dependent proteasomal degradation of oncogenic proteins can promote cancer cell survival even upon mTOR suppression (Carracedo et al., 2008; Koo et al., 2014; Li et al., 2013; Zhang et al., 2019). Accordingly, mTOR itself can be regulated via canonical AKT pathway or by Akt-independent Adenosine Monophosphate-activated Protein Kinase (AMPK) pathway (Lee et al., 2010), and depending on conditions, mTOR can promote transcriptional programs for both, OXPHOS and glycolysis (Fan et al., 2021; Morita et al., 2015; Rosario et al., 2019).

Another explanation for rapid tumor growth caused by NM1 KO cells can be due to a change in the tumor microenvironment. Several studies have indeed shown that metabolic resetting has an early active role in cellular reprograming into pluripotent or cancer stem cells and only after the initial switch to stemness metabolism can pluripotency transcriptional regulators induce additional factors to achieve stemness (Menendez, 2015; Prigione et al., 2014; Riggs et al., 2013; Son et al., 2013; Wegiel et al., 2018; Zhang et al., 2011; Zhu et al., 2010). Additionally, hypoxia, low nutrient content, and increased acidity due to an abundance of lactic acid not only affect cancer/stem cell metabolism but also influence surrounding cells and tissues which can further promote tumorigenesis (Buck et al., 2015; Mockler et al., 2014). Similarly, early epigenetic changes leading to downregulation of differentiation programs are a prerequisite for metabolic switch and achievement of stemness (Riggs et al., 2013), and suppressing mTOR in a timely manner by stemness and oncogenic transcription factor Sox2 is needed for stemness acquisition (Wang et al., 2013). As we have shown that loss of NM1 suppresses the global level of active histone marks associated with gene promoters and enhancers, and increases levels of heterochromatin histone marks (Venit et al., 2020b), and its deletion leads to a metabolic switch from OXPHOS to glycolysis, combination of these factors in NM1 KO cells may provide sufficient signal for cell transition to cancer gene programs even though they have suppressed PI3K/Akt/mTOR signaling pathway. We speculate that this can serve as a rescue mechanism for cancer cells during mTOR targeted cancer therapy as Rapamycin treatment leads to decreased binding of NM1 and active histone marks around transcription start sites of mitochondrial transcription factors similar to NM1 KO cells. The question of whether suppressing OXPHOS by NM1 depletion is sufficient for the metabolic switch to glycolysis by itself, or some other pro-glycolytic pathways must be activated remains to be elucidated. However, suppression of p53 protein was shown to induce expression of glucose transporters Glut1 and lactate/pyruvate transporter Mct1 leading to increased glucose metabolism and glycolysis (Boidot et al., 2012; Schwartzenberg-Bar-Yoseph et al., 2004) and we have shown that NM1 regulates the expression of p21 in cooperation with p53 (Venit et al., 2020b). As we have shown here that NM1 deletion leads to upregulation of Glut1 and Mct1 both in cells and tumors, it is plausible that deletion of NM1 not only directly suppresses OXPHOS via mitochondrial transcription factors but could also promote glycolysis via affecting the expression of some p53 target genes.

Glycolysis serves as a primary source of energy production in pluripotent and cancer stem cells and switching to OXPHOS is one of the defining hallmarks of gradual differentiation. NM1-directed repression could help to preserve the stemness of pluripotent stem cells as several studies showed that forced expression of glycolytic enzymes in pluripotent cells could protect them from differentiation (Tsogtbaatar et al., 2020). On the other hand, targeted activation of NM1 could push cells towards OXPHOS preventing the onset of cancer development or in the differentiation and maintenance of specific cell types such as neurons which are heavily dependent on OXPHOS. While this remains to be investigated, taken altogether our results suggest a novel role for NM1 as a tumor suppressor through a mechanism regulating mitochondrial transcription factors via mTOR signaling.

## Acknowledgments

This work is supported by grants from NYU Abu Dhabi, the Sheikh Hamdan Bin Rashid Al Maktoum Award for Medical Sciences, Cancerfonden (Swedish Cancer Society), and a donation from the Cipriani family to PP. We thank the NYU Abu Dhabi Center for Genomics and Systems Biology, in particular Marc Arnoux and Mehar Sultana for RNA sequencing, as well as Core Technology Platform Resources, including the NYU Abu Dhabi imaging center, in particular Rachid Rezgui for his help with the microscopy. We appreciate the computational platform provided by NYU Abu Dhabi HPC team and are especially thankful to Nizar Drou for technical help.

## Author Contributions

TV and PP conceived the research and wrote the manuscript. TV performed all RNA-seq analyses, imaging, and biochemical experiments and made the figures. OS performed High content screening and help with biochemical phenotyping of NM1 KO cells. WSA and YI performed metabolomics analysis, NHE and SRM performed ChIP-seq and all computational analysis, ST and RP performed electron microscopy experiments, and MM and PL analyzed NM1 KO tumors in nude mice. PP supervised the research. All authors read and approved the manuscript.

## Methods

### Cell culture, reagents, and antibodies

Nuclear Myosin 1 Knock-Out (NM1 KO) cell lines were derived from wild-type mouse embryonic fibroblasts (MEFs) (ATCC® CRL-2752) (NM1 WT) using the CRISPR/Cas9 system (Venit et al., 2020b). Cells were grown in a DMEM medium containing 10% fetal bovine serum, 100 U/ml penicillin, and 100 mg/ml streptomycin (Millipore-Sigma) in a humidified incubator with 5% CO_2_ at 37 °C. Hypoxia experiments were performed in a nitrogen pressured incubator to keep stable 5% CO_2_ and 1% O_2_ conditions for 48hrs before subsequent experiments. For drug treatments, cells were incubated with 100nM Rapamycin in a normal full DMEM medium for 2 hours to specifically inhibit the mTorc1 complex.

Antibodies against GAPDH (ab8245), The Total OXPHOS Rodent WB Antibody Cocktail kit (ab110413), Alexa Fluor 555 Goat Anti-Rabbit (ab150078), Horseradish peroxidase (HRP)-fused Goat Anti-Rabbit (ab6721) and Rabbit Anti-Mouse (ab6728) were purchased from Abcam. Antibodies against TFAM (ABE483) were purchased from Merc Millipore. Antibodies against EGFR (2-18C9) and Ki-67 (M1B1) were obtained from Agilent (Santa Clara, CA). The anti-MCT1 antibody was obtained from Merck (Rockville, MD). The anti-NM1 antibody was described and characterized earlier (Almuzzaini et al., 2015; Fomproix and Percipalle, 2004; Sarshad et al., 2013; Sarshad et al., 2014). Where applicable, all antibodies were used according to manufacturers’ protocols. MitoTracker™ Deep Red (M22426) and MitoTracker™ Orange (M7510) were purchased from ThermoFisher Scientific. Hoechst 43222 (H1399) and ProLong Gold Antifade Mountant with 4′,6- diamidino-2-phenylindole (DAPI; P36931) were purchased from Invitrogen.

### High-content phenotypic profiling

96-well clear-bottom assay plates (Corning) were used to culture cells at a density of 10,000 cells per well for the high-content phenotypic profiling. Cells were incubated with 200 nM MitoTracker™ Deep Red and Hoechst (4µM) for 30 minutes, washed twice with 1xPBS, fixed in 4% formaldehyde for 10 min, and permeabilized with 0.5% using Triton X-100 for 10 min. Cells were then stained with primary antibodies overnight and washed with 1xPBST (PBS with 0.5% Tween-20) three times. The washed cells were stained with secondary antibodies for 2 h, followed by Hoechst staining (5000x dilution) for 20 minutes, three washes with PBST, and storage in 1xPBS buffer. The plate was analyzed via the Cellomics ArrayScan XTI High-Content Screening platform (ThermoFisher Scientific), and image analysis was performed using the Compartment Analysis BioApplication software (Thermo Fisher Scientific). Primary objects (Circ) were defined using the Hoechst stained nuclei. Secondary masks were used to define mitochondria as “spots” with a positive fluorescent signal. Based on the analysis, the average fluorescence intensity or area of these spots was measured to quantify mitochondrial mass and distribution within the cell. For each experiment, 3 wells per condition, each containing at least 5000 cells were used and obtained data were plotted as a mean value for each well. Final graphs are compiled from two independent experiments.

### Western blotting

RIPA buffer (50 mM Tris-HCl pH 7.5, 150 mM NaCl, 1 mM EDTA, 1 % NP-40, 0.5 % sodium deoxycholate, and 0.1 % SDS) containing 1x cOmplete protease inhibitor cocktail (Roche) was used to collect total cellular lysates from MEFs. Pierce BCA protein assay kit (ThermoFisher Scientific) was used to measure protein concentration in all samples. 20μg of total protein per sample together with 1x Laemmli buffer was loaded to a 10% SDS-polyacrylamide gel electrophoresis (PAGE) gel and separated under reducing conditions. The separated proteins were transferred to a polyvinylidene difluoride membrane, after which the membrane was blocked with 3% milk in 1x TBST buffer (20 mM Tris pH 7.5, 150 mM NaCl, 0.1% Tween 20) for 1hr. Immunoblotting with primary antibodies was performed overnight at 4°C in a rotator, followed by 3 washes with 1x TBST, subsequent incubation with HRP-fused secondary antibodies for 4 hours at 4°C in a rotator, and 3 washes with 1xTBST. Protein bands were developed with ECL Western Blot Substrate (BioRad) and western blots were imaged by a ChemiDoc MP Imaging system (BioRad). The quantification of protein bands was performed by ImageJ software by comparing the signal from each antibody to the GAPDH signal which served as a loading control. Four blots were used for the quantification of each sample. For protein phosphorylation studies, SuperSep Phos-tag gel was used according to the manufacturer’s protocol (Fujifilm Wako).

### Quantitative RT-PCR

RNazol (Sigma) was used to extract total RNA from NM1 WT and KO MEFs according to the manufacturer’s protocol. The extract was cleaned from residual gDNA by Turbo DNA-free kit (Ambion). 1ug of RNA per sample was reverse transcribed using RevertAID First Strand cDNA Synthesis Kit (ThermoFisher Scientific) and diluted in water. The resulting cDNA was used as a template for quantitative PCR by Maxima SYBR Green qPCR Mix (ThermoFisher Scientific) with relevant primers for selected genes (Supplementary Table 1). Three-step cycling protocol was used to amplify the signal in the StepOnePlus Real-Time Thermal Cycler (Thermo Fisher Scientific). The expression data for each sample was normalized to Non-POU Domain Containing Octamer Binding (Nono) protein expression (Eissa et al., 2017). qPCR analysis for each gene was performed in triplicates and final results are compiled from at least 3 independent experiments.

### Chromatin immunoprecipitation and qPCR

Chromatin immunoprecipitation (ChIP) analysis was performed on WT and KO cells grown under normoxia or hypoxia conditions, and on cells treated with mTorc1-inhibitor Rapamycin. Approximately 2 million cells per ChIP were fixed with 1% formaldehyde, followed by 10 minutes of incubation with 0.125 M glycine to stop the reaction. Cells were lysed with the lysis buffer (10 mM Tris pH 8.0, 10 mM NaCl, 0.2% NP-40) supplemented with protease inhibitors and nuclei were recovered and resuspended in nuclei lysis buffer (50 mM Tris pH 8.1, 10 mM EDTA pH 8.0, 1% SDS, ddH2O) with protease inhibitors. Chromatin was sonicated by QSONICA sonicator using 4 rounds of DNA shearing on ice with a probe submerged into the bottom of the sample (70% Amplitude, 1 sec on/1sec off for 5 minutes). The final DNA fragment size was checked by DNA electrophoresis to be 200–500 bp. Sheared chromatin was divided equally for immunoprecipitation with antibodies fused to Dynabeads (Thermo Fisher Scientific) in IP Dilution buffer (20 mM Tris pH 8.1, 2 mM EDTA pH 8.0, 150 mM NaCl, 1% Triton X-100, 0.01% SDS, ddH2O). 10 µg anti-NM1, 2 µg anti-H3k9ac, and 2 µg of anti-H3K4me3 antibodies were used in each ChIP condition. 10% of sheared chromatin served as input control. Samples were incubated overnight, rotating at 4 °C. The IP samples were washed with IP Wash buffer 1 (20 mM Tris pH 8.1, 2 mM EDTA pH 8.0, 50 mM NaCl, 1% Triton X-100, 0.1% SDS, ddH2O) and IP Wash buffer 2 (20 mM Tris pH 8.1, 1 mM EDTA pH 8.0, 0.25 M LiCl, 1% NP-40, 1% sodium deoxycholate monohydrate, ddH2O). Reverse crosslinking and elution of DNA were performed by using the elution buffer (100 mM NaHCO_3_, 1% SDS, ddH2O). The samples were incubated with 5 M NaCl and 10 mg/ml of RNase A at 65 °C for 1 h. Proteinase K (20 mg/ml) was then added and incubated at 55 °C for 2 h. The samples were then placed on a magnet and the immunoprecipitated DNA samples in the supernatant along with the input samples were purified using ChIP Purification Kit (Zymo Research), according to the manufacturer’s protocol. They were diluted in 12 μl of elution buffer and the concentration was measured by Qubit (Thermo Fisher Scientific).

qPCR analysis was performed on diluted immunoprecipitated sample or input control in each reaction mixed with Maxima SYBR Green/Rox qPCR Master mix (Thermo Fisher Scientific) and appropriate set of primers (Supplementary Table 1), followed by a three-step cycling protocol in the StepOnePlus Real-Time Thermal Cycler (Thermo Fisher Scientific). Each graph represents combined data from 3 biological replicates, each performed in 4 technical replicates. The samples whose ct-value differed more than 0.5 cycles from the mean value for the given sample were removed from the analysis but a minimum of 9 measurements were performed for each experimental condition and each sample was normalized to the adjusted input.

### Mitochondrial mass and mitochondrial membrane potential measurement

To analyze differences in mitochondrial mass and membrane potential between NM1 WT and KO cells were seeded in 96 well plates (20,000 cells/well) and incubated with Hoechst (4µM) (Ex/Em 361/497) and 200nM MitoTracker™ Deep Red (Ex/Em 644/665) or 100nM MitoTracker™ Orange (Ex/Em 554/576) respectively, for 30 minutes. Cells were washed with 1xPBS twice, and the total fluorescence signal for each dye was measured with Synergy H1 Hybrid microplate Reader (BioTek). The signal obtained from the Mitotracker dyes was normalized to Hoechst staining. The visualized data represent the compilation of 3 independent experiments with 16 wells separately measured for each condition in each experiment.

### Microscopic methods

Cells grown overnight on glass slides were stained with 200 nM MitoTracker™ Deep Red FM for 20 min. Cells were washed twice with 1xPBS, fixed with 4% formaldehyde, permeabilized with 0.5% Triton X-100 in PBS for 15 min, and blocked with 1% BSA for 1 h. This was followed by staining with primary antibodies against TFAM (1:200) overnight. Cells were washed thrice with 1xPBST buffer and stained with Alexa Fluor 555 Goat Anti-Rabbit antibody (1:400) for 4 hours. After 3 additional washes with 1xPBST buffer, ProLong Gold anti-fade mounting media with DAPI (Invitrogen) was used to mount the coverslips to a glass slide. The Leica TCS SP8 STED 3X microscope equipped with HyD SMD2 detector and Leica HCPL APO CS2 63x/1.4 oil objective was used to acquire confocal images, the Software Leica application SuiteX was used to capture and analyze images, and the Huygens Professional software was used for deconvolution. Final data were processed using the Fiji software.

### Transmission Electron Microscopy (TEM) sample preparation

NM1 WT and KO cells were washed twice with PBS and trypsinized and 10 million cells for each genotype were collected and spun down at low speed to form a firm pellet. High-pressure freezing was performed using the Leica ICE high-pressure freezer apparatus with a gold-plated specimen carrier (carrier A, 3mm diameter, 100 μm deep, and carrier B with flat side down). Specimen carriers were lightly coated with 0.1% soy lecithin in chloroform to ensure a smooth and easy opening of the carrier without damaging the pellet. Carrier A was filled with well-pelleted cells and carrier B was placed on top with the flat side down before freezing at a programmed pressure of 2100 bars. After freezing, the sample pod was released automatically into a liquid nitrogen bath; the sample carrier was then separated from the specimen pod using precooled fine-tipped tweezers under liquid nitrogen and transferred to Leica freeze-substitution AFS2 set up in a 2 mL solution of cold dry absolute acetone (v/v) containing 1% osmium tetroxide. The AFS unit was slowly warmed from −90°C to 0°C (2°C/h), with the temperature being held at both −90°C for a period of 15hrs and thereafter at −60°C and −30°C for a period of 8hrs each. Samples were cleared of osmium by rinsing with absolute acetone (3 times × 5 mins) and thereafter infiltrated with low viscosity resin with increasing concentrations of 30% and 66% for 4h each, and 100% overnight. Individual samples were embedded in 1 mL of 100% low viscosity resin and polymerized for 30 h at 60°C. The resin blocks were tapered into a pyramid shape and polished using a Target Sample Preparation Unit (Leica TXP) and sectioned using Leica ultramicrotome (UC7) with a diamond knife at low speed to get 60nm-75 nm thick sections. Ultrathin sections were mounted on 200 mesh copper grids for imaging.

### Transmission Electron Microscopy (TEM) imaging and analysis

High-resolution transmission electron microscopy (HRTEM) images were obtained using a Talos F200X Scanning/Transmission Electron Microscope equipped with a CETA 16M camera at an accelerating voltage of 200 kV having a lattice-fringe resolution of 0.14 nm. All the relevant areas were marked using bright field (BF) imaging mode at spot size 5 and later scanned using the BF mode at spot size 5 and screen current between 3-4 nA. The data was analyzed using FIJI ImageJ software. For each sample, at least 150 individual mitochondria with clearly visible outer membrane were used for the analysis. A segmented line tool was used for manual tracking of mitochondrial parameters measurement. Mitochondrial perimeter/cristae length ratio was used to define the mass of the cristae and mitochondrial length/width aspect ratio to define the circularity of mitochondria.

### Intracellular calcium measurement

Intracellular calcium was measured by using Indo-1 AM Calcium Sensor Dye (ThermoFisher Scientific). The protocol was similar to the previous measurement of mitochondrial mass, just cells were stained separately with Hoechst (4µM) (Ex/Em 361/497) and Indo-1 AM (1µM) (Ex/Em 346/475) due to similar excitation/emission spectra. Hoechst staining in control wells was used for checking seeding density between the samples and for normalization of Indo-1 AM dye in experimental wells. The visualized data represent the compilation of 3 independent experiments with 16 wells separately measured for each condition in each experiment.

### Metabolites measurement

For metabolites measurements, L-Lactate Assay Kit (ab65330), Pyruvate Assay Kit (ab65342), ADP assay kit (ab83359), ATP Assay Kit (ab83355), and Hydrogen Peroxide Assay Kit (ab102500) were purchased from Abcam and performed according to manufacturer’s protocols for the assays. In short, NM1 WT and KO MEFs were grown overnight in 6 well plates (300,000 cells/well), after which they were trypsinized, washed in PBS, and resuspended in the appropriate assay buffer. The protein concentration of each sample was quantified using the Pierce BCA protein assay kit (ThermoFisher Scientific). The amount of each metabolite was measured fluorometrically according to the manufacturer’s recommendations and normalized to the total protein concentration of each sample. Each experiment was repeated at least 3 times with 4 replicates per sample per assay and final graphs are compiled from all experiments together.

### Glucose uptake assay

MEFs were grown overnight in 96 well plates (20000 cells/well) in full DMEM medium containing glucose (4).5g/l), after which they were washed twice with 1xPBS and incubated in DMEM medium without glucose for 2hr, 37°C, 5%CO2. The medium was exchanged with fresh medium containing Hoechst stain (4µM) and 2-NBDG fluorescent glucose analog (100µg/ml) (ThermoFisher Scientific) and cells were incubated for additional 30 minutes. The fluorescent intensity was measured for both dyes and relative glucose uptake normalized to Hoechst staining. The experiment was repeated 4 times with 5 replicates per sample final graph is compiled from all experiments together.

### Intracellular pH measurement

Intracellular pH measurement was performed with pHrodo Green AM Intracellular pH Indicator (ThermoFisher Scientific) according to the manufacturer’s protocol. Cells were grown overnight in 96 well plates (20000 cells/well) in full DMEM medium followed by 30 minutes of incubation with Hoechst stain (4µM) and pHrodo green dye mixed with PowerLoad concentrate in phenol red-free DMEM medium. After that, cells were washed twice with 1xPBS, and fluorescence intensity was measured for both dyes in a phenol red-free medium. The higher the pHrodo green fluorescent signal, the more acidic intracellular pH is. The pHrodo fluorescence data were normalized to the Hoechst stain. The experiment was repeated 4 times with 5 replicates per sample final graph is compiled from all experiments together.

### Ultra-Performance Liquid Chromatography High-Resolution Mass Spectrometry

5 replicates of NM1 WT and KO cells were used for metabolomic profiling. Overnight grown cells (approximately 3x10^5^ per sample) were washed twice with ice-cold 0.9% NaCl solution and 300µl of 100% Methanol was added to each well. After 3 minutes of incubation on ice, cells were scraped using a precooled cell scraper and moved to a cold Eppendorf tube. Cell extracts were spun down at 15000rpm for 15 min at 4°C and 200µl of each supernatant was moved to a new tube for further processing. Next, samples were vacuum dried and reconstituted in 200µl of cyclohexane/water (1:1) solution for subsequent analysis by Ultra Performance Liquid Chromatography High-Resolution Mass Spectrometry (UPLC-HRMS) performed at the VIB Metabolomics core facility (Belgium). 10 ul of each sample were injected on a Waters Acquity UHPLC device connected to a Vion HDMS Q-TOF mass spectrometer. Chromatographic separation was carried out on an ACQUITY UPLC BEH C18 (50 × 2.1 mm, 1.7 μm) column from Watersunder under the constant temperature of 40°C. A gradient of two buffers was used for separation: buffer A (99:1:0.1 water:acetonitrile:formic acid, pH 3) and buffer B (99:1:0.1 acetonitrile:water: formic acid, pH 3), as follows: 99% A for 0.1 min decreased to 50% A in 5 min, decreased to 30% from 5 to 7 minutes, and decreased to 0% from 7 to 10 minutes. The flow rate was set to 0. 5 mL min−1. Both positive and negative Electrospray Ionization (ESI) was applied to screen for a broad array of chemical classes of metabolites present in the samples. The LockSpray ion source was operated in positive/negative electrospray ionization mode under the following specific conditions: capillary voltage, 2.5 kV; reference capillary voltage, 2.5 kV; source temperature, 120°C; desolvation gas temperature, 600°C; desolvation gas flow, 1000 L h−1; and cone gas flow, 50 L h−1. The collision energy for the full MS scan was set at 6 eV for low energy settings, for high energy settings (HDMSe) it was ramped from 28 to 70 eV. Mass range was set from 50 to 1000Da, scan time was set at 0.1s. Nitrogen (greater than 99.5%) was employed as desolvation and cone gas. Leucine-enkephalin (250 pg/μL solubilized in water:acetonitrile 1:1 [v/v], with 0.1% formic acid) was used for the lock mass calibration, with scanning every 1 min at a scan time of 0.1 s. Profile data was recorded through Unifi Workstation v2.0 (Waters).

### Data curation and statistical analysis of the metabolomic data

Data normalization was performed to remove potential variation resulting from instrument inter-run tuning differences. Raw MS peak data representing the abundance of each detected compound were subject to median standardization and missing values were imputed by the minimum value. Compounds with missing values in more than 50% of samples were considered missing. Standardized data that passed the quality control step were then log2 transformed and IQR normalized using JMP Genomics v8 (SAS Institute, Cary, NC) to remove potential technical artifacts and outliers. PCA and hierarchical clustering were done to explore the correlation structure in the data across the two conditions (WT and KO).

### Functional and metabolic pathway enrichment analysis

Functional analysis of curated normalized peak data was performed using MetaboAnalyst v5.0 using an existing protocol (Pang et al., 2021). Implemented Gene Set Enrichment Analysis (GSEA) method in the Functional Analysis module of MetaboAnalyst v5.0 (Accessed in 2022 from http://www.metaboanalyst.ca/) was used to identify sets of functionally related compounds and evaluate their enrichment of potential functions defined by metabolic pathways. GSEA analysis of compounds identified using positive and negative ionization was performed separately. Putative annotation of MS peaks data considering different adducts and ion modes was performed. *m*/*z* values and retention time dimensions both were used to increase confidence in identifying compounds and improve the accuracy of functional interpretations. Annotated compounds were then mapped onto Mus musculus (mouse) [KEGG] (Mouse Genome Sequencing et al., 2002) and a curated 912 metabolic data sets predicted to change due to dysfunctional enzymes based on human metabolism, separately (Pang et al., 2021), for pathway activity prediction (Supplementary table 2). GSEA calculates the Enrichment score (ES) by walking down a ranked list of metabolites, increasing a running-sum statistic when a metabolite is in the metabolite set and decreasing it when it is not. A metabolite set is defined in this context as a group of metabolites with collective biological functions or common behaviors, regulations, or structures. In this method, the ES of each enriched pathway is calculated to reflect the degree to which a metabolite set is overrepresented at the top or bottom of a ranked list of metabolites. Each ES is then normalized by the average of all ES scores against all permutations of the expression dataset to generate normalized enrichment scores (NES) that are used to compare analysis results across metabolite sets. By normalizing the enrichment score, GSEA accounts for differences in metabolite set size and correlations between metabolite sets and the expression dataset. A positive NES indicates metabolite set enrichment at the top of the ranked list; a negative NES indicates metabolite set enrichment at the bottom of the ranked list (Subramanian et al., 2005). Finally, compound hits were identified for each enriched pathway.

### HiC-Seq sample preparation and analysis

NM1 WT and KO cells were grown under standard conditions to 80% confluency with two biological replicates used for each condition. After trypsinization, cells were washed twice in 1x PBS buffer and 1 million cells per condition were fixed with 2% formaldehyde for 10 mins. The cell pellets were washed twice by 1× PBS, fast-frozen on dry ice, and stored at −80 °C. All subsequent processing and Hi-C were performed by Genome Technology Center at NYU Langone Health, NY, using standard DNA extraction and library preparation protocols and the Arima Hi-C kit (Arima Genomics, San Diego, CA), respectively. Subsequently, sequencing libraries were prepared by using a modified version of the KAPA HyperPrep library kit (KAPA Biosystems, Willmington, MA) and sequenced by a NovaSeq instrument. Raw sequencing data were processed using the HiCUP (Wingett et al., 2015) pipeline and analyzed using HOMER (Heinz et al., 2010) (http://homer.ucsd.edu/homer/). For preprocessing with HiCUP, a digest file compatible with the Arima protocol was produced with HiCUP digester using the option –arima. Processed bam files produced by HiCUP were converted to HOMER format using the script hicup2homer followed by conversion to homer tag directories using the command makeTagDirectory -format HiCsummary. PCA analysis was performed using HOMER with the command runHiCpca.pl -genome mm10 -res 500000 - window 500000 followed by annotation and differential analysis with the scripts annotatePeaks.pl and getDiffExpression.pl. Bins changing PC1 values from positive to negative or vice versa with an FDR of less than 0.05 between WT and KO cells were classified as switching. Protein-coding genes with TSS overlapping each compartment were identified using the valr package (Riemondy et al., 2017). TADs were called on replicate-merged tag directories of WT-MEFs using the HOMER function findTADsAndLoops.pl with -window and -res set to 50000. The insulation score of the identified domains in all replicates of each condition was extracted using the command findTADsAndLoops.pl with the “score” option followed by differential analysis using getDiffExpression.pl. Domains showing the change in insulation score with an adjusted p-value less than 0.05 were classified as a differential. Data is available in the Gene Expression Omnibus (GEO) database under accession number GSE198989.

### ATAC-Seq sample preparation and analysis

NM1 WT and KO cells were used for the analysis performed commercially by Novogene (Beijing, China) with two biological replicates used for each condition. Isolated cell nuclei were mixed with Tn5 Transposase with two adapters for 30 min at 37 °C for DNA fragmentation, followed by DNA purification and amplification with a limited PCR cycle using index primers. Libraries were prepared according to recommended Illumina protocols and sequenced by NovaSeq 6000 instrument. ATAC-Seq processing and quality control were performed by Novogene (Beijing, China). Adapter trimming on raw fastq files was performed using trim-galore with default settings. Surviving paired reads were aligned against the relevant reference genome (GRCm38) using Burrows-Wheeler Aligner BWA-MEM. The Resulting BAM alignments were cleaned, sorted, and deduplicated (PCR and Optical duplicates) with PICARD tools (http://broadinstitute.github.io/picard). Processed bam files were converted to HOMER tag directories followed by annotation and differential analysis with the scripts annotatePeaks.pl and getDiffExpression.pl. ATAC-Seq peaks were called on cleaned, deduplicated bam files of both replicates of each condition together using macs2 with the parameters -q 0.05 -g mm/hg --keep-dup all --nomodel -- shift −100 --extsize 200 -B --broad -f BAMPE. Peaks of the two conditions being compared were merged using homer command mergePeaks and annotated with annotatePeaks.pl. Differential peaks were identified using the standard DESeq2 pipeline with lfcThreshold =1.5 and alpha=0.05. Data is available in the Gene Expression Omnibus (GEO) database under accession number GSE198988.

### ChIP-Seq sample preparation and analysis

Cells were crosslinked (two biological replicates per condition and input controls) using 1% formaldehyde for 10 min followed by quenching with 0.125 M Glycine for 5 min and lysis with lysis buffer 1 (50 mM Hepes KOH pH 7.5, 10 mM NaCl, 1 mM EDTA, 10% glycerol, 0.5% NP-40, 0.25% Triton X-100). Nuclei were pelleted, collected, and washed using lysis buffer 2 (10 mM Tris-HCl pH 8, 200 mM NaCl, 1 mM EDTA, 0.5 mM EGTA). This was followed by lysis using lysis buffer 3 (10 mM Tris-HCl pH 8; 100 mM NaCl, 1 mM EDTA; 0.5 mM EGTA; 0.1% Na-Deoxycholate, 0.5% N-laurylsarcosine). Chromatin was sheared using Qsonica Sonicator (4 cycles of 3 min at 70 % Amplitude), and then checked on 0.8% agarose gel. 100 μg of fragmented chromatin was mixed with the appropriate antibody. The protein-antibody immunocomplexes were recovered by the Pierce Protein A/G Magnetic Beads. Beads and attached immunocomplexes were washed twice using Low salt wash buffer (0.1% SDS; 2 mM EDTA, 1% Triton X- 100, 20 mM Tris-HCl pH 8, 150 mM NaCl), and High Salt wash buffer (0.1% SDS, 2 mM EDTA, 1% Triton X- 100, 20 mM Tris-HCl pH 8, 500 mM NaCl), respectively. The beads were resuspended in elution buffer (50 mM Tris-HCl pH 8, 10 mM EDTA, 1% SDS). De-crosslinking was achieved by adding 8 µL of 5 M NaCl and incubating at 65 °C overnight. RNase A (1 µL 10 mg/mL) was added for a 30 min incubation at 37 °C. Then, 4 µL 0.5 M EDTA, 8 µL 1 M Tris-HCl, and 1 µL 20 mg/mL proteinase K (0.2 mg/mL) were added for a 2-hour incubation at 42 °C to digest the chromatin. DNA was purified by a QIAquick PCR purification kit for qPCR analysis and sequencing. Raw reads were quality trimmed using Trimmomatic (Bolger et al., 2014) and analyzed with FastQC (http://www.bioinformatics.babraham.ac.uk/projects/fastqc) to trim low-quality bases, systematic base-calling errors, and sequencing adapter contamination. Specific parameters used were “trimmomatic_adapter.fa:2:30:10 TRAILING:3 LEADING:3 SLIDINGWINDOW:4:15 MINLEN:36”. Surviving paired reads were then aligned against the mouse reference genome (GRCm38) using Burrows-Wheeler Aligner BWA-MEM. The resulting BAM alignments were cleaned, sorted, and deduplicated (PCR and Optical duplicates) with PICARD tools (http://broadinstitute.github.io/picard). Bigwig files were generated using deeptools (Ramirez et al., 2016) command bamCoverage -bs 10 -e -- ignoreDuplicates –normalizeUsingRPKM after excluding encode blacklisted regions. Bigwig files were analyzed with the computeMatrix function of deeptools to plot the average signal around regions of interest. Peaks were called using macs2 (Zhang et al., 2008) on replicate-merged bam files with relevant input control and q=0.05. The –broad flag was used for H3K27me3 while narrow peaks were called for other epigenetic marks. Peaks of the two conditions being compared were merged using homer command mergePeaks followed by annotation and differential analysis with the scripts annotatePeaks.pl and getDiffExpression.pl. Peaks showing differential expression with an adjusted p-value less than 0.05 were classified as differentially expressed. Data is available in the Gene Expression Omnibus (GEO) database under accession number <u>GSE202716</u>.

### Animals work

Female (12-16 weeks old), Balb/c nude mice were purchased from Jackson Laboratory, USA. Nude mice were housed in a pathogen-free sterile ventilation system supplied with sterile woodchip and *ad libitum* feed and water supply. Mice were kept in a room maintained at a temperature of 21-24 °C on a schedule of 12 h light/dark cycle. All animal experiments were performed after approval by the NYUAD-IACUC (Protocol 21-0005).

### Tumor induction assay

A total of 10 female 16-week-old nude mice, 25-35 g in weight, were divided into two groups comprised of 5 mice per group. The mice were injected with NM1 WT and NM1 KO embryonic fibroblasts (0.1 mL, 3 x 10^6^ cells/100 µL in 10 mM PBS, pH 7.4) subcutaneously into the right flank of each nude mouse. Mice were weighed daily and monitored for mobility, respiratory distress, and signs of pain for up to 30 days. A weight loss of more than 20% was deemed to be unacceptable and would lead to the early sacrifice of the mouse. The tumor size was measured by caliper in each mouse daily and at the end of 30 days cycle, all mice from each group were euthanized by CO_2_ exposure and perfused with isotonic PBS followed by fixation with 10% neutral buffered formalin. Tumors were rapidly removed, weighed, and kept in 10% neutral buffered formalin for subsequent analysis (Palanikumar et al., 2020).

### Immunohistochemistry and histopathology

To analyze the tumor induction, the expression of Mct1, Egfr, Bclxl, and Ki-67 tumorigenic factors was addressed by immunohistochemistry analysis (IHC). Tumors were excised after the end of the tumor growth (after 30 days) and fixed using the neutral buffered formalin followed by the paraffin embedding. The embedded tumor samples were cut into 4 μm thick sections with a microtome (Leica microtome) and mounted on glass slides, deparaffinized by incubation in 3% methanol-hydrogen peroxide solution, followed by treatment with 10 mM EDTA (pH 8.0) at 95 °C. Then the slides were cooled to room temperature, rinsed with a phosphate-buffered saline solution containing 0.05% of tween 20 (PBST), and incubated with individual primary antibodies (Egfr, Ki-67, E-cadherin, Bcl-xl, Mct1) for 2 h. Subsequently, the slides were rinsed with PBST and incubated with the appropriate HRP-conjugated secondary antibody at room temperature for 30 min, counterstained with hematoxylin, dehydrated in ethanol, and mounted with DPX mounting media for further microscopic analysis. For the histopathological analysis, tissues from the lungs, liver, spleen, kidney, and heart along with tumor tissue were fixed with neutral buffered formalin, paraffin-embedded, and sectioned for hematoxylin and eosin staining (H & E). The tissues were then examined by light microscopy (Palanikumar et al., 2021).

### RNA-Seq library preparation, sequencing, and analysis

Total RNA from 4 replicates of NM1 KO MEFs was extracted with TRI Reagent (Millipore-Sigma) according to the manufacturer’s protocol. For transcriptional profiling of tumor tissues, the biggest (fast-growing tumor, TF), and the smallest (slow-growing tumor, TS) tumors were selected for comparison. Each tumor was divided into 4 equal pieces and each piece was separately homogenized in TRI Reagent by using the Bead Ruptor 96 Homogenizer (Omni International). The RNA-Seq library was prepared by using the NEBNext Ultra II RNA Library Prep Kit for Illumina (NEB) and sequenced with the NextSeq 500/550 sequencing platform (performed at the NYUAD Sequencing Center). All of the subsequent analysis, including quality trimming, was executed using the BioSAILs workflow execution system. Trimmomatic (version 0.36) was used for quality trimming of the raw reads to get rid of low-quality bases, systematic base-calling errors, as well sequencing adapter contamination (Bolger et al., 2014). The quality of the sequenced reads pre/post quality trimming was assessed by FastQC and only the reads that passed quality trimming in pairs were retained for downstream analysis (https://www.bioinformatics.babraham.ac.uk/projects/fastqc/). The quality-trimmed RNA-Seq reads were aligned to the Mus musculus GRCm38 (mm10) genome using HISAT2 (version 2.0.4) (Kim et al., 2015). The conversion and sorting of SAM alignment files for each sequenced sample to BAM format were done by using SAMtools (version 0.1.19) (Li et al., 2009). The BAM alignment files were processed using HTseq-count, using the reference annotation file to produce raw counts for each sample. The raw counts were then analyzed using the online analysis portal NASQAR (http://nasqar.abudhabi.nyu.edu/), to merge, normalize, and identify differentially expressed genes (DEG). DEG by at least twofold (log2(FC) ≥ 1 and adjusted p-value of <0.05 for upregulated genes, and log2(FC) ≤ −1 and adjusted p-value of <0.05 for downregulated genes) between the NM1 WT and KO MEFs were subjected to GO enrichment using DAVID Bioinformatics (https://david.ncifcrf.gov/) (Huang da et al., 2009). RNA-Seq data on stable NM1 KO MEFs and tumors derived from these cells were deposited in the Gene Expression Omnibus (GEO) database under accession number (GSE206858).

The RNA-Seq data set of NM1 WT and KO primary mouse embryonic fibroblast used in this study (Figure 1E, 1F, and 1G), was described previously (Venit et al., 2020b) and is available in the Gene Expression Omnibus (GEO) database under accession number GSE133506. The parameters and subsequent analysis used to define differentially expressed genes were the same as for RNA-seq analysis of tumor tissues. The list of genes used for comparative analysis was based on GO terms “oxidative phosphorylation” (GO:0006119) and “mitochondrion” (GO:0005739) (http://geneontology.org) (Ashburner et al., 2000; Gene Ontology, 2021).

