## Supplemental information for "Regulation of oxidative phosphorylation by Nuclear myosin 1 protects cells from metabolic reprogramming and tumorigenesis in mice"

Supplementary figure 1

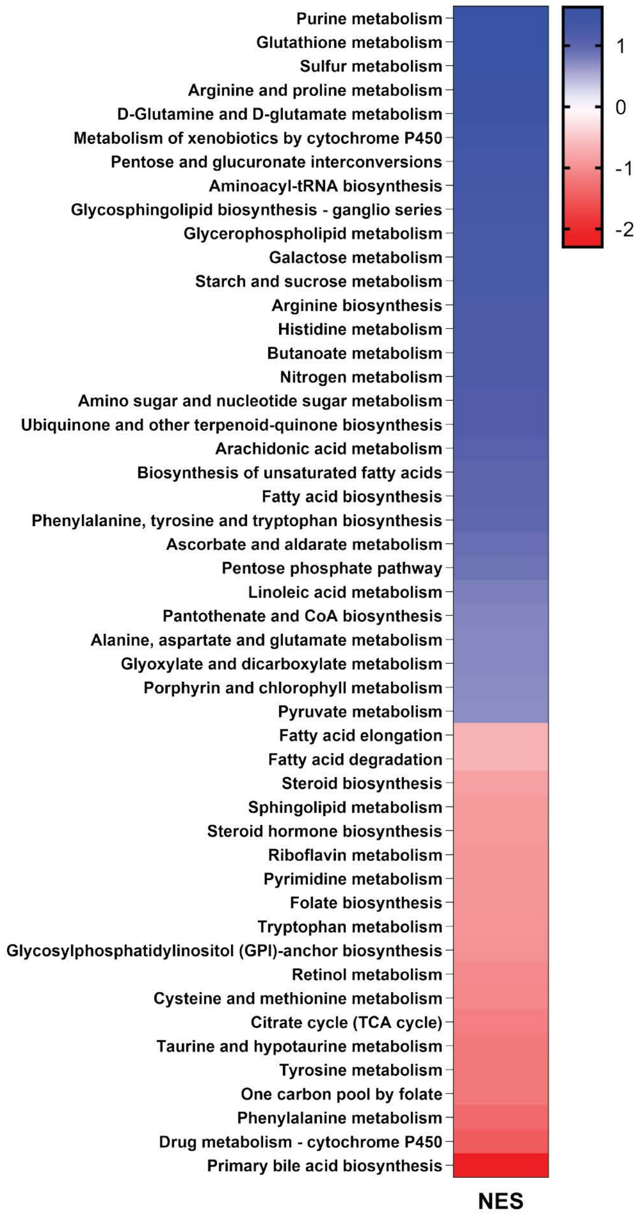

Supplementary figure 2

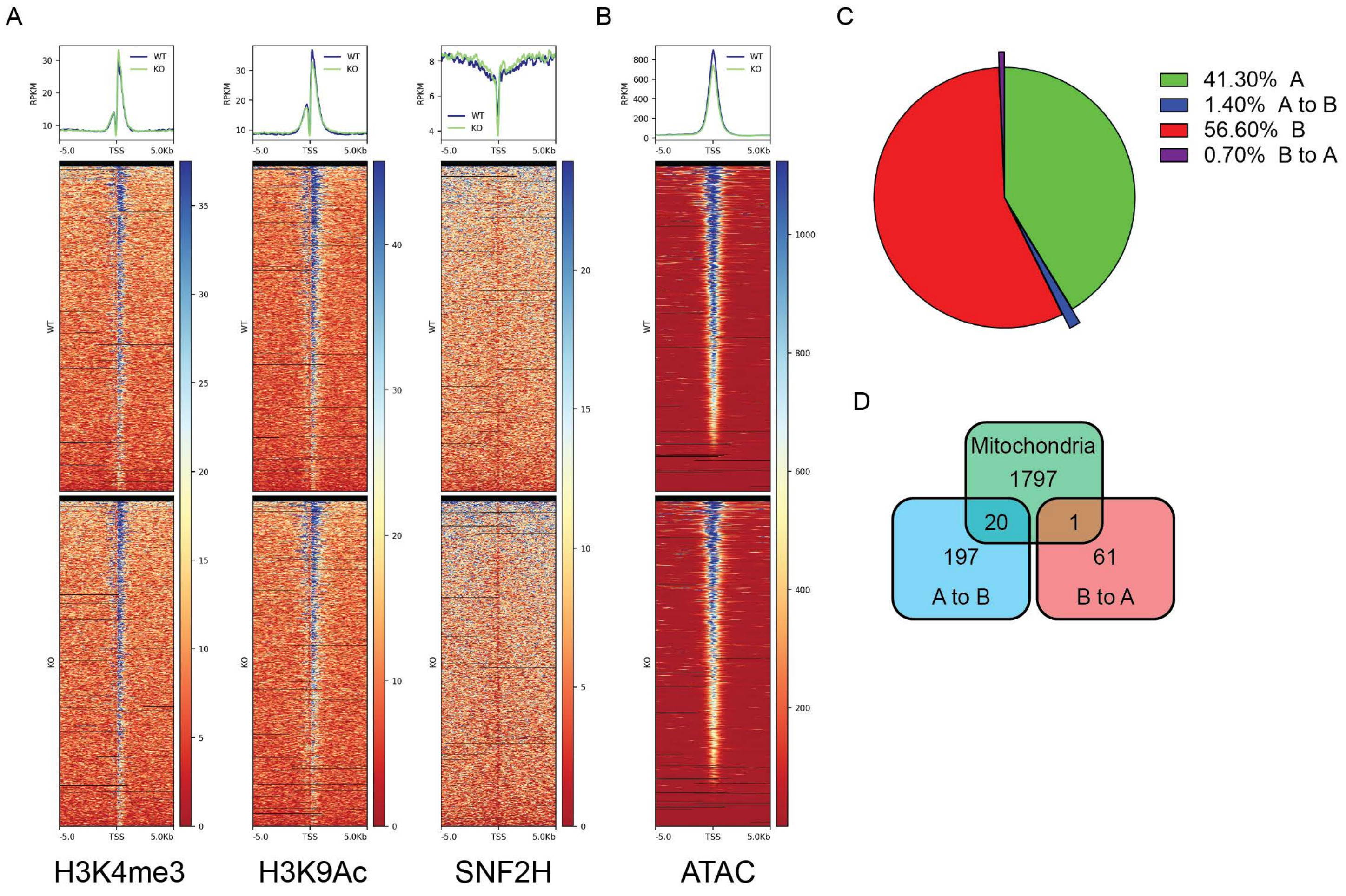

**Supplementary Table 1**

| Primer name | Type of analysis | Primer sequence |
|---|---|---|
| qNono F | Rt-qPCR | GCCAGAATGAAGGCTTGACTAT |
| qNono R | Rt-qPCR | TATCAGGGGGAAGATTGCCCA |
| qNdufs3-C1 F | Rt-qPCR | TGGCAGCACGTAAGAAGGG |
| qNdufs3-C1 R | Rt-qPCR | CTTGGGTAAGATTTCAGCCACAT |
| qSDHA-C2 F F | Rt-qPCR | GGAACACTCCAAAAACAGACCT |
| qSDHA-C2 F R | Rt-qPCR | CCACCACTGGGTATTGAGTAGAA |
| qUQCRB-C3 F | Rt-qPCR | GGCCGATCTGCTGTTTCAG |
| qUQCRB-C3 R | Rt-qPCR | CATCTCGCATTAACCCCAGTT |
| qCox5a-C4 F | Rt-qPCR | GCCGCTGTCTGTTCCATTC |
| qCox5a-C4 R | Rt-qPCR | GCATCAATGTCTGGCTTGTTGAA |
| qATP5D F | Rt-qPCR | TGTGACACTGGACATGCTG |
| qATP5D R | Rt-qPCR | TTCACTAGGGCTTCATTGGC |
| qUCP2 F | Rt-qPCR | ATGGTTGGTTTCAAGGCCACA |
| qUCP2 R | Rt-qPCR | CGGTATCCAGAGGGAAAGTGAT |
| MT-ND1 (mRNA) F | Rt-qPCR | TCGACCTGACAGAAGGAGAATCA |
| MT-ND1 (mRNA) R | Rt-qPCR | GGGCCGGCTGCGTATT |
| MT-CyB (mRNA) F | Rt-qPCR | AGACAAAGCCACCTTGACCC |
| MT-CyB (mRNA) R | Rt-qPCR | GATTGCTAGGGCCGCGATAA |

| | | |
|---|---|---|
| MT-CO1 F | Rt-qPCR | TTGCAACCCTACACGGAGGT |
| MT-CO1 R | Rt-qPCR | TCCGGTTAGACCACCAACTGT |
| q mt-16SrRNA F | Rt-qPCR | CAGGACATCCCAATGGTGTAG |
| q mt-16SrRNA R | Rt-qPCR | CTCTTGTCCTTTCGTACTGGG |
| MT-CyB gene F | Mitochondrial copy number | AGCCACCTTGACCCGATTCT |
| MT-CyB gene R | Mitochondrial copy number | CGTGGAGGAAGAGGAGGTGA |
| MT-ND1 gene F | Mitochondrial copy number | ACA CTT ATT ACA ACC CAA GAA CAC AT |
| MT-ND1 gene R | Mitochondrial copy number | TCA TAT TAT GGC TAT GGG TCA GG |
| MT-ATP6 gene F | Mitochondrial copy number | GCAGTCCGGCTTACAGCTAA |
| MT-ATP6 gene R | Mitochondrial copy number | GGTAGCTGTTGGTGGGCTAA |
| nuc-Terf gene F | Mitochondrial copy number | CTAGCTCATGTGTCAAGACCCTCTT |
| nuc-Terf gene R | Mitochondrial copy number | GCCAGCACGTTTCTCTCGTT |
| qFIS1 F | Rt-qPCR | AGCTGGTTCTGTGTCCAAG |
| qFIS1 R | Rt-qPCR | TGTTCCTCTTTGCTCCCTTTG |
| qDnm1L F | Rt-qPCR | TCCCAATTCCATTATCCTCGC |
| qDnm1L R | Rt-qPCR | CATCAGTACCCGCATCCATG |
| qMfn1 F | Rt-qPCR | CATTGCGTTTCGGTTTTCCC |
| qMfn1 R | Rt-qPCR | GAAGGAGCAGTAGGAGTTGAAG |
| qOpa1 F | Rt-qPCR | GTGTGCTGGAAATGATTGCTC |
| qOpa1 R | Rt-qPCR | TGGTGAGATCAAATTCCCGAG |
| qPINK1 F | Rt-qPCR | GACTCCCACCTTTCCCTTTG |

| | | |
|---|---|---|
| qPINK1 R | Rt-qPCR | GTAACTGCTCCATACTCTCCAG |
| qSnca F | Rt-qPCR | GGCAGTGAGGCTTATGAAATG |
| qSnca R | Rt-qPCR | TGGAACTGAGCACTTGTACG |
| qBecn1 F | Rt-qPCR | GTACCGACTTGTTCCCTATGG |
| qBecn1 R | Rt-qPCR | ACACAGTCCAGAAAAGCTACC |
| qSqstm1 F | Rt-qPCR | CCTATACCCACATCTCCCACC |
| qSqstm1 R | Rt-qPCR | TGTCGTAATTCTTGGTCTGTAGG |
| qHK2 F | Rt-qPCR | TCAAAGAGAACAAGGGCGAG |
| qHK2 R | Rt-qPCR | AGGAAGCGGACATCACAATC |
| qAldoa F | Rt-qPCR | CCCCAAGTTATCAAGTCCAAGG |
| qAldoa R | Rt-qPCR | GTTCAGACAGCCCATCCAG |
| qPkm F | Rt-qPCR | CCATTCTCTACCGTCCTGTTG |
| qPkm R | Rt-qPCR | TCCATGTAAGCGTTGTCCAG |
| qLDHA F | Rt-qPCR | GCTCCCCAGAACAAGATTACAG |
| qLDHA R | Rt-qPCR | TCGCCCTTGAGTTTGTCTTC |
| qEno3 F | Rt-qPCR | CTTCATCCAGAACTATCCCGTG |
| qEno3 R | Rt-qPCR | GAGGTCATCTCCCACAATCTG |
| qSlc2a1 F | Rt-qPCR | GATTGGTTCCTTCTCTGTCGG |
| qSlc2a1 R | Rt-qPCR | CCCAGGATCAGCATCTCAAAG |
| qEsr1 F | Rt-qPCR | AACCGCCCATGATCTATTCTG |
| qEsr1 R | Rt-qPCR | AGATTCAAGTCCCCAAAGCC |

| | | |
|---|---|---|
| qNr3c1 F | Rt-qPCR | TGACGTGTGGAAGCTGTAAAG |
| qNr3c1 R | Rt-qPCR | GACATTTTCGATAGCGGCATG |
| qEssra F | Rt-qPCR | GCCTCCAATGAGTGTGAGATC |
| qEssra R | Rt-qPCR | TTTGTACTTCTGCCGTCCG |
| qYy1 F | Rt-qPCR | GATACCTGGCATTGACCTCTC |
| qYy1 R | Rt-qPCR | ATAGCAGAGTTATCCCTGAACATC |
| qSrebf1 F | Rt-qPCR | CCATCGACTACATCCGCTTC |
| qSrebf1 R | Rt-qPCR | GCCCTCCATAGACACATCTG |
| qPpargc1a F | Rt-qPCR | CACCAAACCCACAGAAAACAG |
| qPpargc1a R | Rt-qPCR | GGGTCAGAGGAAGAGATAAAGTTG |
| qNRF1 F | Rt-qPCR | TTGATGGACACTTGGGTAGC |
| qNRF1 R | Rt-qPCR | TGTTCAGTTTGGGTCACTCC |
| Nrf2 F | Rt-qPCR | TCCCATTTGTAGATGACCATGAG |
| Nrf2 R | Rt-qPCR | CCATGTCCTGCTCTATGCTG |
| qTFAM F | Rt-qPCR | CACCCAGATGCAAAACTTTCAG |
| qTFAM R | Rt-qPCR | CTGCTCTTTATACTTGCTCACAG |
| qTfb2m F | Rt-qPCR | ACCAAAACCCATCCCGTC |
| qTfb2m R | Rt-qPCR | TCTGTAAGGGCTCCAAATGTG |
| qIGF1 F | Rt-qPCR | GAGACTGGAGATGTACTGTGC |
| qIGF1 R | Rt-qPCR | CTCCTTTGCAGCTTCGTTTTC |
| qIgf1r F | Rt-qPCR | TGCTGTCTATGTCAAGGCTG |

| | | |
|---|---|---|
| qIgf1r R | Rt-qPCR | AGAGGAAGAGTTTGATGCTGAG |
| qINSR F | Rt-qPCR | GGAAGCTACATCTGATTCGAGG |
| qINSR R | Rt-qPCR | TGAGTGATGGTGAGGTTGTG |
| qPik3c2a F | Rt-qPCR | AATGCCAGTGTGAAGGTCTC |
| qPik3c2a R | Rt-qPCR | CTGCCAACATCCACTTGATTC |
| qPik3r1 F | Rt-qPCR | GGATGCTGAATGGTACTGGG |
| qPik3r1 R | Rt-qPCR | TGTAAGAGTGTAATCGCCGTG |
| qPTEN F | Rt-qPCR | TGTAAAGCTGGAAAGGGACG |
| qPTEN R | Rt-qPCR | CCTCTGACTGGGAATTGTGAC |
| qAKT1 F | Rt-qPCR | GCCCTCAAGTACTCATTCCAG |
| qAKT1 R | Rt-qPCR | ACACAATCTCCGCACCATAG |
| qTSC2 F | Rt-qPCR | CTAAGGGTTTGCGTTCCAATG |
| qTSC2 R | Rt-qPCR | CAGGTGGAGAGTTATTCAGGC |
| qmTOR F | Rt-qPCR | ATTCAATCCATAGCCCCGTC |
| qmTOR R | Rt-qPCR | TGCATCACTCGTTCATCCTG |
| qp70S6K F | Rt-qPCR | ACAGGAGCAAATACTGGGAAG |
| qp70S6K R | Rt-qPCR | TCAGGTCCACAATGAAAGGG |
| qCh_TFAM_TSS F | ChIP-qPCR | TCCACATTCCCCGGAACAGC |
| qCh_TFAM_TSS R | ChIP-qPCR | ATTGCGTGAGACGAACCGGA |
| qCh_PPARGC1A_TSS F | ChIP-qPCR | CCAGCCTCCCTTCTCCTGTG |
| qCh_PPARGC1A_TSS R | ChIP-qPCR | TTGAAGCCTCCCAAAGGCCA |

| | | |
|---|---|---|
| qCh_MTOR_TSS F | ChIP-qPCR | AGGGTACCAAACCGTGGCTC |
| qCh_MTOR_TSS R | ChIP-qPCR | AACACTTCCGGTACGTCGCT |

**Supplementary Table 1.**

| Ionization | Pathway | Pathway_Total | Hits | P_val | P_adj | NegLog10Pval | NES | EC.Hits |
|---|---|---|---|---|---|---|---|---|
| Positive | fatty-acid--CoA ligase | 7 | 7 | 0.01333 | 0.05068 | 1.87517 | 1.864 | EC00013 |
| Positive | fatty acid transport via diffusion | 7 | 7 | 0.01333 | 0.05068 | 1.87517 | 1.864 | EC00028;EC00029;EC00030;EC00014;EC00013;EC00024;EC00027 |
| Positive | Beta oxidation of long chain fatty acid | 13 | 13 | 0.01282 | 0.05068 | 1.89211 | 1.817 | EC0004;EC00016;EC00017 |
| Positive | alpha-methylacyl-CoA racemase | 9 | 9 | 0.01316 | 0.05068 | 1.88074 | 1.792 | EC0004;EC00016;EC00017 |
| Positive | alpha-methylacyl-CoA racemase (reductase) | 9 | 9 | 0.01316 | 0.05068 | 1.88074 | 1.792 | EC0004;EC00016;EC00017 |
| Positive | Chenodeoxyglycocholate exchange | 9 | 9 | 0.01316 | 0.05068 | 1.88074 | 1.792 | EC0004;EC00016;EC00017 |
| Positive | FADH2 transporter, peroxisomal | 9 | 9 | 0.01316 | 0.05068 | 1.88074 | 1.792 | EC0004;EC00016;EC00017 |
| Positive | glycocholate exchange | 9 | 9 | 0.01316 | 0.05068 | 1.88074 | 1.792 | EC0004;EC00016;EC00017 |
| Positive | Taurocholic acid exchange | 9 | 9 | 0.01316 | 0.05068 | 1.88074 | 1.792 | EC0004;EC00016;EC00017 |
| Positive | acyl-Coenzyme A oxidase 2, branched chain | 9 | 9 | 0.01316 | 0.05068 | 1.88074 | 1.792 | EC0004;EC00016;EC00017 |
| Positive | hydroxysteroid (17-beta) dehydrogenase 4 | 9 | 9 | 0.01316 | 0.05068 | 1.88074 | 1.792 | EC0004;EC00016;EC00017 |
| Positive | 5-beta-cholestane-3-alpha,7-alpha,12-alpha-triol 27-hydroxylase | 9 | 9 | 0.01316 | 0.05068 | 1.88074 | 1.792 | EC0004;EC00016;EC00017 |
| Positive | peroxisomal thiolase 2 | 9 | 9 | 0.01316 | 0.05068 | 1.88074 | 1.792 | EC0004;EC00016;EC00017 |
| Positive | bile acid intracellular transport | 9 | 9 | 0.01316 | 0.05068 | 1.88074 | 1.792 | EC0004;EC00016;EC00017 |
| Positive | dimethylallyltranstransferase | 9 | 9 | 0.01316 | 0.05068 | 1.88074 | 1.792 | EC0004;EC00016;EC00017 |
| Positive | geranyltranstransferase | 9 | 9 | 0.01316 | 0.05068 | 1.88074 | 1.792 | EC0004;EC00016;EC00017 |

| | | | | | | | | |
|---|---|---|---|---|---|---|---|---|
| Positive | isopentenyl-diphosphate D-isomerase | 9 | 9 | 0.01316 | 0.05068 | 1.88074 | 1.792 | EC0004;EC00016;EC00017 |
| Positive | carnitine O-aceyltransferase, mitochondrial | 9 | 9 | 0.01316 | 0.05068 | 1.88074 | 1.792 | EC0004;EC00016;EC00017 |
| Positive | carnitine O-aceyltransferase, peroxisomal | 9 | 9 | 0.01316 | 0.05068 | 1.88074 | 1.792 | EC0004;EC00016;EC00017 |
| Positive | carnitine-propcarnitine carrier, peroxisomal | 9 | 9 | 0.01316 | 0.05068 | 1.88074 | 1.792 | EC0004;EC00016;EC00017 |
| Positive | cytochrome P450, family 7, subfamily A, polypeptide 1 | 9 | 9 | 0.01316 | 0.05068 | 1.88074 | 1.792 | EC0004;EC00016;EC00017 |
| Positive | taurine transport (sodium symport) (cytosol to peroxisome) | 9 | 9 | 0.01316 | 0.05068 | 1.88074 | 1.792 | EC0004;EC00016;EC00017 |
| Positive | taurochenodeoxycholate exchange | 9 | 9 | 0.01316 | 0.05068 | 1.88074 | 1.792 | EC0004;EC00016;EC00017 |
| Positive | transport into the mitochondria from cytosol (carnitine) | 9 | 9 | 0.01316 | 0.05068 | 1.88074 | 1.792 | EC0004;EC00016;EC00017 |
| Positive | Cytochrome P450 27 | 9 | 9 | 0.01316 | 0.05068 | 1.88074 | 1.792 | EC0004;EC00016;EC00017 |
| Positive | aldo-keto reductase family 1, member C4 (chlordecone reductase; 3-alpha hydroxysteroid dehydrogenase, type I; dihydrodiol dehydrogenase 4) | 9 | 9 | 0.01316 | 0.05068 | 1.88074 | 1.792 | EC0004;EC00016;EC00017 |
| Positive | aldo-keto reductase family 1, member D1 (delta 4-3-ketosteroid-5-beta-reductase) | 9 | 9 | 0.01316 | 0.05068 | 1.88074 | 1.792 | EC0004;EC00016;EC00017 |
| Positive | Very-long-chain-fatty-acid-CoA ligase | 9 | 9 | 0.01316 | 0.05068 | 1.88074 | 1.792 | EC00013;EC00014 |
| Positive | 5-beta-cytochrome P450, family 27, subfamily A, polypeptide 1 | 9 | 9 | 0.01316 | 0.05068 | 1.88074 | 1.792 | EC00013 |
| Positive | glycochenodeoxycholate exchange | 9 | 9 | 0.01316 | 0.05068 | 1.88074 | 1.792 | EC00013 |
| Positive | bile acid Coenzyme A: amino acid N-acyltransferase | 9 | 9 | 0.01316 | 0.05068 | 1.88074 | 1.792 | EC00044;EC00045;EC00028;EC00029;EC00030;EC00052 |
| Positive | sterol 12-alpha-hydroxylase | 9 | 9 | 0.01316 | 0.05068 | 1.88074 | 1.792 | EC00044 |

| | | | | | | | | |
|---|---|---|---|---|---|---|---|---|
| Positive | sterol 12-alpha-hydroxylase (nadh) | 9 | 9 | 0.01316 | 0.05068 | 1.88074 | 1.792 | EC00044 |
| Positive | hydroxy-delta-5-steroid dehydrogenase, 3 beta- and steroid delta-isomerase 7 | 9 | 9 | 0.01316 | 0.05068 | 1.88074 | 1.792 | EC00044 |
| Positive | taurine transport (sodium symport) (2:1) | 9 | 9 | 0.01316 | 0.05068 | 1.88074 | 1.792 | EC00045;EC00054 |
| Positive | Proline dehydrogenase | 6 | 6 | 0.01389 | 0.05068 | 1.8573 | 1.786 | EC00014 |
| Positive | Carbon monoxide exchange | 6 | 6 | 0.01389 | 0.05068 | 1.8573 | 1.786 | EC00014 |
| Positive | CO transporter via diffusion | 6 | 6 | 0.01389 | 0.05068 | 1.8573 | 1.786 | EC00014 |
| Positive | coproporphyrinogen oxidase (O2 required) | 6 | 6 | 0.01389 | 0.05068 | 1.8573 | 1.786 | EC00014 |
| Positive | Ferrochelatase, mitochondrial | 6 | 6 | 0.01389 | 0.05068 | 1.8573 | 1.786 | EC00014 |
| Positive | Heme oxygenase 1 | 6 | 6 | 0.01389 | 0.05068 | 1.8573 | 1.786 | EC00014 |
| Positive | Heme transport to cytosol | 6 | 6 | 0.01389 | 0.05068 | 1.8573 | 1.786 | EC00014 |
| Positive | hydroxymethylbilane synthase | 6 | 6 | 0.01389 | 0.05068 | 1.8573 | 1.786 | EC00018;EC00038 |
| Positive | iron (II) transport | 6 | 6 | 0.01389 | 0.05068 | 1.8573 | 1.786 | EC00015;EC00020 |
| Positive | Nad(p)h biliverdin reductase | 6 | 6 | 0.01389 | 0.05068 | 1.8573 | 1.786 | EC00021 |
| Positive | protoporphyrinogen IX mitochondrial transport | 6 | 6 | 0.01389 | 0.05068 | 1.8573 | 1.786 | EC00021 |
| Positive | uroporphyrinogen decarboxylase (uroporphyrinogen III) | 6 | 6 | 0.01389 | 0.05068 | 1.8573 | 1.786 | EC00022 |
| Positive | uroporphyrinogen-III synthase | 6 | 6 | 0.01389 | 0.05068 | 1.8573 | 1.786 | EC00022 |
| Positive | Nitric Oxide Synthase (NO forming) | 6 | 6 | 0.01389 | 0.05068 | 1.8573 | 1.786 | EC00022 |
| Positive | hydrogen peroxide transport via diffusion | 6 | 6 | 0.01389 | 0.05068 | 1.8573 | 1.786 | EC0005;EC0006 |
| Positive | o2 transport (diffusion) | 6 | 6 | 0.01389 | 0.05068 | 1.8573 | 1.786 | EC00018;EC00034;EC0005;EC0006;EC00020;EC00038 |

| | | | | | | | | |
|---|---|---|---|---|---|---|---|---|
| Positive | fatty acid intracellular transport | 6 | 6 | 0.01389 | 0.05068 | 1.8573 | 1.786 | EC00034;EC0005;EC0006 |
| Negative | hydroxy-delta-5-steroid dehydrogenase, 3 beta- and steroid delta-isomerase 7 | 6 | 6 | 0.03846 | 0.1916 | 1.41499 | 1.717 | EC00024 |
| Positive | lipid, flip-flop intracellular transport | 20 | 20 | 0.02564 | 0.09231 | 1.59108 | 1.7 | EC00034;EC0005;EC0006 |
| Positive | Glutathione dehydrogenase (dehydroascorbate reductase) | 9 | 9 | 0.01316 | 0.05068 | 1.88074 | 1.698 | EC0007;EC0008;EC0009;EC00054 |
| Positive | dehydroascorbate transport (uniport) | 5 | 5 | 0.02985 | 0.1033 | 1.52506 | 1.672 | EC00046;EC00047;EC00031;EC00049;EC00014;EC00028;EC00029;EC00030;EC0004;EC00016;EC00017;EC00024;EC00026;EC00027 |
| Positive | dihydroceramide desaturase | 7 | 7 | 0.02667 | 0.09351 | 1.57398 | 1.652 | EC00045 |
| Positive | 3-beta-hydroxysteroid-delta(8),delta(7)-isomerase | 11 | 11 | 0.01235 | 0.05068 | 1.90833 | 1.644 | EC00023 |
| Positive | C-3 sterol dehydrogenase (4-methylzymosterol) | 11 | 11 | 0.01235 | 0.05068 | 1.90833 | 1.644 | EC00023 |
| Positive | C-4 methyl sterol oxidase | 11 | 11 | 0.01235 | 0.05068 | 1.90833 | 1.644 | EC00044;EC00046;EC00047;EC0003;EC00031;EC00056;EC00055;EC0004;EC0001;EC0002;EC00026;EC00037 |
| Positive | acetyl-CoA carboxylase | 8 | 8 | 0.01316 | 0.05068 | 1.88074 | 1.644 | EC0007;EC0008;EC0009;EC00054;EC00010;EC00011 |
| Positive | protoporphyrinogen oxidase, mitochondrial | 14 | 14 | 0.01235 | 0.05068 | 1.90833 | 1.63 | EC00032 |
| Positive | C-14 sterol reductase | 14 | 14 | 0.01235 | 0.05068 | 1.90833 | 1.63 | EC0001;EC0002 |
| Positive | C-3 sterol keto reductase (zymosterol) | 14 | 14 | 0.01235 | 0.05068 | 1.90833 | 1.63 | EC0001;EC0002 |
| Positive | C-4 sterol methyl oxidase (4,4-dimethylzymosterol) | 14 | 14 | 0.01235 | 0.05068 | 1.90833 | 1.63 | EC0001;EC0002 |

| | | | | | | | | |
|---|---|---|---|---|---|---|---|---|
| Positive | cholesterol precursor intracellular transport | 14 | 14 | 0.01235 | 0.05068 | 1.90833 | 1.63 | EC0001;EC0002 |
| Positive | cytochrome P450 lanosterol 14-alpha-demethylase | 14 | 14 | 0.01235 | 0.05068 | 1.90833 | 1.63 | EC0003;EC00056;EC00042 |
| Positive | FAD transporter, endoplasmic reticulum | 14 | 14 | 0.01235 | 0.05068 | 1.90833 | 1.63 | EC0003;EC00056;EC00042 |
| Positive | FADH2 transporter, endoplasmic reticulum | 14 | 14 | 0.01235 | 0.05068 | 1.90833 | 1.63 | EC0003;EC00056;EC00042 |
| Positive | lanosterol synthase | 14 | 14 | 0.01235 | 0.05068 | 1.90833 | 1.63 | EC0003;EC00056;EC00042 |
| Positive | Previtamin D3 formation | 14 | 14 | 0.01235 | 0.05068 | 1.90833 | 1.63 | EC00023 |
| Positive | Squalene epoxidase, endoplasmic reticular (NADP) | 14 | 14 | 0.01235 | 0.05068 | 1.90833 | 1.63 | EC00023 |
| Positive | Squalene synthase | 14 | 14 | 0.01235 | 0.05068 | 1.90833 | 1.63 | EC00023 |
| Positive | Vitamin D3 exchange | 14 | 14 | 0.01235 | 0.05068 | 1.90833 | 1.63 | EC00023 |
| Positive | Vitamin D3 formation | 14 | 14 | 0.01235 | 0.05068 | 1.90833 | 1.63 | EC00023 |
| Positive | 7-dehydrocholesterol reductase | 14 | 14 | 0.01235 | 0.05068 | 1.90833 | 1.63 | EC00044;EC0003;EC00056;EC00055;EC00041;EC00049;EC0004;EC0001;EC0002;EC00025;EC00043;EC00037 |
| Positive | L-Phenylalanine,tetrahydrobiopterin:oxygen oxidoreductase (4-hydroxylating) | 8 | 8 | 0.02632 | 0.09349 | 1.57971 | 1.567 | EC00044;EC00045;EC00034;EC0005;EC0006;EC00015;EC00020 |
| Positive | 24-dehydrocholesterol reductase [Precursor] | 13 | 13 | 0.01282 | 0.05068 | 1.89211 | 1.519 | EC0004;EC00016;EC00017 |
| Positive | Lathosterol oxidase | 13 | 13 | 0.01282 | 0.05068 | 1.89211 | 1.519 | EC0004;EC00016;EC00017 |
| Positive | carnitine O-acetyltransferase, reverse direction, peroxisomal | 3 | 3 | 0.04545 | 0.1534 | 1.34247 | 1.473 | EC00014;EC00028;EC00029;EC00030;EC0004;EC00016;EC00017;EC00024;EC00027 |

| | | | | | | | | |
|---|---|---|---|---|---|---|---|---|
| Positive | carnitine-acetylcarnitine carrier, peroxisomal | 3 | 3 | 0.04545 | 0.1534 | 1.34247 | 1.473 | EC00014;EC00028;EC00029;EC00030;EC0004;EC00016;EC00017;EC00024;EC00027 |
| Positive | Reduced glutathione exchange | 5 | 5 | 0.07463 | 0.222 | 1.12709 | 1.46 | EC00014;EC00028;EC00029;EC00030;EC0004;EC00016;EC00017;EC00024;EC00027 |
| Positive | Iodide:hydrogen-peroxide oxidoreductase 4 | 5 | 5 | 0.08955 | 0.2467 | 1.04793 | 1.411 | EC00046;EC00047;EC00031;EC00049;EC00014;EC00028;EC00029;EC00030;EC0004;EC00016;EC00017;EC00024;EC00026;EC00027 |
| Positive | L-Thyroxine exchange | 5 | 5 | 0.08955 | 0.2467 | 1.04793 | 1.411 | EC00046;EC00047;EC00031;EC00049;EC00014;EC00028;EC00029;EC00030;EC0004;EC00016;EC00017;EC00024;EC00026;EC00027 |
| Positive | carboxylic acid dissociation | 6 | 6 | 0.09722 | 0.2652 | 1.01224 | 1.368 | EC00046;EC00047;EC00031;EC00049;EC00014;EC00028;EC00029;EC00030;EC0004;EC00016;EC00017;EC00024;EC00026;EC00027 |
| Positive | L-Ascorbate exchange | 3 | 3 | 0.1364 | 0.3506 | 0.86519 | 1.363 | EC00046;EC00047;EC00031;EC00049;EC00014;EC00028;EC00029;EC00030;EC0004;EC00016;EC00017;EC00024;EC00026;EC00027 |
| Positive | cysteinesulfinic acid oxidase | 3 | 3 | 0.1364 | 0.3506 | 0.86519 | 1.363 | EC00014;EC00028;EC00029;EC00030;EC0004;EC00016;EC00017;EC00024;EC00027 |
| Positive | Hypotaurine oxidase | 3 | 3 | 0.1364 | 0.3506 | 0.86519 | 1.363 | EC00014;EC00028;EC00029;EC00030;EC0004;EC00016;EC00017;EC00024;EC00027 |

| | | | | | | | | |
|---|---|---|---|---|---|---|---|---|
| Positive | Arachidic acid exchange | 3 | 3 | 0.1364 | 0.3506 | 0.86519 | 1.363 | EC00014;EC00028;EC00029;EC00030;EC0004;EC00016;EC00017;EC00024;EC00027 |
| Positive | Adrenaline secretion via secretory vesicle (ATP driven) | 4 | 4 | 0.0625 | 0.2033 | 1.20412 | 1.309 | EC00014;EC00028;EC00029;EC00030;EC0004;EC00016;EC00017;EC00024;EC00027 |
| Positive | dopamine beta-monooxygenase | 4 | 4 | 0.0625 | 0.2033 | 1.20412 | 1.309 | EC00014;EC00028;EC00029;EC00030;EC0004;EC00016;EC00017;EC00024;EC00027 |
| Positive | metanephrine secretion via secretory vesicle (ATP driven) | 4 | 4 | 0.0625 | 0.2033 | 1.20412 | 1.309 | EC00014;EC00028;EC00029;EC00030;EC0004;EC00016;EC00017;EC00024;EC00027 |
| Positive | pyridoxal transport via diffusion | 4 | 4 | 0.07812 | 0.222 | 1.10724 | 1.273 | EC00014;EC00028;EC00029;EC00030;EC0004;EC00016;EC00017;EC00024;EC00027 |
| Positive | pyridoxal kinase | 4 | 4 | 0.07812 | 0.222 | 1.10724 | 1.273 | EC00014;EC00028;EC00029;EC00030;EC0004;EC00016;EC00017;EC00024;EC00027 |
| Negative | Iodide:hydrogen-peroxide oxidoreductase | 1 | 1 | 0.3115 | 0.4547 | 0.50654 | 1.265 | EC00018 |
| Negative | Iodide:hydrogen-peroxide oxidoreductase 2 | 1 | 1 | 0.3115 | 0.4547 | 0.50654 | 1.265 | EC0004;EC00012 |
| Negative | L-Tyrosine carboxy-lyase | 1 | 1 | 0.3115 | 0.4547 | 0.50654 | 1.265 | EC0007 |
| Negative | Tyrosine:dopa oxidase | 1 | 1 | 0.3115 | 0.4547 | 0.50654 | 1.265 | EC0007 |
| Positive | 2,4 dihydroxy nitrophenol exchange | 2 | 2 | 0.1607 | 0.3981 | 0.79398 | 1.236 | EC00014;EC00028;EC00029;EC00030;EC0004;EC00016;EC00017;EC00024;EC00027 |

| | | | | | | | | |
|---|---|---|---|---|---|---|---|---|
| Positive | 4-Nitrophenol Sulfotransferase | 2 | 2 | 0.1607 | 0.3981 | 0.79398 | 1.236 | EC00014;EC00028;EC00029;EC00030;EC0004;EC00016;EC00017;EC00024;EC00027 |
| Positive | 4-Nitrophenyl sulfate exchange | 2 | 2 | 0.1607 | 0.3981 | 0.79398 | 1.236 | EC00014;EC00028;EC00029;EC00030;EC0004;EC00016;EC00017;EC00024;EC00027 |
| Positive | cytochrome P450 2E1 | 2 | 2 | 0.1607 | 0.3981 | 0.79398 | 1.236 | EC00014;EC00028;EC00029;EC00030;EC0004;EC00016;EC00017;EC00024;EC00027 |
| Positive | Glutathione:cystine oxidoreductase | 5 | 5 | 0.209 | 0.5037 | 0.67985 | 1.225 | EC00014;EC00028;EC00029;EC00030;EC0004;EC00016;EC00017;EC00024;EC00027 |
| Negative | intracellular transport | 1 | 1 | 0.3115 | 0.4547 | 0.50654 | 1.205 | EC00015 |
| Negative | Prostaglandin D2 exchange | 1 | 1 | 0.3115 | 0.4547 | 0.50654 | 1.205 | EC00018 |
| Negative | Prostaglandin-H2 D-isomerase [Precursor] | 1 | 1 | 0.3115 | 0.4547 | 0.50654 | 1.205 | EC0004;EC00012 |
| Negative | Thromboxane A2 exchange | 1 | 1 | 0.3115 | 0.4547 | 0.50654 | 1.205 | EC0004;EC00012 |
| Negative | Thromboxane-A synthase | 1 | 1 | 0.3115 | 0.4547 | 0.50654 | 1.205 | EC00018 |
| Negative | Prostaglandin G/H synthase | 1 | 1 | 0.3115 | 0.4547 | 0.50654 | 1.205 | EC00018 |
| Positive | 3,4-Dihydroxy-L-phenylalanine transport | 6 | 6 | 0.2222 | 0.5263 | 0.65326 | 1.195 | EC00014;EC00028;EC00029;EC00030;EC0004;EC00016;EC00017;EC00024;EC00027 |
| Positive | 3-Hydroxy-L-tyrosine carboxy-lyase | 6 | 6 | 0.2222 | 0.5263 | 0.65326 | 1.195 | EC00014;EC00028;EC00029;EC00030;EC0004;EC00016;EC00017;EC00024;EC00027 |
| Negative | taurine transport (sodium symport) (2:1) | 7 | 7 | 0.2453 | 0.4031 | 0.6103 | 1.174 | EC00023;EC00021;EC00022 |

| | | | | | | | | |
|---|---|---|---|---|---|---|---|---|
| Positive | fatty-acyl-CoA elongation (n-C18:3CoA) | 2 | 2 | 0.25 | 0.5769 | 0.60206 | 1.171 | EC00014;EC00028;EC00029;EC00030;EC0004;EC00016;EC00017;EC00024;EC00027 |
| Negative | Steryl-sulfatase | 3 | 3 | 0.254 | 0.4031 | 0.59517 | 1.168 | EC00011;EC0007;EC00015 |
| Negative | cytochrome c oxidase, mitochondrial Complex IV | 3 | 3 | 0.254 | 0.4031 | 0.59517 | 1.168 | EC00011;EC0007 |
| Negative | sulfite oxidase | 3 | 3 | 0.254 | 0.4031 | 0.59517 | 1.168 | EC00011;EC0007;EC00013;EC00015 |
| Negative | Cyanide sulfurtransferase, mitochondrial | 3 | 3 | 0.254 | 0.4031 | 0.59517 | 1.168 | EC00011;EC00015 |
| Negative | Cyanide transport via diffusion (mitochondrial) | 3 | 3 | 0.254 | 0.4031 | 0.59517 | 1.168 | EC00011;EC00015 |
| Negative | Thiocyanate transport via diffusion (mitochondrial) | 3 | 3 | 0.254 | 0.4031 | 0.59517 | 1.168 | EC0004;EC00012 |
| Negative | thiosulfate transport via sodium symport | 3 | 3 | 0.254 | 0.4031 | 0.59517 | 1.168 | EC00018 |
| Negative | Cyanide transport via diffusion (extracellular to cytosol) | 3 | 3 | 0.254 | 0.4031 | 0.59517 | 1.168 | EC00023 |
| Negative | Thiocyanate exchange | 3 | 3 | 0.254 | 0.4031 | 0.59517 | 1.168 | EC00023 |
| Negative | Thiocyanate transport via diffusion (cytosol to extracellular) | 3 | 3 | 0.254 | 0.4031 | 0.59517 | 1.168 | EC00023 |
| Positive | adenosylhomocysteinase | 8 | 8 | 0.3026 | 0.5871 | 0.51913 | 1.163 | EC00044;EC00045;EC00014;EC00028;EC00029;EC00030;EC00024;EC00052 |
| Positive | methionine adenosyltransferase | 8 | 8 | 0.3026 | 0.5871 | 0.51913 | 1.163 | EC00014;EC00028;EC00029;EC00030;EC00024;EC00027 |
| Positive | fatty acyl-CoA synthase (n-C8:0CoA), lumped reaction | 2 | 2 | 0.25 | 0.5769 | 0.60206 | 1.156 | EC00014;EC00028;EC00029;EC00030;EC0004;EC00016;EC00017;EC00024;EC00027 |

| | | | | | | | | |
|---|---|---|---|---|---|---|---|---|
| Negative | carboxylic acid dissociation | 1 | 1 | 0.459 | 0.6245 | 0.33819 | 1.145 | EC0005 |
| Negative | R group artificial flux | 1 | 1 | 0.459 | 0.6245 | 0.33819 | 1.145 | EC0005 |
| Negative | fatty acyl-CoA synthase (n-C8:0CoA), lumped reaction | 1 | 1 | 0.459 | 0.6245 | 0.33819 | 1.145 | EC0005 |
| Negative | fatty acyl-CoA synthase (n-C10:0CoA) | 1 | 1 | 0.459 | 0.6245 | 0.33819 | 1.145 | EC0005 |
| Negative | fatty-acyl-CoA synthase (n-C12:0CoA) | 1 | 1 | 0.459 | 0.6245 | 0.33819 | 1.145 | EC0005 |
| Negative | fatty-acyl-CoA synthase (n-C14:0CoA) | 1 | 1 | 0.459 | 0.6245 | 0.33819 | 1.145 | EC0005 |
| Positive | Tetrahydrobiopterin-4a-carbinolamine dehydratase | 9 | 9 | 0.2895 | 0.5871 | 0.53835 | 1.144 | EC00014;EC00028;EC00029;EC00030;EC0004;EC00016;EC00017;EC00024;EC00027 |
| Negative | 5'-nucleotidase (AMP), extracellular | 2 | 2 | 0.339 | 0.4913 | 0.4698 | 1.12 | EC0007 |
| Positive | fatty acyl-CoA desaturase (n-C20:3CoA -> n-C20:4CoA) | 1 | 1 | 0.4082 | 0.5871 | 0.38913 | 1.099 | EC00013;EC00044;EC00045;EC00052 |
| Positive | intracellular transport | 1 | 1 | 0.4082 | 0.5871 | 0.38913 | 1.099 | EC0001;EC0002;EC00026 |
| Positive | Prostaglandin D2 exchange | 1 | 1 | 0.4082 | 0.5871 | 0.38913 | 1.099 | EC0001;EC0002;EC00026 |
| Positive | Prostaglandin-H2 D-isomerase [Precursor] | 1 | 1 | 0.4082 | 0.5871 | 0.38913 | 1.099 | EC0001;EC0002;EC00026 |
| Positive | Thromboxane A2 exchange | 1 | 1 | 0.4082 | 0.5871 | 0.38913 | 1.099 | EC00043 |
| Positive | Thromboxane-A synthase | 1 | 1 | 0.4082 | 0.5871 | 0.38913 | 1.099 | EC00043 |
| Positive | Prostaglandin G/H synthase | 1 | 1 | 0.4082 | 0.5871 | 0.38913 | 1.099 | EC00044;EC00052 |
| Negative | fatty-acid--CoA ligase | 2 | 2 | 0.4576 | 0.6245 | 0.33951 | 1.068 | EC0007 |
| Negative | acetyl-CoA carboxylase | 2 | 2 | 0.4576 | 0.6245 | 0.33951 | 1.068 | EC0007 |

| | | | | | | | | |
|---|---|---|---|---|---|---|---|---|
| Positive | Vitamin D3 uptake | 3 | 3 | 0.3485 | 0.5871 | 0.4578 | 1.068 | EC00046;EC00047;EC00031;EC00049;EC00014;EC00028;EC00029;EC00030;EC00024;EC00026;EC00027 |
| Positive | ATP transporter, peroxisomal | 3 | 3 | 0.3485 | 0.5871 | 0.4578 | 1.068 | EC00014;EC00028;EC00029;EC00030;EC00024;EC00027 |
| Positive | O2 transport, endoplasmic reticulum | 3 | 3 | 0.3485 | 0.5871 | 0.4578 | 1.068 | EC00014;EC00028;EC00029;EC00030;EC00024;EC00027 |
| Positive | NADP transporter, peroxisome | 3 | 3 | 0.3485 | 0.5871 | 0.4578 | 1.068 | EC00014;EC00028;EC00029;EC00030;EC00024;EC00027 |
| Positive | NADPH transporter, peroxisome | 3 | 3 | 0.3485 | 0.5871 | 0.4578 | 1.068 | EC00014;EC00028;EC00029;EC00030;EC00024;EC00027 |
| Positive | Vitamin D-25-hydroxylase (D3) | 3 | 3 | 0.3485 | 0.5871 | 0.4578 | 1.068 | EC00014;EC00028;EC00029;EC00030;EC00024;EC00027 |
| Positive | sterol O-acyltransferase (acyl-Coenzyme A: cholesterol acyltransferase) 1 | 3 | 3 | 0.3485 | 0.5871 | 0.4578 | 1.068 | EC00014;EC00028;EC00029;EC00030;EC00024;EC00027 |
| Positive | 25-Hydroxyvitamin D3 exchange | 3 | 3 | 0.3485 | 0.5871 | 0.4578 | 1.068 | EC00014;EC00028;EC00029;EC00030;EC00024;EC00027 |
| Positive | cholesterol efflux (ATP depedent) | 3 | 3 | 0.3485 | 0.5871 | 0.4578 | 1.068 | EC00014;EC00028;EC00029;EC00030;EC00024;EC00027 |
| Positive | cholesterol ester exchange | 3 | 3 | 0.3485 | 0.5871 | 0.4578 | 1.068 | EC00014;EC00028;EC00029;EC00030;EC00024;EC00027 |
| Positive | Cholesterol exchange | 3 | 3 | 0.3485 | 0.5871 | 0.4578 | 1.068 | EC00014;EC00028;EC00029;EC00030;EC00024;EC00027 |
| Positive | cholesterol intracellular transport | 3 | 3 | 0.3485 | 0.5871 | 0.4578 | 1.068 | EC00014;EC00028;EC00029;EC00030;EC00024;EC00027 |
| Positive | diphosphomevalonate decarboxylase | 3 | 3 | 0.3485 | 0.5871 | 0.4578 | 1.068 | EC00014;EC00028;EC00029;EC00030;EC00024;EC00027 |

| | | | | | | | | |
|---|---|---|---|---|---|---|---|---|
| Positive | Hydroxymethylglutaryl CoA reductase (ir) | 3 | 3 | 0.3485 | 0.5871 | 0.4578 | 1.068 | EC00014;EC00028;EC00029;EC00030;EC00024;EC00027 |
| Positive | Hydroxymethylglutaryl-CoA reversible peroxisomal transport | 3 | 3 | 0.3485 | 0.5871 | 0.4578 | 1.068 | EC00046;EC00047;EC00031;EC00049;EC00014;EC00028;EC00029;EC00030;EC00016;EC00017;EC00024;EC00026;EC00027;EC0004 |
| Positive | mevalonate kinase (atp) | 3 | 3 | 0.3485 | 0.5871 | 0.4578 | 1.068 | EC00046;EC00047;EC00031;EC00049;EC00014;EC00028;EC00029;EC00030;EC00016;EC00017;EC00024;EC00026;EC00027 |
| Positive | phosphomevalonate kinase | 3 | 3 | 0.3485 | 0.5871 | 0.4578 | 1.068 | EC00046;EC00047;EC00031;EC00049;EC00014;EC00028;EC00029;EC00030;EC00016;EC00017;EC00024;EC00026;EC00027 |
| Positive | 7-alpha,24(S)-Dihydroxycholesterol exchange | 3 | 3 | 0.3485 | 0.5871 | 0.4578 | 1.068 | EC00014;EC00028;EC00029;EC00030;EC00024;EC00027 |
| Positive | 7-alpha,25-Dihydroxycholesterol exchange | 3 | 3 | 0.3485 | 0.5871 | 0.4578 | 1.068 | EC00014;EC00028;EC00029;EC00030;EC00024;EC00027 |
| Positive | 7-alpha,27-Dihydroxycholesterol exchange | 3 | 3 | 0.3485 | 0.5871 | 0.4578 | 1.068 | EC00046;EC00047;EC00031;EC00049;EC00014;EC00028;EC00029;EC00030;EC0004;EC00016;EC00017;EC00024;EC00027 |
| Positive | cholesterol 25-hydroxylase | 3 | 3 | 0.3485 | 0.5871 | 0.4578 | 1.068 | EC00024;EC00014;EC00028;EC00029;EC00030;EC00027 |
| Positive | cytochrome P450, family 46, subfamily A, polypeptide 1 | 3 | 3 | 0.3485 | 0.5871 | 0.4578 | 1.068 | EC00013 |
| Positive | oxysterol 7-alpha-hydroxylase | 3 | 3 | 0.3485 | 0.5871 | 0.4578 | 1.068 | EC00054 |
| Positive | 24 trihydroxy cholesterol transport | 3 | 3 | 0.3485 | 0.5871 | 0.4578 | 1.068 | EC00054 |

| | | | | | | | | |
|---|---|---|---|---|---|---|---|---|
| Positive | 25 trihydroxy cholesterol transport | 3 | 3 | 0.3485 | 0.5871 | 0.4578 | 1.068 | EC00054 |
| Positive | 27 trihydroxy cholesterol transport | 3 | 3 | 0.3485 | 0.5871 | 0.4578 | 1.068 | EC00022 |
| Positive | oxysterol 7alpha-hydroxylase | 3 | 3 | 0.3485 | 0.5871 | 0.4578 | 1.068 | EC00034 |
| Positive | L-Phenylalanine exchange | 12 | 12 | 0.4384 | 0.5873 | 0.35813 | 1.052 | EC00022 |
| Negative | ATP diphosphohydrolase | 3 | 3 | 0.4286 | 0.6166 | 0.36795 | 1.049 | EC0007 |
| Positive | Succinate exchange | 2 | 2 | 0.4286 | 0.5871 | 0.36795 | 1.047 | EC00044;EC00046;EC00047;EC00045;EC00015;EC00020 |
| Positive | fatty acyl-CoA synthase (n-C10:0CoA) | 3 | 3 | 0.4394 | 0.5873 | 0.35714 | 1.019 | EC00022 |
| Positive | fatty-acyl-CoA synthase (n-C12:0CoA) | 3 | 3 | 0.4394 | 0.5873 | 0.35714 | 1.019 | EC00022 |
| Positive | fatty-acyl-CoA synthase (n-C14:0CoA) | 3 | 3 | 0.4394 | 0.5873 | 0.35714 | 1.019 | EC00022;EC00034;EC0005;EC0006 |
| Positive | 5 alpha dihydrotesterone transport | 1 | 1 | 0.5306 | 0.6641 | 0.27523 | 1.017 | EC00022;EC00034;EC0005;EC0006 |
| Positive | 5alpha-Dihydrotestosterone glucuronide exchange | 1 | 1 | 0.5306 | 0.6641 | 0.27523 | 1.017 | EC00022;EC00034;EC0005;EC0006 |
| Positive | 5alpha-Dihydrotestosterone sulfate exchange | 1 | 1 | 0.5306 | 0.6641 | 0.27523 | 1.017 | EC00022;EC00034;EC0005;EC0006 |
| Positive | 5alpha-Dihydrotestosterone sulfotransferase | 1 | 1 | 0.5306 | 0.6641 | 0.27523 | 1.017 | EC00022;EC00034;EC0005;EC0006;EC00020;EC00038 |
| Positive | sulfonated testosterone transport | 1 | 1 | 0.5306 | 0.6641 | 0.27523 | 1.017 | EC00044;EC00022;EC00045;EC00034;EC00015;EC00020 |
| Positive | glucuronidated compound transport | 1 | 1 | 0.5306 | 0.6641 | 0.27523 | 1.017 | EC00048 |
| Positive | UDP-glucuronosyltransferase 1-10 precursor, microsomal | 1 | 1 | 0.5306 | 0.6641 | 0.27523 | 1.017 | EC00019;EC00050;EC0001;EC0002 |
| Positive | 3-Hydroxy-L-kynurenine hydrolase | 6 | 6 | 0.4306 | 0.5871 | 0.36593 | 1.016 | EC00045;EC00015 |
| Positive | 3-hydroxyanthranilate 3,4-dioxygenase | 6 | 6 | 0.4306 | 0.5871 | 0.36593 | 1.016 | EC00045;EC00015;EC00044 |

| | | | | | | | | |
|---|---|---|---|---|---|---|---|---|
| Positive | 5-hydroxy-L-tryptophan secretion via secretory vesicle (ATP driven) | 6 | 6 | 0.4306 | 0.5871 | 0.36593 | 1.016 | EC00046;EC00047;EC00031;EC00049 |
| Positive | kynurenine 3-monooxygenase | 6 | 6 | 0.4306 | 0.5871 | 0.36593 | 1.016 | EC00046;EC00047;EC00031;EC00049 |
| Positive | L-Tryptophan:oxygen 2,3-oxidoreductase (decyclizing) | 6 | 6 | 0.4306 | 0.5871 | 0.36593 | 1.016 | EC00046;EC00047;EC00031;EC00049 |
| Positive | N-Formyl-L-kynurenine amidohydrolase | 6 | 6 | 0.4306 | 0.5871 | 0.36593 | 1.016 | EC00046;EC00047;EC00031;EC00049;EC00025;EC00026 |
| Positive | nicotinate-nucleotide diphosphorylase (carboxylating) | 6 | 6 | 0.4306 | 0.5871 | 0.36593 | 1.016 | EC00022 |
| Positive | Quinolinate Synthase (Eukaryotic) | 6 | 6 | 0.4306 | 0.5871 | 0.36593 | 1.016 | EC00022 |
| Positive | 2-aminomuconate reductase | 6 | 6 | 0.4306 | 0.5871 | 0.36593 | 1.016 | EC00022 |
| Positive | aminomuconate-semialdehyde dehydrogenase | 6 | 6 | 0.4306 | 0.5871 | 0.36593 | 1.016 | EC00022 |
| Positive | picolinic acid decarboxylase | 6 | 6 | 0.4306 | 0.5871 | 0.36593 | 1.016 | EC00022 |
| Positive | R group artificial flux | 4 | 4 | 0.5312 | 0.6641 | 0.27474 | 0.9901 | EC00048 |
| Positive | Linoleic acid (n-C18:2) transport in via diffusion | 1 | 1 | 0.5714 | 0.6888 | 0.24306 | 0.9891 | EC00048 |
| Positive | fatty acyl-CoA desaturase (n-C18:2CoA -> n-C18:3CoA) | 1 | 1 | 0.5714 | 0.6888 | 0.24306 | 0.9891 | EC00043 |
| Positive | transport into the mitochondria (carnitine) | 1 | 1 | 0.5714 | 0.6888 | 0.24306 | 0.9891 | EC00043 |
| Positive | carnitine O-palmitoyltransferase | 1 | 1 | 0.5714 | 0.6888 | 0.24306 | 0.9891 | EC00013;EC00014;EC00028;EC00029;EC00030;EC00024;EC00027 |

| | | | | | | | | |
|---|---|---|---|---|---|---|---|---|
| Positive | Beta oxidation of fatty acid | 1 | 1 | 0.5714 | 0.6888 | 0.24306 | 0.9891 | EC00055;EC00039 |
| Positive | carnitine transferase | 1 | 1 | 0.5714 | 0.6888 | 0.24306 | 0.9891 | EC00049 |
| Negative | Proline dehydrogenase | 6 | 6 | 0.4808 | 0.6453 | 0.31804 | 0.9808 | EC0005 |
| Negative | dihydroceramide desaturase | 6 | 6 | 0.4808 | 0.6453 | 0.31804 | 0.9808 | EC0005 |
| Positive | ATP diphosphohydrolase | 6 | 6 | 0.5278 | 0.6641 | 0.27753 | 0.9668 | EC00022;EC00034;EC0005;EC0006 |
| Negative | glutamine synthetase | 2 | 2 | 0.5424 | 0.7184 | 0.26568 | 0.9524 | EC0005 |
| Positive | methionine synthase | 6 | 6 | 0.5833 | 0.6888 | 0.23411 | 0.9348 | EC00049 |
| Positive | 5,10-methylenetetrahydrofolatereductase (NADPH) | 6 | 6 | 0.5833 | 0.6888 | 0.23411 | 0.9348 | EC00049 |
| Positive | 1-acylglycerol-3-phosphate O-acyltransferase 1 | 1 | 1 | 0.6122 | 0.6888 | 0.21311 | 0.9342 | EC00049;EC00028;EC00029;EC00030 |
| Positive | glycerol-3-phosphate acyltransferase | 1 | 1 | 0.6122 | 0.6888 | 0.21311 | 0.9342 | EC00044;EC00045;EC00034;EC0005;EC0006;EC00015;EC00020 |
| Positive | R total 2 position exchange | 1 | 1 | 0.6122 | 0.6888 | 0.21311 | 0.9342 | EC00044;EC00045;EC00034;EC0005;EC0006;EC00015;EC00020 |
| Positive | FAD transporter, peroxisomal | 12 | 12 | 0.5616 | 0.6888 | 0.25057 | 0.9111 | EC00048 |
| Positive | nucleoside-triphosphatase (GTP) | 1 | 1 | 0.6122 | 0.6888 | 0.21311 | 0.9067 | EC00044;EC00045;EC00034;EC0005;EC0006;EC00015;EC00020 |

| | | | | | | | | |
|---|---|---|---|---|---|---|---|---|
| Positive | 5'-nucleotidase (GMP), extracellular | 1 | 1 | 0.6122 | 0.6888 | 0.21311 | 0.9067 | EC00045 |
| Positive | nucleoside-diphosphatase (GDP), extracellular | 1 | 1 | 0.6122 | 0.6888 | 0.21311 | 0.9067 | EC00045 |
| Positive | ADPribose diphosphatase | 1 | 1 | 0.6122 | 0.6888 | 0.21311 | 0.9067 | EC00045 |
| Positive | ADPribose transport | 1 | 1 | 0.6122 | 0.6888 | 0.21311 | 0.9067 | EC00045;EC00014;EC00028;EC00029;EC00030;EC00024;EC00027 |
| Positive | nucleoside-diphosphatase (UDP), extracellular | 1 | 1 | 0.6122 | 0.6888 | 0.21311 | 0.9067 | EC00054 |
| Positive | nucleoside-diphosphatase (UTP), extracellular | 1 | 1 | 0.6122 | 0.6888 | 0.21311 | 0.9067 | EC00054 |
| Positive | 5'-nucleotidase (UMP), extracellular | 1 | 1 | 0.6122 | 0.6888 | 0.21311 | 0.9067 | EC00054 |
| Negative | carnitine O-acetyltransferase, reverse direction, peroxisomal | 1 | 1 | 0.8033 | 0.8591 | 0.09512 | 0.9036 | EC00025;EC0003;EC00024;EC00019;EC00026 |
| Negative | carnitine-acetylcarnitine carrier, peroxisomal | 1 | 1 | 0.8033 | 0.8591 | 0.09512 | 0.9036 | EC00025;EC0003;EC00024;EC00019;EC00026;EC00021;EC00022 |
| Negative | fatty acid transport via diffusion | 1 | 1 | 0.8033 | 0.8591 | 0.09512 | 0.9036 | EC00025;EC0003;EC00024;EC00026 |
| Positive | histidine decarboxylase | 2 | 2 | 0.7321 | 0.8169 | 0.13543 | 0.8829 | EC00040 |
| Positive | glycine passive transport to mitochondria | 2 | 2 | 0.7321 | 0.8169 | 0.13543 | 0.8828 | EC00026 |
| Negative | O2 transport, peroxisomal | 1 | 1 | 0.8361 | 0.882 | 0.07774 | 0.8434 | EC0008 |

| | | | | | | | | |
|---|---|---|---|---|---|---|---|---|
| Positive | 5'-nucleotidase (AMP), extracellular | 4 | 4 | 0.8125 | 0.874 | 0.09018 | 0.8251 | EC0003;EC00056;EC00028;EC00029;EC00030;EC00042 |
| Negative | 3',5'-bisphosphate nucleotidase | 1 | 1 | 0.8852 | 0.8897 | 0.05296 | 0.7831 | EC00023;EC00021;EC00022 |
| Negative | adenylyl-sulfate kinase | 1 | 1 | 0.8852 | 0.8897 | 0.05296 | 0.7831 | EC00025;EC0007;EC0003;EC00024;EC0004;EC00019;EC00026;EC0005;EC00023;EC00021;EC00022 |
| Negative | sulfate adenylyltransferase | 1 | 1 | 0.8852 | 0.8897 | 0.05296 | 0.7831 | EC0005 |
| Positive | enolase | 3 | 3 | 0.8636 | 0.8866 | 0.06369 | 0.7821 | EC00046;EC00047;EC00031;EC00049;EC00014;EC00028;EC00029;EC00030;EC00027 |
| Positive | pyruvate kinase | 3 | 3 | 0.8636 | 0.8866 | 0.06369 | 0.7821 | EC00028;EC00029;EC00030 |
| Positive | CO2 exchange | 3 | 3 | 0.8636 | 0.8866 | 0.06369 | 0.7661 | EC00028;EC00029;EC00030 |
| Positive | Phosphatidylserine decarboxylase | 3 | 3 | 0.8636 | 0.8866 | 0.06369 | 0.7661 | EC00028;EC00029;EC00030 |
| Positive | phosphatidylserine flippase | 3 | 3 | 0.8636 | 0.8866 | 0.06369 | 0.7661 | EC00028;EC00029;EC00030 |
| Positive | triose-phosphate isomerase | 3 | 3 | 0.8636 | 0.8866 | 0.06369 | 0.7661 | EC00014;EC00028;EC00029;EC00030;EC00027 |
| Negative | pyruvate carboxylase | 2 | 2 | 0.8644 | 0.882 | 0.06329 | 0.7604 | EC00023;EC00021;EC00022 |
| Negative | Glutamate transport via Na, H symport and K antiport | 2 | 2 | 0.8644 | 0.882 | 0.06329 | 0.7604 | EC00023;EC00021;EC00022 |

| | | | | | | | | |
|---|---|---|---|---|---|---|---|---|
| Negative | citrate synthase | 2 | 2 | 0.8644 | 0.882 | 0.06329 | 0.7604 | EC00023;EC00021;EC00022 |
| Negative | enolase | 2 | 2 | 0.8644 | 0.882 | 0.06329 | 0.7604 | EC00023;EC00021;EC00022 |
| Negative | pyruvate kinase | 2 | 2 | 0.8644 | 0.882 | 0.06329 | 0.7604 | EC00023;EC00021;EC00022 |
| Negative | glyceraldehyde-3-phosphate dehydrogenase | 2 | 2 | 0.8644 | 0.882 | 0.06329 | 0.7604 | EC00023;EC00021;EC00022 |
| Positive | methylmalonyl-CoA mutase | 1 | 1 | 0.9184 | 0.9357 | 0.03697 | 0.7418 | EC00021;EC00036;EC00028;EC00029;EC00030 |
| Positive | Propionyl-CoA carboxylase, mitochondrial | 1 | 1 | 0.9184 | 0.9357 | 0.03697 | 0.7418 | EC00032 |
| Positive | xenobiotic transport | 4 | 4 | 0.9375 | 0.9375 | 0.02803 | 0.7248 | EC00054 |
| Negative | IMP dehydrogenase | 2 | 2 | 0.8983 | 0.8983 | 0.04658 | 0.7123 | EC0005 |
| Positive | pyruvate carboxylase | 2 | 2 | 0.9362 | 0.9375 | 0.02863 | -0.667 | EC00012;EC00046;EC00047;EC0003;EC00022;EC00031;EC00045;EC00056;EC00055;EC00018;EC0001;EC0002;EC00034;EC00015;EC00020;EC00044;EC00026;EC00037 |
| Positive | citrate synthase | 2 | 2 | 0.9362 | 0.9375 | 0.02863 | -0.667 | EC00054 |
| Positive | glyceraldehyde-3-phosphate dehydrogenase | 7 | 7 | 0.8519 | 0.8866 | 0.06961 | -0.6706 | EC0003;EC00056;EC00028;EC00029;EC00030;EC00042 |

| | | | | | | | | |
|---|---|---|---|---|---|---|---|---|
| Positive | R group coenzyme a ligase | 2 | 2 | 0.9362 | 0.9375 | 0.02863 | -0.6809 | EC00026 |
| Positive | ATP synthase (four protons for one ATP) | 6 | 6 | 0.9333 | 0.9375 | 0.02998 | -0.7067 | EC00032 |
| Positive | 3',5'-bisphosphate nucleotidase | 3 | 3 | 0.8611 | 0.8866 | 0.06495 | -0.7271 | EC0003;EC00056;EC00028;EC00029;EC00030;EC00042 |
| Positive | adenylyl-sulfate kinase | 3 | 3 | 0.8611 | 0.8866 | 0.06495 | -0.7271 | EC0003;EC00056;EC00028;EC00029;EC00030;EC00042 |
| Positive | sulfate adenylyltransferase | 3 | 3 | 0.8611 | 0.8866 | 0.06495 | -0.7271 | EC00033;EC00028;EC00029;EC00030;EC00051 |
| Positive | cystathionine beta-synthase | 1 | 1 | 0.8491 | 0.8866 | 0.07104 | -0.7386 | EC0003;EC00056;EC00028;EC00029;EC00030;EC00042 |
| Positive | cystathionine g-lyase | 1 | 1 | 0.8491 | 0.8866 | 0.07104 | -0.7386 | EC0003;EC00056;EC00028;EC00029;EC00030;EC00042 |
| Positive | glycine hydroxymethyltransferase, reversible | 7 | 7 | 0.7778 | 0.84 | 0.10913 | -0.7388 | EC0003;EC00056;EC00028;EC00029;EC00030;EC00042 |
| Positive | phosphoglycerate dehydrogenase | 7 | 7 | 0.7778 | 0.84 | 0.10913 | -0.7388 | EC0003;EC00056;EC00028;EC00029;EC00030;EC00042 |

| | | | | | | | | |
|---|---|---|---|---|---|---|---|---|
| Positive | phosphoserine phosphatase (L-serine) | 7 | 7 | 0.7778 | 0.84 | 0.10913 | -0.7388 | EC0003;EC00056;EC00028;EC00029;EC00030;EC00042 |
| Positive | phosphoserine transaminase | 7 | 7 | 0.7778 | 0.84 | 0.10913 | -0.7388 | EC0003;EC00056;EC00028;EC00029;EC00030;EC00042 |
| Negative | Tetrahydrobiopterin-4a-carbinolamine dehydratase | 3 | 3 | 0.85 | 0.882 | 0.07058 | -0.75 | EC0008 |
| Negative | Iodide:hydrogen-peroxide oxidoreductase 3 | 2 | 2 | 0.7674 | 0.8591 | 0.11498 | -0.7885 | EC00025;EC0003;EC00024 |
| Negative | Triiodothyronine exchange | 2 | 2 | 0.7674 | 0.8591 | 0.11498 | -0.7885 | EC00025;EC00011;EC0003;EC00024;EC00026;EC00015 |
| Positive | Iodide:hydrogen-peroxide oxidoreductase | 1 | 1 | 0.7736 | 0.84 | 0.11148 | -0.7933 | EC00040 |
| Positive | Iodide:hydrogen-peroxide oxidoreductase 2 | 1 | 1 | 0.7736 | 0.84 | 0.11148 | -0.7933 | EC00028;EC00029;EC00030;EC00025;EC00043 |
| Positive | L-Tyrosine carboxy-lyase | 1 | 1 | 0.7736 | 0.84 | 0.11148 | -0.7933 | EC00028;EC00029;EC00030;EC00025;EC00043 |
| Positive | Tyrosine:dopa oxidase | 1 | 1 | 0.7736 | 0.84 | 0.11148 | -0.7933 | EC0003;EC00056;EC00028;EC00029;EC00030;EC00042 |
| Negative | L-Phenylalanine,tetrahydrobiopterin:oxygen oxidoreductase (4-hydroxylating) | 4 | 4 | 0.75 | 0.8591 | 0.12494 | -0.8318 | EC00025;EC0003;EC00024 |

| | | | | | | | | |
|---|---|---|---|---|---|---|---|---|
| Negative | xenobiotic transport | 1 | 1 | 0.8542 | 0.882 | 0.06844 | -0.8545 | EC00023;EC00021;EC00022 |
| Negative | nucleoside-triphosphatase (GTP) | 2 | 2 | 0.6744 | 0.8225 | 0.17108 | -0.8617 | EC0005 |
| Negative | 5'-nucleotidase (GMP), extracellular | 2 | 2 | 0.6744 | 0.8225 | 0.17108 | -0.8617 | EC00023;EC00021;EC00022 |
| Negative | nucleoside-diphosphatase (GDP), extracellular | 2 | 2 | 0.6744 | 0.8225 | 0.17108 | -0.8617 | EC00025;EC0003;EC00024 |
| Positive | 5-Hydroxy-L-tryptophan decarboxy-lyase | 3 | 3 | 0.6111 | 0.6888 | 0.21389 | -0.864 | EC00049 |
| Positive | 5-Hydroxy-L-tryptophan exchange | 3 | 3 | 0.6111 | 0.6888 | 0.21389 | -0.864 | EC00049 |
| Positive | L-Tryptophan,tetrahydrobiopterin:oxygen oxidoreductase (5-hydroxylating) | 3 | 3 | 0.6111 | 0.6888 | 0.21389 | -0.864 | EC00049 |
| Positive | Serotonin exchange | 3 | 3 | 0.6111 | 0.6888 | 0.21389 | -0.864 | EC00049;EC00028;EC00029;EC00030 |
| Negative | R group coenzyme a ligase | 1 | 1 | 0.7917 | 0.8591 | 0.10144 | -0.9203 | EC00025;EC00011;EC0003;EC00024;EC00026;EC00015 |
| Negative | GMP synthase | 1 | 1 | 0.7917 | 0.8591 | 0.10144 | -0.9203 | EC00025;EC0003;EC00024;EC00019;EC00014;EC00026 |

| | | | | | | | | |
|---|---|---|---|---|---|---|---|---|
| Negative | glutamine phosphoribosyldiphosphate amidotransferase | 1 | 1 | 0.7917 | 0.8591 | 0.10144 | -0.9203 | EC00025;EC0003;EC00024;EC00019;EC00014;EC00026 |
| Negative | phosphoribosylaminoimidazole carboxylase | 1 | 1 | 0.7917 | 0.8591 | 0.10144 | -0.9203 | EC00025;EC0003;EC00024;EC00019;EC00014;EC00026 |
| Negative | phosphoribosylaminoimidazole synthase | 1 | 1 | 0.7917 | 0.8591 | 0.10144 | -0.9203 | EC00025;EC0003;EC00024;EC00019;EC00014;EC00026 |
| Negative | phosphoribosylaminoimidazolecarboxamide formyltransferase | 1 | 1 | 0.7917 | 0.8591 | 0.10144 | -0.9203 | EC00025;EC0003;EC00024;EC00019;EC00014;EC00026 |
| Negative | phosphoribosylaminoimidazolesuccinocarboxamide synthase | 1 | 1 | 0.7917 | 0.8591 | 0.10144 | -0.9203 | EC00025;EC0003;EC00024;EC00019;EC00014;EC00026 |
| Negative | phosphoribosylformylglycinamidine synthase | 1 | 1 | 0.7917 | 0.8591 | 0.10144 | -0.9203 | EC00025;EC0003;EC00024;EC00019;EC00014;EC00026 |
| Negative | phosphoribosylglycinamide formyltransferase | 1 | 1 | 0.7917 | 0.8591 | 0.10144 | -0.9203 | EC00025;EC0003;EC00024;EC00019;EC00014;EC00026 |
| Negative | phosphoribosylglycinamide synthase | 1 | 1 | 0.7917 | 0.8591 | 0.10144 | -0.9203 | EC00025;EC0003;EC00024;EC00019;EC00014;EC00026 |
| Negative | adenylosuccinate lyase | 1 | 1 | 0.7917 | 0.8591 | 0.10144 | -0.9203 | EC00025;EC0003;EC00024;EC00019;EC00014;EC00026 |

| | | | | | | | | |
|---|---|---|---|---|---|---|---|---|
| Negative | phosphoribosylpyrophosphate synthetase | 1 | 1 | 0.7917 | 0.8591 | 0.10144 | -0.9203 | EC00025;EC0003;EC00024;EC00019;EC00014;EC00026 |
| Negative | ADPribose diphosphatase | 1 | 1 | 0.7917 | 0.8591 | 0.10144 | -0.9203 | EC00025;EC0009;EC0003;EC00024;EC00026;EC00017 |
| Negative | ADPribose transport | 1 | 1 | 0.7917 | 0.8591 | 0.10144 | -0.9203 | EC00025;EC0007;EC0003;EC00024;EC00019;EC00026;EC00021;EC00022 |
| Negative | nucleoside-diphosphatase (UDP), extracellular | 1 | 1 | 0.7917 | 0.8591 | 0.10144 | -0.9203 | EC00025;EC0003;EC00024;EC00026 |
| Negative | nucleoside-diphosphatase (UTP), extracellular | 1 | 1 | 0.7917 | 0.8591 | 0.10144 | -0.9203 | EC00025;EC0003;EC00024;EC00019;EC00026 |
| Negative | 5'-nucleotidase (UMP), extracellular | 1 | 1 | 0.7917 | 0.8591 | 0.10144 | -0.9203 | EC00025;EC0003;EC00024;EC00019;EC00026 |
| Positive | L-Fucose exchange | 2 | 2 | 0.4894 | 0.6399 | 0.31034 | -0.9829 | EC00022;EC00034;EC0005;EC0006 |
| Positive | alpha-fucosidase, extracellular | 2 | 2 | 0.4894 | 0.6399 | 0.31034 | -0.9829 | EC00022;EC00034;EC0005;EC0006 |
| Positive | dihydrofolate reductase | 1 | 1 | 0.4906 | 0.6399 | 0.30927 | -1.012 | EC00022;EC00034;EC0005;EC0006 |
| Positive | folate reductase | 1 | 1 | 0.4906 | 0.6399 | 0.30927 | -1.012 | EC00022;EC00034;EC0005;EC0006 |

| | | | | | | | | |
|---|---|---|---|---|---|---|---|---|
| Positive | folate transport via anion exchange | 1 | 1 | 0.4906 | 0.6399 | 0.30927 | -1.012 | EC00022;EC00034;EC0005;EC0006 |
| Negative | FAD transporter, peroxisomal | 5 | 5 | 0.5106 | 0.6809 | 0.29192 | -1.017 | EC0005 |
| Positive | Iodide:hydrogen-peroxide oxidoreductase 3 | 2 | 2 | 0.4043 | 0.5871 | 0.3933 | -1.018 | EC00013 |
| Positive | Triiodothyronine exchange | 2 | 2 | 0.4043 | 0.5871 | 0.3933 | -1.018 | EC00013 |
| Positive | Glutamate transport via Na, H symport and K antiport | 6 | 6 | 0.3667 | 0.5871 | 0.43569 | -1.042 | EC00022;EC00037 |
| Positive | glutamine synthetase | 6 | 6 | 0.3667 | 0.5871 | 0.43569 | -1.042 | EC00037;EC00022 |
| Negative | adenosylhomocysteinase | 1 | 1 | 0.6458 | 0.8023 | 0.1899 | -1.052 | EC0006 |
| Negative | methionine adenosyltransferase | 1 | 1 | 0.6458 | 0.8023 | 0.1899 | -1.052 | EC00025;EC0003;EC00024;EC00026 |
| Negative | Adrenaline secretion via secretory vesicle (ATP driven) | 1 | 1 | 0.6458 | 0.8023 | 0.1899 | -1.052 | EC00025;EC0003;EC00024;EC00026 |
| Negative | dopamine beta-monooxygenase | 1 | 1 | 0.6458 | 0.8023 | 0.1899 | -1.052 | EC00024;EC00019;EC00026;EC0006;EC00021;EC00022 |
| Negative | metanephrine secretion via secretory vesicle (ATP driven) | 1 | 1 | 0.6458 | 0.8023 | 0.1899 | -1.052 | EC0005 |
| Negative | 3-Hydroxy-L-tyrosine carboxy-lyase | 1 | 1 | 0.6458 | 0.8023 | 0.1899 | -1.052 | EC0005 |
| Positive | H2O transport, lysosomal | 1 | 1 | 0.4151 | 0.5871 | 0.38185 | -1.067 | EC00044;EC00018;EC00038 |

| | | | | | | | | |
|---|---|---|---|---|---|---|---|---|
| Positive | endo-beta-N-acetylglucosaminidase, lysosomal | 1 | 1 | 0.4151 | 0.5871 | 0.38185 | -1.067 | EC00044;EC00018;EC00038 |
| Positive | glycosylasparaginase, lysosomal | 1 | 1 | 0.4151 | 0.5871 | 0.38185 | -1.067 | EC00044;EC00045;EC00015 |
| Positive | beta-N-acetylhexosaminidase, lysosomal | 1 | 1 | 0.4151 | 0.5871 | 0.38185 | -1.067 | EC00044;EC00045;EC00052 |
| Positive | N-acetylglucosamine-6-phosphate deacetylase | 1 | 1 | 0.4151 | 0.5871 | 0.38185 | -1.067 | EC00044;EC00045;EC00052 |
| Positive | glucosamine-6-phosphate deaminase | 1 | 1 | 0.4151 | 0.5871 | 0.38185 | -1.067 | EC00044;EC00045;EC00052 |
| Positive | alpha-mannosidase, lysosomal | 1 | 1 | 0.4151 | 0.5871 | 0.38185 | -1.067 | EC00044;EC00045;EC00015 |
| Positive | beta-mannosidase, lysosomal | 1 | 1 | 0.4151 | 0.5871 | 0.38185 | -1.067 | EC00044;EC00045;EC00015 |
| Positive | DM Asn-X-Ser/Thr(ly) | 1 | 1 | 0.4151 | 0.5871 | 0.38185 | -1.067 | EC00044;EC00045;EC00018;EC00034;EC00015;EC00020;EC00038 |
| Positive | mannose efflux from lysosome | 1 | 1 | 0.4151 | 0.5871 | 0.38185 | -1.067 | EC00044;EC00046;EC00047;EC00031;EC00045;EC00055;EC00021;EC00020 |
| Positive | N-acetyl-glucosamine lysosomal efflux | 1 | 1 | 0.4151 | 0.5871 | 0.38185 | -1.067 | EC00044;EC00046;EC00047;EC00031;EC00045;EC00055;EC00021;EC00020 |
| Positive | N-acetylglucosamine kinase | 1 | 1 | 0.4151 | 0.5871 | 0.38185 | -1.067 | EC00044;EC00046;EC00047;EC00045;EC00015;EC00020 |
| Negative | 5-Hydroxy-L-tryptophan decarboxy-lyase | 1 | 1 | 0.5625 | 0.7258 | 0.24988 | -1.117 | EC00012;EC0005 |

| | | | | | | | | |
|---|---|---|---|---|---|---|---|---|
| Negative | 5-Hydroxy-L-tryptophan exchange | 1 | 1 | 0.5625 | 0.7258 | 0.24988 | -1.117 | EC00016 |
| Negative | L-Tryptophan,tetrahydrobiopterin:oxygen oxidoreductase (5-hydroxylating) | 1 | 1 | 0.5625 | 0.7258 | 0.24988 | -1.117 | EC00015 |
| Negative | Serotonin exchange | 1 | 1 | 0.5625 | 0.7258 | 0.24988 | -1.117 | EC00015 |
| Positive | pyridoxamine 5'-phosphate oxidase | 1 | 1 | 0.3962 | 0.5871 | 0.40209 | -1.122 | EC00046;EC00047;EC0003;EC00031;EC00056;EC00049;EC00025;EC00026;EC00043 |
| Positive | pyridoxamine kinase | 1 | 1 | 0.3962 | 0.5871 | 0.40209 | -1.122 | EC00046;EC00047;EC00031;EC00049;EC00025;EC00026 |
| Positive | pyridoxamine transport via diffusion | 1 | 1 | 0.3962 | 0.5871 | 0.40209 | -1.122 | EC00046;EC00047;EC00031;EC00041;EC00049;EC00025;EC00026;EC00043 |
| Positive | pyridoxine 5'-phosphate oxidase | 1 | 1 | 0.3962 | 0.5871 | 0.40209 | -1.122 | EC00025;EC00043 |
| Positive | pyridoxine kinase | 1 | 1 | 0.3962 | 0.5871 | 0.40209 | -1.122 | EC00025;EC00043 |
| Positive | pyridoxine transport via diffusion | 1 | 1 | 0.3962 | 0.5871 | 0.40209 | -1.122 | EC00018;EC00038 |
| Negative | 3,4-Dihydroxy-L-phenylalanine transport | 2 | 2 | 0.3023 | 0.4547 | 0.51956 | -1.159 | EC00015 |
| Negative | sterol 12-alpha-hydroxylase | 7 | 7 | 0.2041 | 0.3644 | 0.69016 | -1.172 | EC00025;EC0007;EC0003;EC00024;EC00019;EC00026;EC0006;EC00021;EC00022 |

| | | | | | | | | |
|---|---|---|---|---|---|---|---|---|
| Negative | sterol 12-alpha-hydroxylase (nadh) | 7 | 7 | 0.2041 | 0.3644 | 0.69016 | -1.172 | EC0006;EC00025;EC0003;EC00024;EC00019;EC00026;EC00021;EC00022 |
| Positive | ADPribose 2'-phosphate exchange | 1 | 1 | 0.3208 | 0.5871 | 0.49377 | -1.176 | EC00014;EC00024;EC00027 |
| Positive | 1-Methylnicotinamide exchange | 1 | 1 | 0.3208 | 0.5871 | 0.49377 | -1.176 | EC00014;EC00024;EC00027 |
| Positive | N1-Methylnicotinamide transport | 1 | 1 | 0.3208 | 0.5871 | 0.49377 | -1.176 | EC00046;EC00047;EC00031;EC00049;EC00014;EC00028;EC00029;EC00030;EC00024;EC00026;EC00027 |
| Positive | Nicotinamide N-methyltransferase | 1 | 1 | 0.3208 | 0.5871 | 0.49377 | -1.176 | EC00046;EC00047;EC00031;EC00049;EC00014;EC00028;EC00029;EC00030;EC00024;EC00026;EC00027 |
| Positive | Trehalose exchange | 18 | 18 | 0.2727 | 0.5871 | 0.56431 | -1.189 | EC00014;EC00028;EC00029;EC00030;EC0004;EC00016;EC00017;EC00024;EC00027;EC00044;EC00046;EC00047;EC0003;EC00031;EC00056;EC00055;EC0001;EC0002;EC00026;EC00037 |
| Positive | methenyltetrahydrifikate cyclohydrolase, mitochondrial | 1 | 1 | 0.3019 | 0.5871 | 0.52014 | -1.204 | EC00014;EC00028;EC00029;EC00030;EC0004;EC00016;EC00017;EC00024;EC00027 |
| Positive | 3-Dehydrosphinganine reductase | 1 | 1 | 0.3019 | 0.5871 | 0.52014 | -1.204 | EC00014;EC00028;EC00029;EC00030;EC0004;EC00016;EC00017;EC00024;EC00027 |
| Positive | dihydrosphingosine N-acyltransferase | 1 | 1 | 0.3019 | 0.5871 | 0.52014 | -1.204 | EC00014;EC00028;EC00029;EC00030;EC0004;EC00016;EC00017;EC00024;EC00027 |

| | | | | | | | | |
|---|---|---|---|---|---|---|---|---|
| Positive | serine C-palmitoyltransferase | 1 | 1 | 0.3019 | 0.5871 | 0.52014 | -1.204 | EC00014;EC00028;EC00029;EC00030;EC0004;EC00016;EC00017;EC00024;EC00027 |
| Positive | adenylosuccinate synthase | 2 | 2 | 0.2766 | 0.5871 | 0.55815 | -1.229 | EC00014;EC00028;EC00029;EC00030;EC0004;EC00016;EC00017;EC00024;EC00027 |
| Positive | methenyltetrahydrofolate cyclohydrolase | 1 | 1 | 0.2264 | 0.5316 | 0.64512 | -1.258 | EC00014;EC00028;EC00029;EC00030;EC0004;EC00016;EC00017;EC00024;EC00027 |
| Negative | dihydrofolate reductase | 1 | 1 | 0.25 | 0.4031 | 0.60206 | -1.315 | EC00010;EC0005 |
| Negative | folate reductase | 1 | 1 | 0.25 | 0.4031 | 0.60206 | -1.315 | EC00010;EC0005 |
| Negative | folate transport via anion exchange | 1 | 1 | 0.25 | 0.4031 | 0.60206 | -1.315 | EC00010;EC0005 |
| Positive | cytochrome c oxidase, mitochondrial Complex IV | 1 | 1 | 0.1132 | 0.3026 | 0.94615 | -1.34 | EC00046;EC00047;EC00031;EC00049;EC00014;EC00028;EC00029;EC00030;EC0004;EC00016;EC00017;EC00024;EC00026;EC00027 |
| Positive | sulfite oxidase | 1 | 1 | 0.1132 | 0.3026 | 0.94615 | -1.34 | EC00046;EC00047;EC00031;EC00049;EC00014;EC00028;EC00029;EC00030;EC0004;EC00016;EC00017;EC00024;EC00026;EC00027 |
| Positive | GMP reductase | 3 | 3 | 0.1667 | 0.4054 | 0.77806 | -1.351 | EC00014;EC00028;EC00029;EC00030;EC0004;EC00016;EC00017;EC00024;EC00027 |
| Positive | guanine phosphoribosyltransferase | 3 | 3 | 0.1667 | 0.4054 | 0.77806 | -1.351 | EC00014;EC00028;EC00029;EC00030;EC0004;EC00016;EC00017;EC00024;EC00027 |

| | | | | | | | | |
|---|---|---|---|---|---|---|---|---|
| Negative | Trehalose exchange | 11 | 11 | 0.1579 | 0.2871 | 0.80162 | -1.371 | EC00025;EC0003;EC00024;EC00019;EC00026;EC0006;EC00021;EC00022 |
| Negative | lipid, flip-flop intracellular transport | 11 | 11 | 0.1404 | 0.2575 | 0.85263 | -1.386 | EC00025;EC0003;EC00024;EC00019;EC00026;EC0006;EC00021;EC00022 |
| Negative | peroxisomal thiolase 2 | 9 | 9 | 0.1034 | 0.1916 | 0.98548 | -1.43 | EC00025;EC0003;EC00024;EC00019;EC00026;EC0006;EC00023;EC00021;EC00022 |
| Negative | bile acid intracellular transport | 9 | 9 | 0.1034 | 0.1916 | 0.98548 | -1.43 | EC00025;EC0003;EC00024;EC00019;EC00026;EC0006;EC00023;EC00021;EC00022 |
| Negative | carnitine O-aceyltransferase, mitochondrial | 9 | 9 | 0.1034 | 0.1916 | 0.98548 | -1.43 | EC00026 |
| Negative | carnitine O-aceyltransferase, peroxisomal | 9 | 9 | 0.1034 | 0.1916 | 0.98548 | -1.43 | EC00025;EC0007;EC0003;EC00024;EC00019;EC00026;EC0006;EC00021;EC00022 |
| Negative | carnitine-propcarnitine carrier, peroxisomal | 9 | 9 | 0.1034 | 0.1916 | 0.98548 | -1.43 | EC00025;EC0007;EC0003;EC00024;EC00019;EC00026;EC0006;EC00021;EC00022 |
| Negative | cytochrome P450, family 7, subfamily A, polypeptide 1 | 9 | 9 | 0.1034 | 0.1916 | 0.98548 | -1.43 | EC00025;EC0007;EC0003;EC00024;EC00019;EC00026;EC0006;EC00021;EC00022 |
| Negative | taurine transport (sodium symport) (cytosol to peroxisome) | 9 | 9 | 0.1034 | 0.1916 | 0.98548 | -1.43 | EC00025;EC0003;EC00024;EC00019;EC00026;EC0006;EC00021;EC00022 |
| Negative | taurochenodeoxycholate exchange | 9 | 9 | 0.1034 | 0.1916 | 0.98548 | -1.43 | EC00025;EC0003;EC00024;EC00019;EC00026;EC0006;EC00021;EC00022 |

| | | | | | | | | |
|---|---|---|---|---|---|---|---|---|
| Negative | transport into the mitochondria from cytosol (carnitine) | 9 | 9 | 0.1034 | 0.1916 | 0.98548 | -1.43 | EC00025;EC0003;EC00024;EC00019;EC00026;EC0006;EC00021;EC00022 |
| Negative | Cytochrome P450 27 | 9 | 9 | 0.1034 | 0.1916 | 0.98548 | -1.43 | EC00025;EC0003;EC00024;EC00019;EC00026;EC0006;EC00021;EC00022 |
| Negative | aldo-keto reductase family 1, member C4 (chlordecone reductase; 3-alpha hydroxysteroid dehydrogenase, type I; dihydrodiol dehydrogenase 4) | 9 | 9 | 0.1034 | 0.1916 | 0.98548 | -1.43 | EC00025;EC0003;EC00024;EC00019;EC00026;EC0006;EC00021;EC00022 |
| Negative | aldo-keto reductase family 1, member D1 (delta 4-3-ketosteroid-5-beta-reductase) | 9 | 9 | 0.1034 | 0.1916 | 0.98548 | -1.43 | EC00025;EC0003;EC00024;EC00019;EC00026;EC0006;EC00021;EC00022 |
| Negative | Very-long-chain-fatty-acid-CoA ligase | 9 | 9 | 0.1034 | 0.1916 | 0.98548 | -1.43 | EC00025;EC0003;EC00024;EC00019;EC00026;EC0006;EC00021;EC00022 |
| Negative | 5-beta-cytochrome P450, family 27, subfamily A, polypeptide 1 | 9 | 9 | 0.1034 | 0.1916 | 0.98548 | -1.43 | EC00025;EC0003;EC00024;EC00019;EC00026;EC0006;EC00021;EC00022 |
| Negative | bile acid Coenzyme A: amino acid N-acyltransferase | 9 | 9 | 0.1034 | 0.1916 | 0.98548 | -1.43 | EC00025;EC0003;EC00024;EC00019;EC00026;EC0006;EC00021;EC00022 |
| Negative | protoporphyrinogen oxidase, mitochondrial | 9 | 9 | 0.1034 | 0.1916 | 0.98548 | -1.479 | EC00025;EC0003;EC00024;EC00019;EC00026;EC0006;EC00023;EC00021;EC00022 |
| Negative | C-14 sterol reductase | 9 | 9 | 0.1034 | 0.1916 | 0.98548 | -1.479 | EC00025;EC0003;EC00024;EC00019;EC00026;EC0006;EC00023;EC00021;EC00022 |

| | | | | | | | | |
|---|---|---|---|---|---|---|---|---|
| Negative | C-3 sterol keto reductase (zymosterol) | 9 | 9 | 0.1034 | 0.1916 | 0.98548 | -1.479 | EC00025;EC0003;EC00024;EC00019;EC00026;EC0006;EC00023;EC00021;EC00022 |
| Negative | C-4 sterol methyl oxidase (4,4-dimethylzymosterol) | 9 | 9 | 0.1034 | 0.1916 | 0.98548 | -1.479 | EC00025;EC0003;EC00024;EC00019;EC00026;EC0006;EC00023;EC00021;EC00022;EC0007;EC0005 |
| Negative | cholesterol precursor intracellular transport | 9 | 9 | 0.1034 | 0.1916 | 0.98548 | -1.479 | EC00025;EC0003;EC00024;EC00019;EC00026;EC0006;EC00021;EC00022 |
| Negative | cytochrome P450 lanosterol 14-alpha-demethylase | 9 | 9 | 0.1034 | 0.1916 | 0.98548 | -1.479 | EC00025;EC0003;EC00024;EC00019;EC00026;EC0006;EC00021;EC00022;EC00023 |
| Negative | FAD transporter, endoplasmic reticulum | 9 | 9 | 0.1034 | 0.1916 | 0.98548 | -1.479 | EC0003;EC00024;EC00019;EC00026;EC0006;EC00021;EC00022 |
| Negative | FADH2 transporter, endoplasmic reticulum | 9 | 9 | 0.1034 | 0.1916 | 0.98548 | -1.479 | EC0003;EC00024;EC00019;EC00026;EC0006;EC00021;EC00022 |
| Negative | lanosterol synthase | 9 | 9 | 0.1034 | 0.1916 | 0.98548 | -1.479 | EC00019;EC00026;EC0006;EC00023;EC00021;EC00022 |
| Negative | Previtamin D3 formation | 9 | 9 | 0.1034 | 0.1916 | 0.98548 | -1.479 | EC0003;EC00019;EC00026;EC0006;EC00023;EC00021;EC00022 |
| Negative | Squalene epoxidase, endoplasmic reticular (NADP) | 9 | 9 | 0.1034 | 0.1916 | 0.98548 | -1.479 | EC0006;EC00018 |
| Negative | Squalene synthase | 9 | 9 | 0.1034 | 0.1916 | 0.98548 | -1.479 | EC00024;EC00019;EC00026;EC0006;EC00021;EC00022 |
| Negative | Vitamin D3 exchange | 9 | 9 | 0.1034 | 0.1916 | 0.98548 | -1.479 | EC0006 |

| | | | | | | | | |
|---|---|---|---|---|---|---|---|---|
| Negative | Vitamin D3 formation | 9 | 9 | 0.1034 | 0.1916 | 0.98548 | -1.479 | EC0006 |
| Negative | 3-beta-hydroxysteroid-delta(8),delta(7)-isomerase | 9 | 9 | 0.1034 | 0.1916 | 0.98548 | -1.479 | EC00025;EC0003;EC00024;EC00019;EC00026;EC0006;EC00021;EC00022 |
| Negative | C-3 sterol dehydrogenase (4-methylzymosterol) | 9 | 9 | 0.1034 | 0.1916 | 0.98548 | -1.479 | EC00025;EC0003;EC00024;EC00019;EC00026;EC0006;EC00021;EC00022 |
| Negative | C-4 methyl sterol oxidase | 9 | 9 | 0.1034 | 0.1916 | 0.98548 | -1.479 | EC00025;EC0003;EC00024;EC00019;EC00026;EC0006;EC00021;EC00022 |
| Negative | 7-dehydrocholesterol reductase | 9 | 9 | 0.1034 | 0.1916 | 0.98548 | -1.479 | EC00025;EC0003;EC00024;EC00019;EC00026;EC0006;EC00021;EC00022 |
| Negative | 24-dehydrocholesterol reductase [Precursor] | 9 | 9 | 0.1034 | 0.1916 | 0.98548 | -1.479 | EC00025;EC0007;EC0003;EC00024;EC00019;EC00026;EC0006;EC00021;EC00022 |
| Negative | Lathosterol oxidase | 9 | 9 | 0.1034 | 0.1916 | 0.98548 | -1.479 | EC00025;EC0007;EC0003;EC00024;EC00019;EC00026;EC0006;EC00021;EC00022 |
| Negative | Beta oxidation of long chain fatty acid | 9 | 9 | 0.1034 | 0.1916 | 0.98548 | -1.479 | EC00025;EC0007;EC0003;EC00024;EC00019;EC00026;EC0006;EC00021;EC00022 |
| Negative | alpha-methylacyl-CoA racemase | 8 | 8 | 0.09615 | 0.1916 | 1.01705 | -1.488 | EC00025;EC0003;EC00024;EC00019;EC00026;EC0006;EC00021;EC00022 |
| Negative | alpha-methylacyl-CoA racemase (reductase) | 8 | 8 | 0.09615 | 0.1916 | 1.01705 | -1.488 | EC00025;EC0003;EC00024;EC00019;EC00026;EC0006;EC00021;EC00022 |

| | | | | | | | | |
|---|---|---|---|---|---|---|---|---|
| Negative | Chenodeoxyglycocholate exchange | 8 | 8 | 0.09615 | 0.1916 | 1.01705 | -1.488 | EC00025;EC0003;EC00024;EC00019;EC00026;EC0006;EC00021;EC00022 |
| Negative | FADH2 transporter, peroxisomal | 8 | 8 | 0.09615 | 0.1916 | 1.01705 | -1.488 | EC00025;EC0003;EC00024;EC00019;EC00026;EC0006;EC00021;EC00022 |
| Negative | glycocholate exchange | 8 | 8 | 0.09615 | 0.1916 | 1.01705 | -1.488 | EC00025;EC0003;EC00024;EC00019;EC00026;EC0006;EC00021;EC00022 |
| Negative | Taurocholic acid exchange | 8 | 8 | 0.09615 | 0.1916 | 1.01705 | -1.488 | EC00025;EC0003;EC00024;EC00019;EC00026;EC0006;EC00021;EC00022;EC00023 |
| Negative | acyl-Coenzyme A oxidase 2, branched chain | 8 | 8 | 0.09615 | 0.1916 | 1.01705 | -1.488 | EC00025;EC0003;EC00024;EC00019;EC00026;EC0006;EC00021;EC00022;EC00023 |
| Negative | hydroxysteroid (17-beta) dehydrogenase 4 | 8 | 8 | 0.09615 | 0.1916 | 1.01705 | -1.488 | EC00025;EC0007;EC0003;EC00024;EC00019;EC00026;EC0006;EC00021;EC00022 |
| Negative | 5-beta-cholestane-3-alpha,7-alpha,12-alpha-triol 27-hydroxylase | 8 | 8 | 0.09615 | 0.1916 | 1.01705 | -1.488 | EC00025;EC0007;EC0003;EC00024;EC00019;EC00026;EC0006;EC00021;EC00022 |
| Negative | dimethylallyltranstransferase | 8 | 8 | 0.09615 | 0.1916 | 1.01705 | -1.488 | EC00025;EC0007;EC0003;EC00024;EC00019;EC00026;EC0006;EC00021;EC00022 |
| Negative | geranyltranstransferase | 8 | 8 | 0.09615 | 0.1916 | 1.01705 | -1.488 | EC00025;EC0007;EC0003;EC00024;EC00019;EC00026;EC0006;EC00021;EC00022 |

| | | | | | | | | |
|---|---|---|---|---|---|---|---|---|
| Negative | isopentenyl-diphosphate D-isomerase | 8 | 8 | 0.09615 | 0.1916 | 1.01705 | -1.488 | EC00025;EC0007;EC0003;EC00024;EC00019;EC00026;EC0006;EC00021;EC00022 |
| Negative | glycochenodeoxycholate exchange | 8 | 8 | 0.09615 | 0.1916 | 1.01705 | -1.488 | EC00025;EC0007;EC0003;EC00024;EC00019;EC00026;EC0006;EC00021;EC00022 |
| Negative | Carbon monoxide exchange | 8 | 8 | 0.09615 | 0.1916 | 1.01705 | -1.488 | EC00025;EC0007;EC0003;EC00024;EC00019;EC00026;EC0006;EC00021;EC00022 |
| Negative | CO transporter via diffusion | 8 | 8 | 0.09615 | 0.1916 | 1.01705 | -1.488 | EC00025;EC0007;EC0003;EC00024;EC00019;EC00026;EC0006;EC00021;EC00022 |
| Negative | coproporphyrinogen oxidase (O2 required) | 8 | 8 | 0.09615 | 0.1916 | 1.01705 | -1.488 | EC00025;EC0007;EC0003;EC00024;EC00019;EC00026;EC0006;EC00021;EC00022 |
| Negative | Ferrochelatase, mitochondrial | 8 | 8 | 0.09615 | 0.1916 | 1.01705 | -1.488 | EC00025;EC0007;EC0003;EC00024;EC00019;EC00026;EC0006;EC00021;EC00022 |
| Negative | Heme oxygenase 1 | 8 | 8 | 0.09615 | 0.1916 | 1.01705 | -1.488 | EC00025;EC0007;EC0003;EC00024;EC00019;EC00026;EC0006;EC00021;EC00022 |
| Negative | Heme transport to cytosol | 8 | 8 | 0.09615 | 0.1916 | 1.01705 | -1.488 | EC00025;EC0007;EC0003;EC00024;EC00019;EC00026;EC0006;EC00021;EC00022 |
| Negative | hydroxymethylbilane synthase | 8 | 8 | 0.09615 | 0.1916 | 1.01705 | -1.488 | EC00025;EC0007;EC0003;EC00024;EC00019;EC00026;EC0006;EC00021;EC00022 |

| | | | | | | | | |
|---|---|---|---|---|---|---|---|---|
| Negative | iron (II) transport | 8 | 8 | 0.09615 | 0.1916 | 1.01705 | -1.488 | EC00025;EC0003;EC00024;EC00019;EC00026;EC0006;EC00021;EC00022 |
| Negative | Nad(p)h biliverdin reductase | 8 | 8 | 0.09615 | 0.1916 | 1.01705 | -1.488 | EC00025;EC0003;EC00024;EC00019;EC00026;EC0006;EC00021;EC00022 |
| Negative | protoporphyrinogen IX mitochondrial transport | 8 | 8 | 0.09615 | 0.1916 | 1.01705 | -1.488 | EC00025;EC0003;EC00024;EC00019;EC00026;EC0006;EC00021;EC00022 |
| Negative | uroporphyrinogen decarboxylase (uroporphyrinogen III) | 8 | 8 | 0.09615 | 0.1916 | 1.01705 | -1.488 | EC00025;EC0003;EC00024;EC00019;EC00026;EC0006;EC00023;EC00021;EC00022 |
| Negative | uroporphyrinogen-III synthase | 8 | 8 | 0.09615 | 0.1916 | 1.01705 | -1.488 | EC00025;EC0003;EC00024;EC00019;EC00026;EC0006;EC00023;EC00021;EC00022 |
| Negative | Nitric Oxide Synthase (NO forming) | 8 | 8 | 0.09615 | 0.1916 | 1.01705 | -1.488 | EC00025;EC0003;EC00024;EC00019;EC00026;EC0006;EC00023;EC00021;EC00022 |
| Negative | hydrogen peroxide transport via diffusion | 8 | 8 | 0.09615 | 0.1916 | 1.01705 | -1.488 | EC00025;EC0003;EC00024;EC00019;EC00026;EC0006;EC00023;EC00021;EC00022 |
| Negative | o2 transport (diffusion) | 8 | 8 | 0.09615 | 0.1916 | 1.01705 | -1.488 | EC00025;EC0003;EC00024;EC00019;EC00026;EC0006;EC00023;EC00021;EC00022 |
| Negative | fatty acid intracellular transport | 8 | 8 | 0.09615 | 0.1916 | 1.01705 | -1.488 | EC00025;EC0003;EC00024;EC00019;EC00026;EC0006;EC00023;EC00021;EC00022 |

| | | | | | | | | |
|---|---|---|---|---|---|---|---|---|
| Negative | dehydroascorbate transport (uniport) | 7 | 7 | 0.08163 | 0.1916 | 1.08815 | -1.522 | EC00025;EC0003;EC00024;EC00019;EC00026;EC0006;EC00021;EC00022 |
| Negative | Glutathione dehydrogenase (dehydroascorbate reductase) | 8 | 8 | 0.09615 | 0.1916 | 1.01705 | -1.548 | EC00025;EC0003;EC00024;EC00019;EC00026;EC0006;EC00023;EC00021;EC00022 |
| Positive | glutamine phosphoribosyldiphosphate amidotransferase | 4 | 4 | 0.07895 | 0.222 | 1.10265 | -1.569 | EC00014;EC00028;EC00029;EC00030;EC0004;EC00016;EC00017;EC00024;EC00027 |
| Positive | phosphoribosylaminoimidazole carboxylase | 4 | 4 | 0.07895 | 0.222 | 1.10265 | -1.569 | EC00014;EC00028;EC00029;EC00030;EC0004;EC00016;EC00017;EC00024;EC00027 |
| Positive | phosphoribosylaminoimidazole synthase | 4 | 4 | 0.07895 | 0.222 | 1.10265 | -1.569 | EC00014;EC00028;EC00029;EC00030;EC0004;EC00016;EC00017;EC00024;EC00027 |
| Positive | phosphoribosylaminoimidazolecarboxamide formyltransferase | 4 | 4 | 0.07895 | 0.222 | 1.10265 | -1.569 | EC00046;EC00047;EC00031;EC00049;EC00014;EC00028;EC00029;EC00030;EC0004;EC00016;EC00017;EC00024;EC00026;EC00027 |
| Positive | phosphoribosylaminoimidazolesuccinocarboxamide synthase | 4 | 4 | 0.07895 | 0.222 | 1.10265 | -1.569 | EC00046;EC00047;EC00031;EC00049;EC00014;EC00028;EC00029;EC00030;EC0004;EC00016;EC00017;EC00024;EC00026;EC00027 |
| Positive | phosphoribosylformylglycinamidine synthase | 4 | 4 | 0.07895 | 0.222 | 1.10265 | -1.569 | EC00046;EC00047;EC00031;EC00049;EC00014;EC00028;EC00029;EC00030;EC0004;EC00016;EC00017;EC00024;EC00026;EC00027 |
| Positive | phosphoribosylglycinamide formyltransferase | 4 | 4 | 0.07895 | 0.222 | 1.10265 | -1.569 | EC00046;EC00047;EC00031;EC00049;EC00014;EC00028;EC00029;EC |

| | | | | | | | | |
|---|---|---|---|---|---|---|---|---|
| | | | | | | | | 00030;EC0004;EC00016;EC00017;EC00024;EC00026;EC00027 |
| Positive | phosphoribosylglycinamide synthase | 4 | 4 | 0.07895 | 0.222 | 1.10265 | -1.569 | EC00046;EC00047;EC00031;EC00049;EC00014;EC00028;EC00029;EC00030;EC0004;EC00016;EC00017;EC00024;EC00026;EC00027 |
| Positive | adenylosuccinate lyase | 4 | 4 | 0.07895 | 0.222 | 1.10265 | -1.569 | EC00046;EC00047;EC00031;EC00049;EC00014;EC00028;EC00029;EC00030;EC0004;EC00016;EC00017;EC00024;EC00026;EC00027 |
| Positive | phosphoribosylpyrophosphate synthetase | 4 | 4 | 0.07895 | 0.222 | 1.10265 | -1.569 | EC00046;EC00047;EC00031;EC00049;EC00014;EC00028;EC00029;EC00030;EC0004;EC00016;EC00017;EC00024;EC00026;EC00027 |
| Negative | 7-alpha,24(S)-Dihydroxycholesterol exchange | 1 | 1 | 0.04167 | 0.1916 | 1.38018 | -1.578 | EC00019 |
| Negative | 7-alpha,25-Dihydroxycholesterol exchange | 1 | 1 | 0.04167 | 0.1916 | 1.38018 | -1.578 | EC00019 |
| Negative | 7-alpha,27-Dihydroxycholesterol exchange | 1 | 1 | 0.04167 | 0.1916 | 1.38018 | -1.578 | EC0004;EC00012 |
| Negative | cholesterol 25-hydroxylase | 1 | 1 | 0.04167 | 0.1916 | 1.38018 | -1.578 | EC0004;EC00012 |
| Negative | cytochrome P450, family 46, subfamily A, polypeptide 1 | 1 | 1 | 0.04167 | 0.1916 | 1.38018 | -1.578 | EC0005 |
| Negative | oxysterol 7-alpha-hydroxylase | 1 | 1 | 0.04167 | 0.1916 | 1.38018 | -1.578 | EC00026;EC0005 |

| | | | | | | | | |
|---|---|---|---|---|---|---|---|---|
| Negative | 24 trihydroxy cholesterol transport | 1 | 1 | 0.04167 | 0.1916 | 1.38018 | -1.578 | EC00025;EC0007;EC0003;EC00024;EC00019;EC00026;EC0006;EC00021;EC00022 |
| Negative | 25 trihydroxy cholesterol transport | 1 | 1 | 0.04167 | 0.1916 | 1.38018 | -1.578 | EC00025;EC0003;EC00024;EC00019;EC00026;EC0006;EC00021;EC00022 |
| Negative | 27 trihydroxy cholesterol transport | 1 | 1 | 0.04167 | 0.1916 | 1.38018 | -1.578 | EC00025;EC0003;EC00024;EC00019;EC00026;EC0006;EC00021;EC00022 |
| Negative | oxysterol 7alpha-hydroxylase | 1 | 1 | 0.04167 | 0.1916 | 1.38018 | -1.578 | EC00025;EC0003;EC00024;EC00019;EC00026;EC0006;EC00021;EC00022 |
| Negative | NADP transporter, peroxisome | 2 | 2 | 0.02326 | 0.1916 | 1.63339 | -1.58 | EC00025;EC00011;EC00013;EC0003;EC00024;EC00019;EC00026;EC0005;EC00015 |
| Negative | NADPH transporter, peroxisome | 2 | 2 | 0.02326 | 0.1916 | 1.63339 | -1.58 | EC00025;EC0003;EC00024 |
| Negative | Reduced glutathione exchange | 6 | 6 | 0.04 | 0.1916 | 1.39794 | -1.776 | EC00019 |
| Negative | cysteinesulfinic acid oxidase | 5 | 5 | 0.02128 | 0.1916 | 1.67203 | -1.784 | EC00014 |
| Negative | Hypotaurine oxidase | 5 | 5 | 0.02128 | 0.1916 | 1.67203 | -1.784 | EC00014 |
| Negative | Arachidic acid exchange | 5 | 5 | 0.02128 | 0.1916 | 1.67203 | -1.784 | EC0007;EC00024;EC00019;EC00026;EC0005 |
| Negative | O2 transport, endoplasmic reticulum | 5 | 5 | 0.02128 | 0.1916 | 1.67203 | -1.786 | EC00014 |

| | | | | | | | | |
|---|---|---|---|---|---|---|---|---|
| Negative | pyridoxal transport via diffusion | 4 | 4 | 0.025 | 0.1916 | 1.60206 | - 1.851 | EC00025;EC0003 |
| Negative | pyridoxal kinase | 4 | 4 | 0.025 | 0.1916 | 1.60206 | - 1.851 | EC00025;EC0003 |
| Negative | L-Ascorbate exchange | 4 | 4 | 0.025 | 0.1916 | 1.60206 | - 1.851 | EC00024 |
| Negative | Glutathione:cystine oxidoreductase | 4 | 4 | 0.025 | 0.1916 | 1.60206 | - 1.851 | EC00024 |
| Negative | Vitamin D3 uptake | 3 | 3 | 0.025 | 0.1916 | 1.60206 | - 1.852 | EC00025;EC0003;EC00024;EC00023 |
| Negative | L-carnitine transport out of mitochondria via diffusion | 3 | 3 | 0.025 | 0.1916 | 1.60206 | - 1.852 | EC00024 |
| Negative | methylmalonyl-CoA mutase | 3 | 3 | 0.025 | 0.1916 | 1.60206 | - 1.852 | EC00024 |
| Negative | Propionyl-CoA carboxylase, mitochondrial | 3 | 3 | 0.025 | 0.1916 | 1.60206 | - 1.852 | EC00024 |
| Negative | 3-Hydroxy-L-kynurenine hydrolase | 6 | 6 | 0.04 | 0.1916 | 1.39794 | - 1.853 | EC00024 |
| Negative | 3-hydroxyanthranilate 3,4-dioxygenase | 6 | 6 | 0.04 | 0.1916 | 1.39794 | - 1.853 | EC00024 |
| Negative | 5-hydroxy-L-tryptophan secretion via secretory vesicle (ATP driven) | 6 | 6 | 0.04 | 0.1916 | 1.39794 | - 1.853 | EC00024 |
| Negative | kynurenine 3-monooxygenase | 6 | 6 | 0.04 | 0.1916 | 1.39794 | - 1.853 | EC00018 |
| Negative | L-Tryptophan:oxygen 2,3-oxidoreductase (decyclizing) | 6 | 6 | 0.04 | 0.1916 | 1.39794 | - 1.853 | EC0005 |

| | | | | | | | | |
|---|---|---|---|---|---|---|---|---|
| Negative | N-Formyl-L-kynurenine amidohydrolase | 6 | 6 | 0.04 | 0.1916 | 1.39794 | -1.853 | EC0001;EC0005;EC0002 |
| Negative | nicotinate-nucleotide diphosphorylase (carboxylating) | 6 | 6 | 0.04 | 0.1916 | 1.39794 | -1.853 | EC0001;EC0005 |
| Negative | Quinolinate Synthase (Eukaryotic) | 6 | 6 | 0.04 | 0.1916 | 1.39794 | -1.853 | EC0006;EC00019 |
| Negative | 2-aminomuconate reductase | 6 | 6 | 0.04 | 0.1916 | 1.39794 | -1.853 | EC00019 |
| Negative | aminomuconate-semialdehyde dehydrogenase | 6 | 6 | 0.04 | 0.1916 | 1.39794 | -1.853 | EC00019 |
| Negative | picolinic acid decarboxylase | 6 | 6 | 0.04 | 0.1916 | 1.39794 | -1.853 | EC00019 |
| Negative | ATP transporter, peroxisomal | 4 | 4 | 0.025 | 0.1916 | 1.60206 | -1.854 | EC00025;EC0003;EC00024;EC00019;EC00023 |
| Negative | L-Phenylalanine exchange | 9 | 9 | 0.03448 | 0.1916 | 1.46243 | -1.886 | EC00024 |
| Negative | Iodide:hydrogen-peroxide oxidoreductase 4 | 6 | 6 | 0.02 | 0.1916 | 1.69897 | -2.023 | EC0008 |
| Negative | L-Thyroxine exchange | 6 | 6 | 0.02 | 0.1916 | 1.69897 | -2.023 | EC00014 |
